# G-quadruplex landscape and its regulation revealed by a new antibody capture method

**DOI:** 10.1101/2022.09.03.506459

**Authors:** Subhamoy Datta, Manthan Patel, Chakkarai Sathyaseelan, Chandrama Ghosh, Akanksha Mudgal, Divyesh Patel, Thenmalarchelvi Rathinavelan, Umashankar Singh

**Affiliations:** HoMeCell Lab, Discipline of Biological Engineering, Indian Institute of Technology Gandhinagar, Gandhinagar, Gujarat 382355, India; Centre for Genomics and Child Health, Blizard Institute, Barts and The London School of Medicine and Dentistry, Queen Mary University of London, London E1 2AT, UK; Department of Biotechnology, Indian Institute of Technology Hyderabad, Kandi Campus, Telangana 502285, India; Azrieli Faculty of Medicine, Bar-Ilan University, Henrietta Szold 8A, Safed, 1311502, Israel; Department of Biopharmacy, Medical University of Lublin, Lublin, 20059, Poland; Research Programs Unit, Applied Tumor Genomics Program, Faculty of Medicine, University of Helsinki, Biomedicum, Helsinki 00290, Finland

**Author notes:** Corresponding author: Umashankar Singh.

**Keywords:** DNA G-quadruplexes, G4-ChIP, CGGBP1, CTCF, G/C-skew

## Abstract

Our understanding of DNA G-quadruplexes (G4s) from *in vitro* studies has been complemented by genome-wide G4 landscapes from cultured cells. Conventionally, the formation of G4s is accepted to depend on G-repeats such that they form tetrads. However, genome-wide G4s characterized through high-throughput sequencing suggest that these structures form at a large number of regions with no such canonical G4-forming signatures. Many G4-binding proteins have been described with no evidence for any protein that binds to and stabilizes G4s. It remains unknown what fraction of G4s formed in human cells are protein-bound. The G4-chromatin immunoprecipitation (G4-ChIP) method hitherto employed to describe G4 landscapes preferentially reports G4s that get crosslinked to proteins in their proximity. Our current understanding of the G4 landscape is biased against representation of G4s which escape crosslinking as they are not stabilized by protein-binding and presumably transient. We report a protocol that captures G4s from the cells efficiently without any bias as well as eliminates the detection of G4s formed artifactually on crosslinked sheared chromatin post-fixation. We discover that G4s form sparingly at SINEs. An application of this method shows that depletion of a repeat-binding protein CGGBP1 enhances net G4 capture at CGGBP1-dependent CTCF-binding sites and regions of sharp interstrand G/C-skew transitions. Thus, we present an improved method for G4 landscape determination and by applying it we show that sequence property-specific constraints of the nuclear environment mitigate G4 formation.

## INTRODUCTION

The containment and maintenance of the genomic DNA inside the human cellular nuclei is a tightly regulated process. Higher-order chromatin conformation and chromosomal territories exemplify the manifestation of an orderly containment of the genomic DNA at a broad scale^1,2^. These arrangements selectively allow regulatory interactions between genomic regions separated by long distances or located *in trans*. It is well established that this orderly arrangement of the genome depends on interactions between DNA and chromatin regulatory proteins, such as CTCF^3^ and its co-operators, such as CGGBP1^4–6^. The complexity of such regulatory proteins and their cognate binding sites in the genome has increased in the course of evolution. Essentially, a significant amount of cellular resources are directed to ensure that the DNA does not randomly adopt conformations which would interfere with DNA-protein interactions^7,8^.

The innate nature of cross-strand annealing combined with repetitive sequences in the human genome makes the DNA prone to randomly adopting multiple conformations. This problem is further exacerbated by the sheer size of the human genome, the restricted nuclear volume and the need to reset the genomic DNA arrangement in the nuclei through replication and transcription^9,10^. Obviously, this challenge could not be addressed just by the regulators of higher-order chromatin conformation with their sequence-specific DNA bindings. In agreement with this, the highly conserved histone proteins, related protein machinery dedicated to packaging DNA into chromatin and single-strand DNA-binding proteins exhibit little DNA sequence preferences^11,12^. These mechanisms disallow the existence of nascent DNA, which would be prone to adopting multiple conformations as allowed biochemically and thermodynamically in the nuclear environment. Collectively, these mechanisms constitute constraints of the nuclear environment on the property of genomic DNA to adopt secondary structures.

The DNA in the nucleus is, however, under a state of flux through various processes that involve strand separation and new strand synthesis. In addition, certain functional regions of the genome, such as transcription enhancers and promoter elements, interact facultatively with transcription factors and may evade dense packaging with histone and related proteins in a cell-type and developmental state-specific manner. The DNA in these regions, often associated with G/C richness^13,14^, becomes amenable to adopt secondary structures. One such secondary structure of DNA that has attracted wide interest is the G4s^8^. The G4s form due to a preference for Hoogsteen base pairing over Watson-Crick base pairing in certain sequence contexts and involves four strands of DNA. The G4s could form from multiple parallel or antiparallel conformations of the same or different DNA strands. The variety of G4s discovered so far indicates that the structure formed is the lowest energy state conformation possible in a certain biochemical context, a process solely dependent on the chemical properties of the DNA, its nucleotide sequence, strandedness and the biochemical environment^15–17^. Much of our understanding of the nature of G4s comes from *in vitro* characterization of DNA secondary structures^18,19^. A commonly accepted signature sequence for G4 formation is a tandem repeat of [G_3_-N_(1-7)_] four times on a single strand. However, in a complex mix of DNA, such as the human genome, single units of [G_3_-N_(1-7)_] from different molecules could collaborate and form intermolecular G4s^18,19^. It is obvious that the formation of secondary structures such as G4s in cellular nuclei would be enhanced if the DNA escapes the constraints of the nuclear environment. Determination of the G4 landscape of the human genome has the potential to elucidate their biological significance.

The adoption of secondary structures by the DNA is a hindrance to genomic DNA usage and maintenance. Hairpin loops and G4s formed by tandem repeats cause polymerase stalling and replication errors^20^. Repetitive sequences prone to the formation of secondary quadruplex structures depend on helicases^21^ for efficient replication and repair. These energy-dependent secondary structure resolution mechanisms collaborate with the constraints of the nuclear environment to mitigate secondary structure formation. More recently, regulatory functions have been attributed to quadruplex structures^8,11,22,23^. Studies indicate that the regulation of expression of some genes depends on quadruplex formation in their cis-regulatory regions^24–26^. It remains unclear if the G4s formation in cis-regulatory regions is a consequence of nucleosome depletion and relaxation of constraints of the nuclear environment or if there are dedicated mechanisms to ensure the formation and stabilization of such regulatory quadruplexes. An increasing number of studies have reported an array of G4-binding proteins^27–31^. However, most of these studies do not distinguish between proteins binding to sequences capable of forming G4s or actual G4 structures. The consequences of protein-G4 interactions remain unknown for many such candidate quadruplex binding proteins. There is evidence that many helicases with RRM domains or RGG/UP1 motifs bind to and resolve the G4s^32–34^. Hundreds of other proteins have been identified as associated with G4s with no indications of any common quadruplex-specific binding domain^27^.

While there is overwhelming evidence that G4s affect DNA-protein interactions, thereby affecting the epigenome, there is no evidence for a targeted G4-stabilizing mechanism in human cells. Most of these G4-binding proteins seem to have an affinity for generally GC-rich DNA and nucleosome-depleted regions, suggesting that their binding could exert constraints of the nuclear environment on G4 formation. The G4 landscape determination with knockdown of these G4-binders (and putative G4 regulators) will elucidate the role of these proteins in the quadruplex establishment in cells. A comprehensive determination of the G4 landscape and their regulation by quadruplex-binding proteins shall address the non-canonical G4s as well.

Valuable studies have described the genome-wide G4 landscapes in human cells^15,35,36^. The canonical [G_3_-N_(1-7)_]xn signature is only mildly enriched in these quadruplex pulldown-sequencing assays. Intriguingly, there is an unexplained bias for G/C-rich open chromatin and nucleosome-depleted regions in G4 landscape datasets, and the same G/C-rich regions are predominantly the binding sites for the G4-binding proteins (unpublished observations derived from data published vide Hänsel-Hertsch and coworkers^35^ and Zhang and coworkers^27^). The influence of protein-binding on the capture and detection of G4s in these assays remains unknown. Interestingly, these studies also raise questions about the nature and scope of G4s at locations other than the nucleosome-depleted regions with low or no occupancy of G4-binding proteins.

Our current understanding of the DNA sequence properties associated with G4 formation *in cellulo* depends on the global G4 profiles, primarily generated using the G4-ChIP protocol^36^. Formaldehyde crosslinking, as used in conventional chromatin immunoprecipitation protocols, has been adopted in G4-ChIP as well. Although several G4-binding proteins have been described, no known dedicated G4-binding domains can stabilize the quadruplexes common to them. Formaldehyde crosslinking can efficiently capture only those G4s which are stably bound to proteins. This creates a detection bias against the G4s not bound to proteins. That formaldehyde is a passive DNA denaturing agent^37^o^38^ suggests that protein unbound G4s would be underrepresented in the G4-ChIP. In addition, tetrads formed by non-G nucleotides also exist interspersed between G tetrads, with some even stabilizing the

G4s to varying degrees^39–42^. Currently, we do not know if the large fraction of DNA pulled down by antibodies specific for G4s that does not show any [G_3_-N_(1-7)_]xn signature is an artifact or not. It remains unknown if the G/C-rich and non-G/C-rich G4s are regulated by the same or distinct mechanisms. Nevertheless, it remains established that although G/C richness is not an essential feature of quadruplex-forming regions, the G/C-rich sequences readily form stable quadruplexes *in vitro* and, to a certain extent, *in cellulo*. On the other hand, an overrepresentation of the [G_3_-N_(1-7)_]xn sequences in the G4-ChIP-seq could be due to the rapid adoption of quadruplex structures by [G_3_-N_(1-7)_]xn sequences post-fixation. DNA melted passively by formaldehyde as well as sheared by sonication into single-stranded overhangs would be amenable to forming quadruplexes post-fixation in the lysates or during any step before the anti-G4 antibody is added. The antibodies would capture the *in vitro* formed quadruplexes, which would be falsely represented in the G4-ChIP-seq results as bonafide quadruplexes *in cellulo*. Such a potential misrepresentation of the G4 landscape in the conventional G4-ChIP-seq experiments contaminates our understanding of *in cellulo* G4 landscape with *in vitro* formed quadruplexes.

Thus, multiple interrelated challenges hinder our interpretation of the G4 landscape and its regulation. Does formaldehyde fixation truly capture the *in cellulo* G4s only? Do post-fixation formed quadruplexes *in vitro* get misrepresented as *in cellulo* G4s? Does G4 formation/stabilization depend on some regulatory protein(s)? Are there constraints of the nuclear environment exerted by G4-binding proteins on quadruplex formation? In this work, we shed light on these questions. We have developed a ChIP-sequencing protocol for G4s to stabilize *in cellulo* quadruplexes and minimize any quadruplexes formed post-fixation *in vitro*. We use this protocol with a control DNA, having the [G_3_-N_(1-7)_]xn sequence signature, to establish the extent of constraints of the nuclear environment on quadruplex formation by [G_3_-N_(1-7)_]xn sequences *in cellulo* quantitatively. We use this protocol to describe the most native *in cellulo* G4 landscape and demonstrate that G4s form abundantly at satellite repeats while short-interspersed nuclear elements (SINEs) resist G4 formation.

CGGBP1 is a nuclear protein with a binding preference for G/C-rich repetitive DNA and interspersed repeats, including Alu-SINEs^43–45^. CGGBP1 regulates cytosine methylation at DNA sequences with interstrand G/C-skew^45^, an implied feature of the [G_3_-N_(1-7)_]xn signature attributed to G4 formation. We apply our protocol in cells depleted for CGGBP1 and find that CGGBP1 exerts constraints of the nuclear environment on G4 flux selectively. While CGGBP1 depletion leaves the low G4 prevalence at SINEs unchanged, it causes a net enhancement of G4 formation at regions with G/C-skew, which correlates with enhanced occupancy of CTCF at CGGBP1-regulated CTCF-binding sites. Our results suggest that the genome is under a constant widespread G4-flux, and the prevalence of G4 is determined by an interplay between the local sequence properties and their protein-binding. The new G4 capture method we present thus reveals a more widespread nature of the G4 landscape and its regulation.

## RESULTS

### 1. G4s captured by G4-ChIP also form spontaneously on genomic DNA in vitro

The data derived from G4-ChIP are expected to represent an *in cellulo* G4 landscape. These *in cellulo* G4 landscapes are expected to be different from the G4 profile of purified genomic DNA where there are no constraints of the nuclear environment. We needed to compare the G4 landscapes derived from G4-ChIP with those derived entirely from purified genomic DNA *in vitro*. So we first performed *in vitro* G4 DNA immunoprecipitations (DNA-IPs) on purified genomic DNA from HEK293T in two different ways using the 1H6 antibody. In one of the DNA-IPs, called G4 DNA-IP^*nat*^, the double-stranded genomic DNA stored under naturing conditions for seven days was incubated with the antibody. In a second DNA-IP, called (G4 DNA-IP^*denat*^), the same DNA was heat-denatured and snap-chilled just before the antibody incubation step. The quadruplexes expected to be pulled down in G4 DNA-IP^*nat*^ were stable G4s that would form spontaneously in largely double-stranded genomic DNA. In contrast, the G4s expected in G4 DNA-IP^*denat*^ would be those which form rapidly only when the DNA is rendered single-stranded (after heat denaturation and snap-chilling). In both cases, the binding of antibodies would stabilize the G4s for capture and characterization through sequencing. Both the DNA-IPs and their common input were sequenced (Table S1) and compared.

Compared to G4 DNA-IP^*nat*^, the G4 DNA-IP^*denat*^ showed a higher correlation with the input, suggesting that when the constraints of double-strandedness are overcome, G4 formation on the genomic DNA is rapid and widespread genome-wide *in vitro* (Fig S1). This inference was supported by an analysis of how the G4 enrichments in G4 DNA-IP^*denat*^ and G4 DNA-IP^*nat*^ compared with each other. Using the same conditions of peak-calling, while 15709 peaks were discovered in G4 DNA-IP^*denat*^, only 226 peaks could be called in G4 DNA-IP^*nat*^ (Fig 1A and Table S2). The genome-wide signals from the two DNA-IPs were plotted reciprocally in their peaks (Fig S2). Only a small fraction of the 15709 G4 DNA-IP^*denat*^ peaks were rich in signals from G4 DNA-IP^*nat*^ (Fig S2A). On the contrary, nearly all of the 226 G4 DNA-IP^*nat*^ peaks were rich in G4 DNA-IP^*denat*^ signals (Fig S2B). Strikingly, 92% of the G4 DNA-IP^*nat*^ peaks overlapped with just 1% of the G4 DNA-IP^*denat*^ peaks (Fig S2C). An overwhelming 99% of the G4 DNA-IP^*denat*^ peaks found no overlap with the G4 DNA-IP^*nat*^ peaks (Fig S2D). This suggested that a small fraction of the rapidly forming G4s in G4 DNA-IP^*denat*^ are stable quadruplexes also present in G4 DNA-IP^*nat*^. These results also showed that the immense G4-forming potential of human genomic DNA is restrained by its double strandedness at most of the regions. At the same time, at some regions, the G4s prevail over the double-strandedness and form spontaneously.

**Fig 1:**
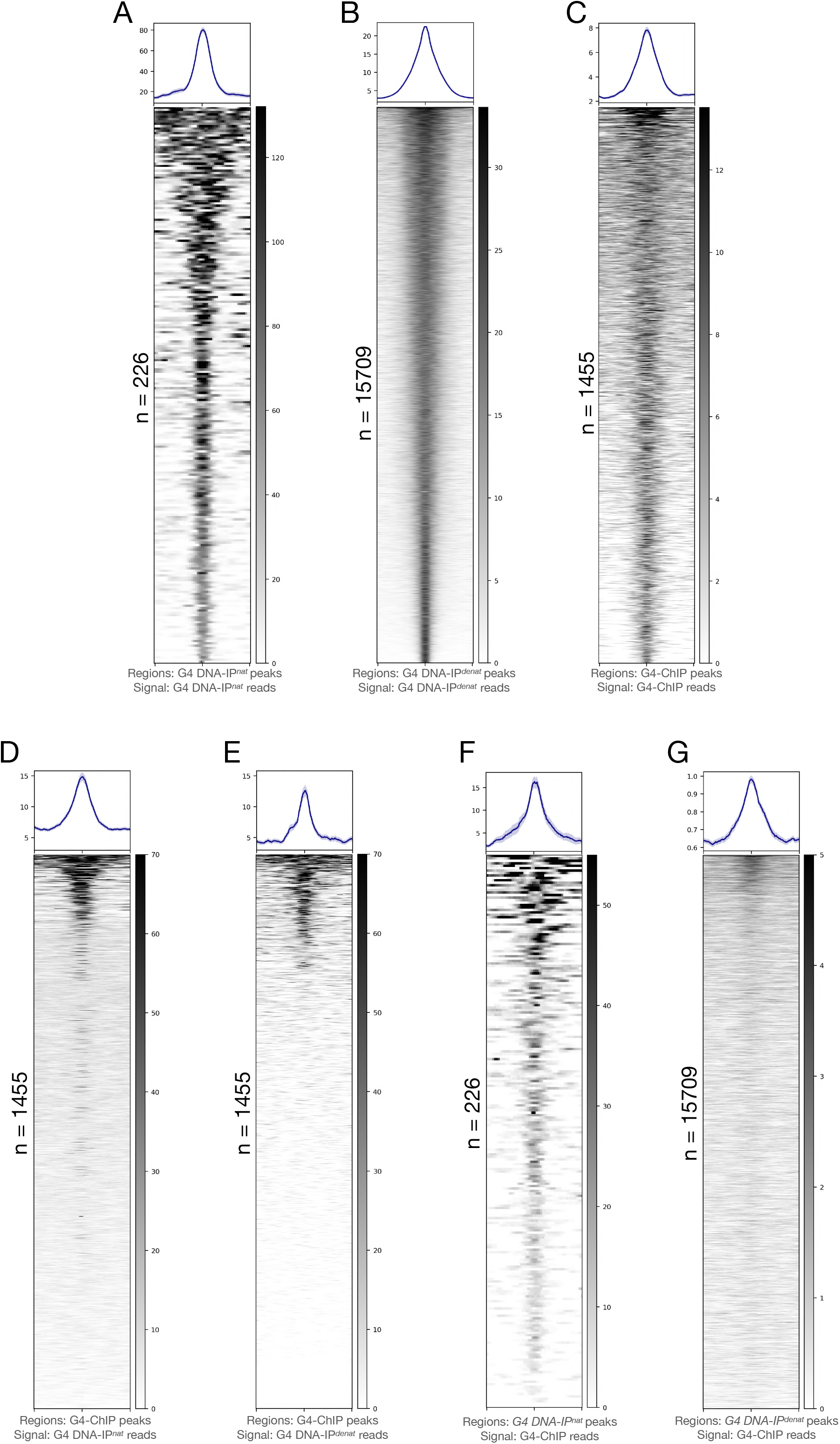
G4 potential of human genome characterized through G4 immunoprecipitation *in vitro* and its comparison with G4-ChIP output: A-C: Peaks from G4 DNA-IP^*nat*^ (A), G4 DNA-IP^*denat*^ (B) and G4-ChIP (C) show specificities of G4 DNA pulldown. The G4 pulldown pattern gives rise to far fewer peaks in G4 DNA-IP^*nat*^ (A) on one extreme and a large number of peaks in G4 DNA-IP^*denat*^ (B) on the other. G4-ChIP peak count is higher but with a weaker mean signal than either G4 DNA-IP^*nat*^ or G4 DNA-IP^*denat*^. D and E: A plotting on G4-ChIP peaks of signals from G4 DNA-IP^*nat*^ (D) or G4 DNA-IP^*denat*^ (F) shows that visibly more G4-ChIP peaks contain G4 DNA-IP^*nat*^ signals than G4 DNA-IP^*denat*^ signals. F and G: Most G4 DNA-IP^*nat*^ peaks (F) contained a strong G4-ChIP signal, whereas, on G4 DNA-IP^*denat*^ peaks (G), the G4-ChIP strong signals were restricted to a very small fraction of the peaks. The peak counts are mentioned beside the heatmaps. Signals are plotted as mean±SE on the Y axes, and bed coordinates are arranged in decreasing order of the signal magnitude. All signals are plotted in regions 5 kb±peak centers.

Stable G4s form at the canonical [G_3_-N_(1-7)_]xn signature sequences. Also, it is known that the 1H6 antibody can capture secondary DNA structures without restricting to the canonical [G_3_-N_(1-7)_]xn signature sequences, especially upon denaturation^42^. Quality-filtered reads from the two assays were searched for the presence of G4-forming [G_3_-N_(1-7)_]xn sequences using *allquads*^19^. We observed that the [G_3_-N_(1-7)_]xn sequences were enriched in the G4 DNA-IP^*nat*^ sample, whereas the G4 DNA-IP^*denat*^ sample showed such sequences only at an expected level as in the input (Table S3). The spontaneity of the formation of G4s and their stability have been shown to depend on the GC-content^46^. As expected, we observed that the GGG/CCC contents of G4 DNA-IP^*nat*^ peaks were significantly much lower compared to G4 DNA-IP^*denat*^ (Fig S3). Thus, a majority of the G4s identified in G4 DNA-IP^*denat*^ seemed to be rapidly forming at G/C-rich DNA, where the energetic constraints of double-stranded DNA are overcome by heat denaturation. The structural turnover of the chromatin in a cell would allow abundant possibilities for such G4s to form.

We next asked how these two types of G4s are captured by the conventional G4-ChIP^36^. G4-ChIP was performed in HEK293T cells using a previously described protocol^36^ (except that we used the 1H6 antibody) and compared its peaks and genome-wide signal with those of G4 DNA-IP^*denat*^ and G4 DNA-IP^*nat*^. Although the signals from the three experiments were highly uncorrelated with each other, the G4-ChIP and G4 DNA-IP^*nat*^ signals clustered together differently from G4 DNA-IP^*denat*^ (Fig S4A). This weak similarity between G4-ChIP and G4 DNA-IP^*nat*^ was relevant as the 1455 G4-ChIP peaks contained strong signals mostly from G4 DNA-IP^*nat*^ (Fig 1D) and only weakly from G4 DNA-IP^*denat*^ (Fig 1E and S4B). The G4-ChIP peaks overlapped with over 70% of the G4 DNA-IP^*nat*^ peaks and less than 20% of the G4 DNA-IP^*denat*^ peaks. Reciprocally, the G4 DNA-IP^*nat*^ peaks contained a stronger and higher abundance of G4-ChIP signals (Fig 1F) as opposed to the G4 DNA-IP^*denat*^ peaks (Fig 1G). As expected, an allquads analysis showed that the G4-ChIP sample was also rich in [G_3_-N_(1-7)_]xn sequences compared to the respective input (Table S4). A similar enrichment of [G_3_-N_(1-7)_]xn sequences was found in independently reported G4-ChIP data using BG4 antibody in HaCaT cells and normal human epidermal keratinocytes (NHEK) (Table S5). Recapitulating the similarities between G4-ChIP peaks and G4 DNA-IP^*nat*^ peaks, the two peak sets also showed a similar abundance of relatively GGG/CCC-poor sequences (Fig S3).

We could conclude from these findings that some regions of the genome, especially those with the [G_3_-N_(1-7)_]xn signature sequences and low GGG/CCC levels, are captured similarly *in vitro* (G4 DNA-IP^*nat*^) and from the chromatin using the G4-ChIP protocol. Although the G4-ChIP predominantly captured these spontaneously-forming G4s, it had a much lower representation of the G4s formed at GGG/CCC-rich regions and sequences poor in [G_3_-N_(1-7)_]xn signature. Although we found the latter type of G4s as the most prevalent in G4 DNA-IP^*denat*^, the reasons for their paucity in the G4-ChIP data, and the possibility of their existence and relevance in the chromatin, remained unaddressed by using the G4-ChIP protocol.

To investigate this further, we analyzed how various steps of the G4-ChIP protocol could affect the capture and enrichment of G4s. G4-ChIP relies on a protocol originally invented to capture DNA-protein interactions. In this protocol, the capture of G4s is expected to be favoured by their stability, crosslinking to a protein by formaldehyde and no steric hindrance to antibody-G4 binding in the downstream steps. There is no evidence for G4 stabilization by G4-binding proteins. The similarity between the G4 profiles of G4 DNA-IP^*nat*^ (derived from protein-unbound DNA) and G4-ChIP suggests that such spontaneously forming G4s could be captured similarly from the DNA or chromatin after fixation and shearing by sonication, as the protocols of G4 DNA-IP^*nat*^, as well as of G4-ChIP, allow stabilization of G4 by their binding to the antibody during the incubation steps. Thus, the possibility of the formation of G4s *in vitro* on the fixed-sheared chromatin and their enrichment during G4-ChIP can not be ruled out. These G4s formed on fixed-sheared chromatin would not represent the G4 landscape in the cells and yet explain the similarity we observed between the G4 DNA-IP^*nat*^ and G4-ChIP datasets. It thus becomes impossible to segregate chromatin-derived G4s from possible *in* vitro-derived G4s in the G4-ChIP data. These results also indicated other possibilities: if the single-stranded DNA could rapidly fold into G4s *in vitro* (G4 DNA-IP^*denat*^), could this also happen in the chromatin? Do such G4s form readily on the DNA in chromatin and are they physiologically relevant? These questions could be answered only if the G4s were captured solely from the chromatin without any possibility of contamination from G4s formed after fixation and shearing.

To clearly establish the G4 landscape under the constraints of the nuclear environment, it was important to first generate the quadruplex landscape in cells with a minimum *in vitro* artifacts. We worked on eliminating the possibility that some sequences, which do not exist as quadruplexes in live cells, adopt quadruplex structures during various stages of the G4-ChIP protocol post-fixation.

### 2. A method for capture of G4s genome-wide without in vitro interference

The G4-ChIP protocol relies on the fixation of G4-protein complexes by formaldehyde followed by lysis of the cellular nuclei, sonication and target precipitation using an antibody of interest. The capture of the G4s by this method depends on (i) an assumed interaction of the quadruplexes with proteins required for formaldehyde fixation and (ii) that the binding of the anti-G4 antibody is not sterically hindered by proteins covalently fixed onto the quadruplexes. Together these factors are expected to create false negatives in the G4 landscape identified by G4-ChIP (Fig 2A). Additionally, the melting effect of formaldehyde on nascent DNA segments^37,38^ of the sonicated chromatin creates single-stranded DNA capable of forming false positive G4s. Moreover, G4s seem to form rapidly and spontaneously without any need for DNA denaturation. These latter types of G4s can create false positives in the G4-ChIP data. To overcome these limitations, we developed a variation of the G4-ChIP to minimize these false positives and false negatives.

**Fig 2:**
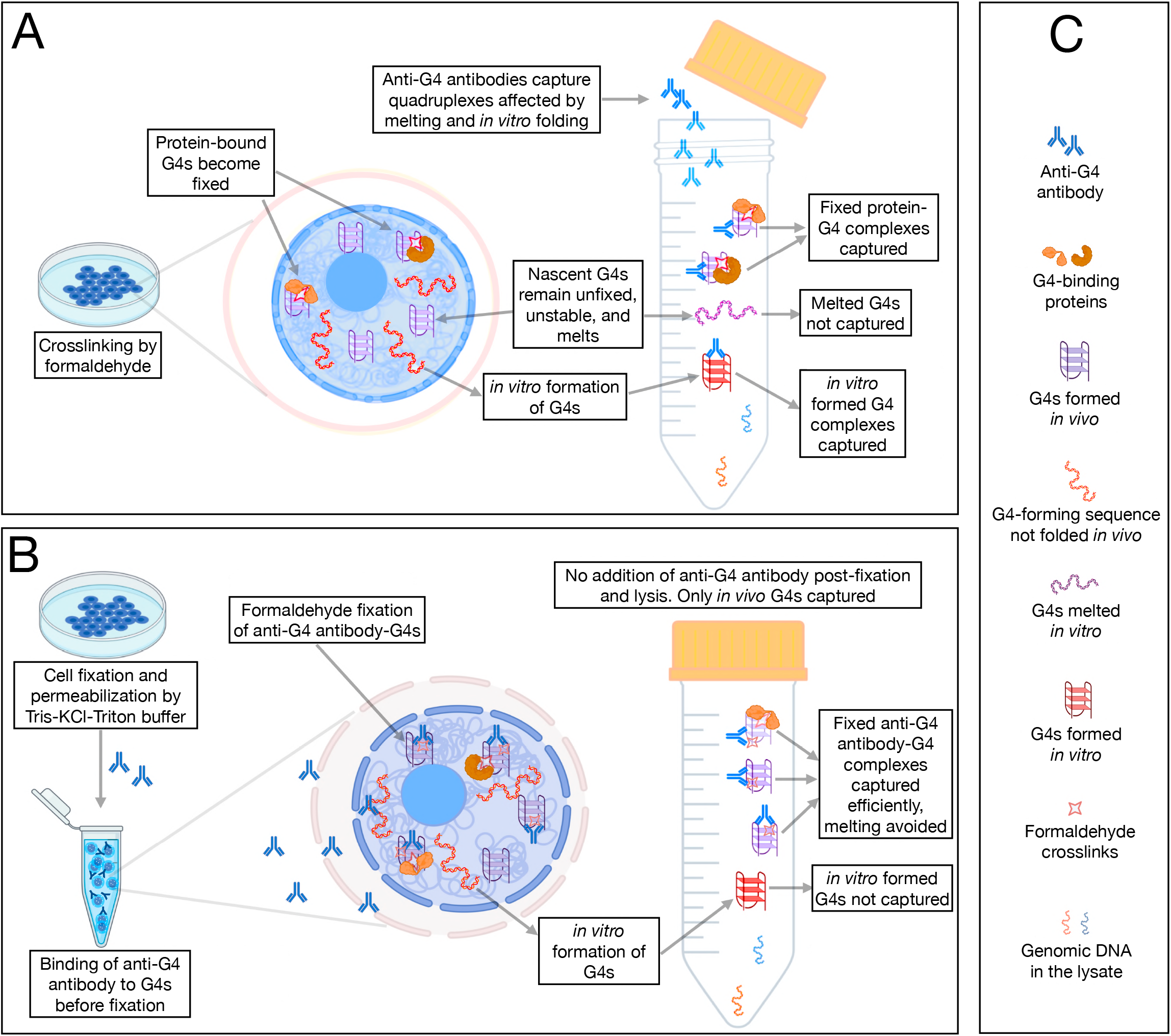
A comparative analysis of G4 landscape in cultured cells using *AbC* G4-ChIP: A: A schematic of the conventional G4-ChIP highlighting the possibilities of false positives and negatives. B: A schematic of the *AbC* G4-ChIP protocol highlighting the key differences from the conventional G4-ChIP and minimization of false positives and false negatives. C: Character legend for A and B.

HEK293T cells were pretreated with formaldehyde briefly, followed by incubation with a buffer containing Triton-X 100 for permeabilization and KCl for maintaining/stabilizing the G4s in their native states (Fig 2B and C). These permeabilized cells were bathed with 1H6 antibody for two hours at room temperature. The antibody-bound G4s were fixed with formaldehyde ensuring that the quadruplexes were captured as antibody-G4 complexes, thereby minimizing false negatives. Also, no further addition of the antibody during the rest of the protocol ensured that any quadruplexes formed post-fixation were not captured. We call this new protocol Antibody Capture G4-ChIP (*AbC* G4-ChIP, detailed protocol is mentioned in methods). The DNA captured using *AbC* G4-ChIP was expected to enrich only the nuclear G4s. Due to the stabilizing effect of antibody-binding on G4s that would form transiently, the *AbC* G4-ChIP protocol allowed the capture of G4s as they formed in the nuclei, thereby generating a wider repertoire of G4s.

This AbC G4-ChIP sample (called CT^*nuc*^) was sequenced (Table S6) and compared against the *in vitro* G4 DNA-IPs and G4-ChIP data (Table S1). The G4-ChIP and G4 DNA-IP^*nat*^ coclustered with each other, whereas the CT^*nuc*^ formed an independent cluster (Fig S4A), suggesting that the *AbC* G4-ChIP returns a G4 landscape substantially different from either of the *in vitro* G4-DNA-IPs (heat-denatured or not) and the G4-ChIP (Fig S4B). To further understand how CT^*nuc*^ was related to G4-ChIP and *in vitro* G4 DNA-IPs, we compared the signals of these datasets on their peaks reciprocally (Table S2). The CT^*nuc*^ signals were distinctly observed at the G4 DNA-IP^*nat*^ and G4 DNA-IP^*denat*^ peaks (Fig S5A and B, respectively). Overlapping subsets of CT^*nuc*^ peaks were rich in signals from G4 DNA-IP^*nat*^ as well as G4 DNA-IP^*denat*^ (Fig S5C and D, respectively). While CT^*nuc*^ signals were abundant at G4-ChIP peaks, the G4-ChIP signals were poor at CT^*nuc*^ peaks (Fig S6A and B, respectively). Very few CT^*nuc*^ peaks showed intersects with G4-DNA-IP^*nat*^ peaks (Fig S7A), whereas nearly half of CT^*nuc*^ peaks showed overlaps with the G4 DNA-IP^*denat*^ peaks (Fig S7B); overlaps between peaks of G4-ChIP and CT^*nuc*^ were also relatively weaker (Fig S7C and S4B). The peaks were also compared by Jaccard indices (Fig S8). These indices showed that the small number of G4 DNA-IP^*nat*^ peaks or the far numerous G4-DNA-IP^*denat*^ peaks had similar overlaps with the CT^*nuc*^ peaks (Table S2). The G4-ChIP peaks, which overlapped strongly with *in vitro* G4-DNA-IP^*nat*^ peaks, showed a very weak intersection with CT^*nuc*^ peaks. We also analyzed the presence of known G4-forming sequence properties in these peaks. The [G_3_-N_(1-7)_]xn content was slightly different between CT^*nuc*^ and G4-ChIP (Table S4). However, the GGG/CCC content between the G4-ChIP and CT^*nuc*^ peaks was strikingly different (p-values <0.0001) (Fig S3). Unlike G4-ChIP, the CT^*nuc*^ peaks were almost devoid of the low GGG/CCC content reads, a feature associated with the spontaneously forming G4s in the G4-DNA-IP^*nat*^.

These properties of the CT^*nuc*^ sequence data showed that the *AbC* G4-ChIP protocol captures G4s of both kinds as represented by G4 DNA-IP^*nat*^ and *in vitro* G4-DNA-IP^*denat*^. In AbC G4-ChIP however, the spontaneously-forming G4s at GGG/CCC-poor regions account for a far lower fraction of the total G4s as compared to that in G4-ChIP. In contrast, G4-ChIP captures almost exclusively the G4s represented by G4 DNA-IP^*nat*^ and fails to capture the ones represented by G4 DNA-IP^*denat*^. Overall, the overwhelming similarities between G4-ChIP and *in vitro* G4-DNA-IP^*nat*^ show that G4-ChIP captures spontaneously forming G4s at low GGG/CCC sequences, including the G4s that form post-fixation. A comparison with CT^*nuc*^ showed that these low GGG/CCC quadruplexes are not available for antibody binding in intact cells. These are likely *in vitro* artifacts captured in G4-ChIP but eliminated in CT^*nuc*^.

A measure of this artifact was derived by analyzing G4 signals captured in these assays at randomly drawn genomic regions (details in the methods). Unlike the regions captured in these assays using the 1H6 antibody (where G4 signals were expected), these randomly selected regions were not expected to be rich in G4 signals. As expected, the G4 signals in these random genomic regions captured in CT^*nuc*^ were very low; across all the samples, about 78-83% of the regions contained just 0.001% of all the G4 signals captured (Fig S9A). However, strikingly, the largest deviation from low or no signals in these random regions was observed in G4-ChIP and G4 DNA-IP^*nat*^ (Fig S9A). When normalized for sequencing depth, the G4 signals in these randomly drawn regions were highest in G4 DNA-IP^*nat*^ followed by G4 DNA-IP^*denat*^, G4-ChIP and CT^*nuc*^ (Fig S9B to S9E, respectively) in decreasing order. In previously published G4-ChIP datasets, also we observed a higher signal in randomly drawn regions than in CT^*nuc*^ (Fig S10). The overall low levels of signals at randomly drawn regions showed that the 1H6 antibody does not capture DNA randomly, and the pull-down has specificity. The remarkable difference in the G4 signals between G4-ChIP and CT^*nuc*^ reinforced the fact that (i) G4s form spontaneously at many regions in the genome that can be drawn in any randomly chosen regions of the genome, (ii) the chromatin environment prevents the formation of these G4s, and (iii) between G4-ChIP and CT^*nuc*^, the latter is better at not capturing G4s formed at these randomly drawn regions thereby delivering a cleaner G4 landscape from the chromatin. Thus, the formation of G4s at random regions gives rise to artifacts similar to the G4 DNA-IP^*nat*^ and G4-ChIP. The formation of G4s at random regions in G4-ChIP likely happens during the antibody incubation step.

### 3. An application of AbC G4-ChIP identifies CGGBP1 as an agent of constraints of the nuclear environment on G4 formation

To test if *AbC* G4-ChIP actually eliminates false positive quadruplexes, we applied additional controls to differentiate the *in cellulo* G4s from the *in vitro* formed G4s confidently. We generated a Control DNA that contained three repeats of the unit canonical [*G_3_-N_5_-G_3_-N_5_-G_3_-N_5_-G_3_*] signature sequence (Fig 3A) and cloned it in the pGEM-T Easy vector. The quadruplex-forming property of a unit [*G_3_-N_5_-G_3_-N_5_-G_3_-N_5_-G_3_*] signature sequence (labeled DG in Fig S11) was established using CD spectroscopy under different concentrations of KCl. DG showed a positive peak around 210 nm in the CD spectra (Fig S11A). The negative peak around 240 nm and the positive peaks at 260 nm and 290 nm further confirmed the formation of hybrid G4s^47,48^. This G4-forming property of DG was lost when it was paired with its reverse complementary oligonucleotide in a Watson-Crick pairing (labeled WC in Fig S11). The hypochromic thermal melting curve (at 260 nm) observed for DG was independent of the KCl concentration, indicating the formation of G4s^49,50^ (Fig S11B). Electrophoretic mobility shift assays further indicated that the hybrid G4 conformations preferred by DG were intramolecular (Lanes 2-6 in Fig S11C) as it moved faster than the d(T)_27_ oligonucleotide (Lane 1 in Fig S11C). We could conclude from these CD and EMSA experiments that the Control DNA could form an intramolecular hybrid G4 which can have any of the quadruplex folds depicted in Fig S11D. Subsequently, this Control DNA was used in the *AbC* G4-ChIP protocol as a spike-in to distinguish G4s formed only under constraints of the nuclear environment from those formed only *in vitro*.

**Fig 3:**
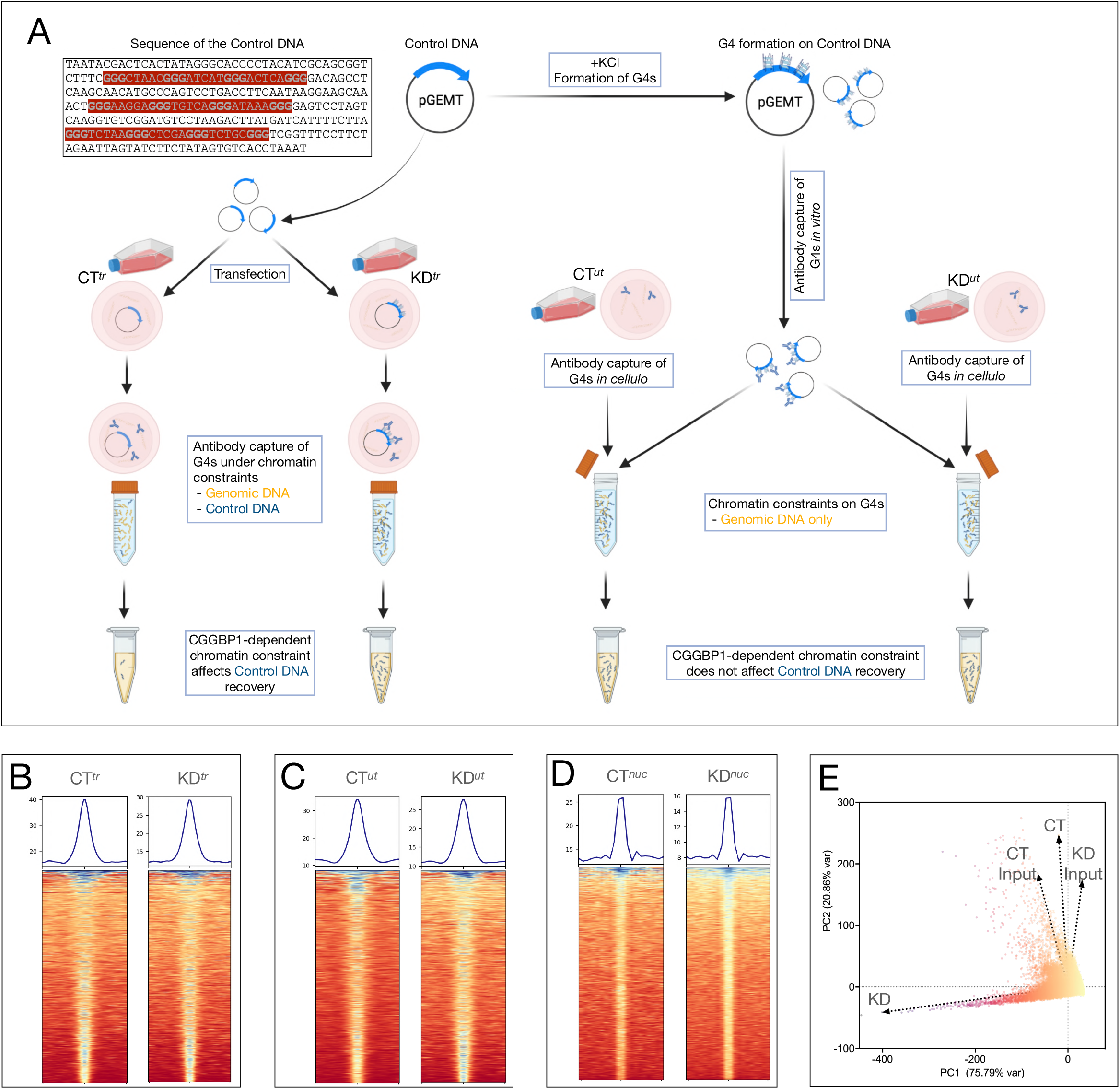
Validation of the *AbC* G4-ChIP protocol using a control DNA and its application to a CGGBP1 knockdown system: A: A schematic of the control DNA (sequence with G-triplets highlighted in bold) cloned in pGEM-T Easy vector and its usage in *AbC* G4-ChIP. The left flank shows the protocol in which the control DNA (along with the carrier DNA) is transfected (samples CT^*tr*^ and KD^tr^). The right flank shows the samples CT^*ut*^ and KD^*ut*^ in which the preformed G4s on the control DNA are added to the lysates. As indicated, the effects of the intranuclear environments of live cells on G4s would be seen only in CT^*tr*^ and KD^*tr*^ but not in CT^*ut*^ and KD^*ut*^. B-D: Heatmaps and signal plots showing G4 enrichment at peak regions as compared to the neighbouring 5 kb flanks. The signals were consistently weaker at the peaks in KD samples as compared to the respective CT samples. E: A PCA plot of the combined signals of the three CT and KD experiments shows that CGGBP1 depletion remarkably alters the G4 landscape. The values used for PCA loading were derived from the summation of signals in synchronous 0.2 kb genomic bins after the removal of bins with signals < 3 and > 100.

This Control DNA could be used in two different ways: (i) added to the sonicated lysate to represent the *in vitro* false-positive quadruplexes formed without any constraints of the nuclear environment, or (ii) transfected into the cells so that the quadruplexes formation on the same sequence would be subjected to constraints of the nuclear environment (Fig 3A). The differential capture of the Control DNA in the two samples would reveal the extent of constraints of the nuclear environment on G4s formation on sequences with canonical G4s forming signatures. We applied the *AbC* G4-ChIP with the Control DNA in these two different ways for functional studies on a candidate G4-regulatory protein CGGBP1.

CGGBP1 is a GC-rich DNA triplet repeat-binding protein. It exhibits a heightened occupancy at putatively G4-forming GGG triplets in growth-stimulated cells (as described elsewhere^43^ and Fig S12). We hypothesized that if the possibility of constraints of the nuclear environment by CGGBP1 is true, then the depletion of CGGBP1 would affect the G4 landscape. *AbC* G4-ChIP with the Control DNA was employed to determine the G4 landscape in HEK293T cells with or without CGGBP1.

CGGBP1 was knocked down in HEK293T cells (Fig S13) as described elsewhere^5^. The non-targeting shRNA sample was called CT, and the CGGBP1-depleted sample was called KD. The CT and KD samples were transfected with the Control DNA and subjected to *AbC* G4-ChIP (Fig 3A). To rule out any transfection efficiency bias between CT and KD samples due to secondary structure formation in the Control DNA, we co-transfected an equimolar amount of a carrier DNA, with no detectable G4-forming sequence (all tetrad-forming GGG sequences replaced with ATG; *allquads* output, not shown), also cloned in pGEM-T Easy vector. The recovery of the Control and the carrier DNA from the inputs were verified by quantitative PCR (Fig S14). There was no significant difference in the transfection efficiency between the CT and KD. These samples were called CT^*tr*^ and KD^*tr*^. In these samples, the Control DNA was expected to be captured with 1H6 antibody only if it formed quadruplexes in the nuclei, presumably under constraints of the nuclear environment. In parallel, CT and KD were also processed for *AbC* G4-ChIP without any transfections of either Control or carrier DNA (CT^*ut*^ and KD^*ut*^, respectively). These samples were processed till nuclear lysis without any transfection. Then, an equimolar mix of Control and carrier DNA was allowed to fold *in vitro* in the presence of KCl, incubated with 1H6 antibody, formaldehyde-fixed and added to the lysates of CT^*ut*^ and KD^*ut*^ (Fig 3A). The CT^*tr*^, KD^*tr*^, CT^*ut*^ and KD^*ut*^ samples were processed identically for the rest of the *AbC* G4-ChIP protocol and sequenced. In addition, we also sequenced the *AbC* G4-ChIP pulldown from a KD sample without any transfection or *in vitro* addition of Control or carrier DNA to capture only the nuclear G4s. This sample (called KD^*nuc*^) was compared against the CT^*nuc*^ sample described above (Fig 2 and table S7). Thus, *AbC* G4-ChIP was performed on three non-identical replicates of CT (CT^*nuc*^, CT^*tr*^ and CT^*ut*^) and KD (KD^*nuc*^, KD^*tr*^ and KD^*ut*^).

First, we analyzed the Control DNA pulldown in CT^*tr*^, KD^*tr*^, CT^*ut*^ and KD^*ut*^ datasets. The coverage of sequencing reads on the Control DNA in the input samples were low, with an enrichment in the *AbC* G4-ChIP samples. The number of reads mapping to the Control DNA increased upon CGGBP1 knockdown in KD^*tr*^ compared to CT^*tr*^ (Table S8). The read counts mapping to the Control DNA in CT^*ut*^ and KD^*ut*^ remained unaffected. In CT^*tr*^, the Control DNA was captured as only six reads, which increased to 390 reads in KD^*tr*^. Linked to the Control DNA sequences, the vector backbone also showed similar variations (not shown). It could be concluded from these results that upon transfection, quadruplex formation on the Control DNA was under an inhibitory influence of CGGBP1. Depletion of CGGBP1 relieved this inhibition on the Control DNA, including the linked vector bac kbone. The capture of G4s similarly between CT^*ut*^ and KD^*ut*^ samples and very strongly only in KD^*tr*^ also established that the *AbC* G4-ChIP captures the G4s efficiently and specifically in a quantitatively verifiable manner. Thus, the differences in G4s prevalence on the Control DNA between CT^*tr*^ and KD^*tr*^ could represent the influence of CGGBP1 on G4 formation throughout the genome in the constraints of the nuclear environment. We analyzed the genome-wide G4 landscape captured in the three CT and KD samples and asked if, just like on the Control DNA, the constraints of the nuclear environment exerted by CGGBP1 are also observed on the genomic sequences.

G4 enrichment in CT (CT^*nuc*^, CT^*tr*^, CT^*ut*^) and KD (KD^*nuc*^, KD^*tr*^, KD^*ut*^) was confirmed using peak calling and validated by calculating the read density within the peaks compared to the flanking regions (Fig 3B-D). The three samples also correlated well, such that centrally enriched G4 signals were observed for each dataset pair (Fig S15). We observed a generally slightly weaker enrichment at the peak centers in KD samples as compared to the respective CT samples (Fig 3B-D). This lower concentration of G4s at the peak regions suggested that interspersed *de novo* quadruplexes were formed and captured at non-peak regions or more peaks upon CGGBP1 depletion (number of peaks indicated in Fig 3, B-D). A principal component analysis of the pooled CT and KD datasets and the respective inputs showed that CGGBP1 depletion affected the G4 landscape drastically and accounted for the largest variance (Fig 3E).

To compare the G4 profiles of CT and KD, we first calculated signals in 0.2 kb bins genomewide, selected bins with ≥ 3 (removal of no signal biases) and ≤100 reads (removal of PCR artifacts) in all the samples separately. These G4 signals were then corrected for regional representational biases through a normalization based on signals from the same regions in the respective inputs. Paired t-tests were performed independently for each pair of 873929 genomic bins from CT and KD (n = 3 pairs; CT^*nuc*^-KD^*nuc*^, CT^*tr*^-KD^*tr*^ and CT^*ut*^-KD^*ut*^). We discovered 12319 differentially captured regions (DCRs) with a significant difference in G4s between CT and KD samples (p-value <0.01, Q:1%: two-stage linear step-up procedure of Benjamini, Krieger and Yekutieli; pairwise p and Q values for each bin in Table S9). A correlation analysis between pooled G4 signals in CT and KD (Fig 4A) showed two major categories of 0.2 kb bins; most bins showed no difference between CT and KD (non-DCRs), and the DCRs with a conspicuous increase in the signal in KD over CT (Fig 4A and fig S16). Signals from the three pairs of CT and KD in the flanks of these DCRs showed that they were rich in G4s with a net increase in G4 signal upon CGGBP1 depletion (Fig 4B and C). These were apparently the regions where CGGBP1 exerted constraints of the nuclear environment on G4 formation, and the depletion of CGGBP1 led to a net increase in G4 signals. Conspicuously, there were a much lower number of regions with a decrease in G4 formation in KD as compared to CT (Fig 4A and fig S16). We further analyzed the properties of these differentially captured regions (DCRs) where G4-formation is regulated by CGGBP1.

**Fig 4:**
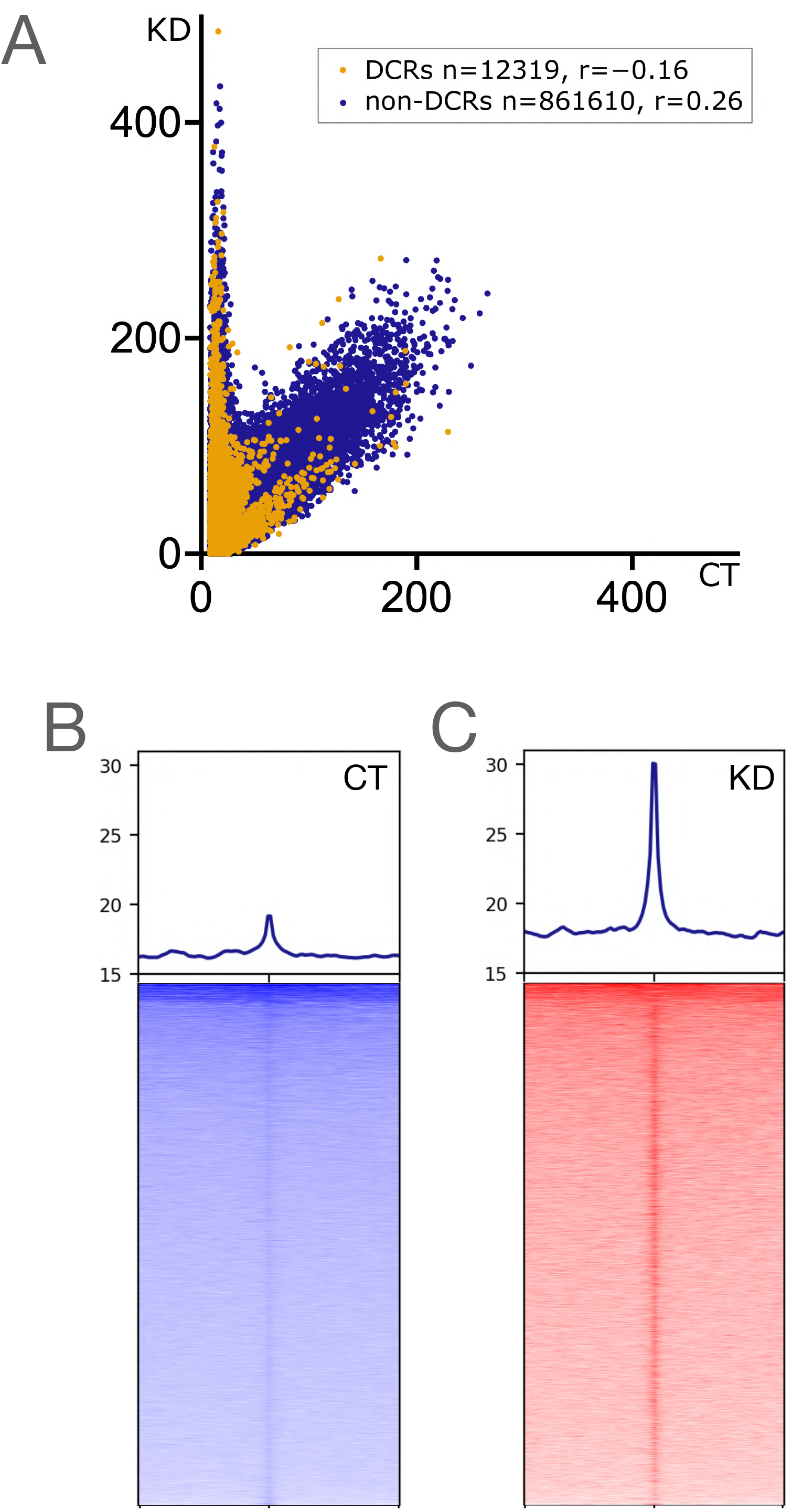
Distribution of G4 signals as captured in pooled CT (CT^*tr*^, CT^*ut*^ and CT^*nuc*^) and KD (KD^*tr*^, KD^*ut*^ and KD^*nuc*^) samples. A: The correlation shows the abundance of G4 signals in the 0.2 kb bins genome-wide (n = 873929). Most 0.2 kb bins show no quantitative difference in the signal (r = 0.26), as represented by the cluster of non-DCRs (dark blue dots). However, a small fraction of the 0.2 kb genomic bins (~0.01%) show a stronger presence of G4 signals in KD than in CT with a negative correlation (r = −0.16) represented by the DCRs (orange dots). The Y-axis represents the input-normalized signal for KD, and the X-axis represents the input-normalized signal for CT per 0.2 kb bins. The DCRs and non-DCRs are negatively correlated. The correlation values for DCRs (yellow dots) and non-DCRs (blue dots) are mentioned in the inset. B-C: The pooled G4 signal in CT and KD show a qualitative difference on the 10 kb flanks from the center of the DCRs. In KD, the center of the DCRs carries higher enrichment of the signal than in CT. In general, the neighboring regions of the DCRs show weaker G4 signals in CT compared to KD. The heatmaps were generated using *deepTools plotHeatmap* with the adjusted Y-axis values for the signal to plot the profile (mean±SE). The DCRs in the heatmaps were sorted in the decreasing order of the signal from the top to the bottom of the plot.

To understand the functional relevance of the DCRs, the proximity of the DCRs with a panel of functional genomic regions was calculated using (i) UCSC annotations for genes and CpG islands, (ii) FANTOM annotations for enhancers, and (iii) ChIP-seq data for markers of replication origins in human cells. There were no obvious spatial associations of the DCRs with any of these functionally annotated landmarks (Fig S17). However, we could establish that in previously reported publicly available G4-ChIP datasets, there is a weak but specific enrichment of G4 signals at DCRs (Fig S18), suggesting that the DCRs are recognized by 1H6 as well as BG4 antibodies. We argued that if the DCRs discovered by the application of the *AbC* G4-ChIP were indeed biologically relevant, they would exhibit features predictable by the known functions of CGGBP1. We analyzed the properties of these regions with reference to the features of DNA sequences associated with regulation by CGGBP1: interspersed repeats, GC-richness, G/C-skew and CTCF-binding sites^43,45,51,52^.

As the net capture of G4s at DCRs was increased due to CGGBP1 knockdown, it was a strong possibility that CGGBP1 occupancy on the DNA mitigates G4 formation. To test this possibility further, we employed a polymerase stop assay as described elsewhere^53^ using the Control DNA and recombinant or immunoprecipitated CGGBP1 (detailed protocol in methods). The Control DNA formed stable G4s in the presence of KCl compared to LiCl (Fig S19A). Pre-incubation of the Control DNA with recombinant CGGBP1 (rCGGBP1) caused a stronger polymerase stop in the presence of LiCl than KCl, resulting in poorer amplification in the presence of LiCl (Fig S19, B and C). Since G4-formation is relatively incompatible with LiCl, it could be concluded that the polymerase stop observed in the presence of LiCl was caused by rCGGBP1 occupying the Control DNA template in the absence of G4s. Such a polymerase-stopping effect by rCCGBP1 was not observed in the presence of KCl. We could recapitulate these findings by using CGGBP1 purified from HEK293T cells as well. Immunoprecipitated and freshly eluted CGGBP1 (Fig S19D) under low salt conditions was used in the polymerase stop assays. A non-specific isotype immunoprecipitate was used as a negative control. Similar to the effect of rCGGBP1, the immunoprecipitated CGGBP1 also exerted polymerase stop in conditions where G4s are not favoured (Fig S19E). We could establish that even in the presence of KCl, where G4 formation on Control DNA is strongly favoured, a pre-incubation with immunoprecipitated CGGBP1 reduced the amount of G4s that could be pulled down using the 1H6 antibody *in vitro* (Fig S19E). Based on these results, we concluded that the presence of CGGBP1 hinders G4 formation through a mechanism that involves the interaction of CGGBP1 with unfolded G4-forming DNA. Such a mechanism could underlie the net increase in G4s upon CGGBP1 depletion observed at the DCRs.

We hypothesized that the CGGBP1-binding sequences would be overrepresented in the DCRs compared to the non-DCRs and tested this using the CGGBP1 ChIP-seq data published elsewhere^43^. Short interspersed nuclear elements (SINEs), of which Alu elements are the most prominent binding sites for CGGBP1, were indeed observed at a lower level at DCRs than that expected from the genomic average or the non-DCRs (Table S10). Such a difference was not observed for LINE elements, another major interspersed repeat regulated by CGGBP1 but not its major binding site. The DCRs where G4 formation was mitigated by CGGBP1 were expectedly rich in GC content (Fig 5A) as Alu elements are concentrated in GC-rich segments of the genome. Since the binding of CGGBP1 was expected to correlate oppositely with G4 formation, we expectedly observed a very weak CGGBP1 occupancy at any population of G4s, including DCRs as well as non-DCRs (Fig 5B). At the DCRs, CGGBP1 occupancy showed a weak but central enrichment, suggesting that these regions are in flux between ‘CGGBP1-bound G4-free’ or ‘CGGBP1-unbound G4-formed’ states. These results were striking as the CGGBP1 occupancy data were derived from a different cell line and suggest that the formation of G4s is a widespread property of DNA even under chromatin constraints, and CGGBP1 is similarly deployed in various cell types to mitigate G4 formation.

**Fig 5:**
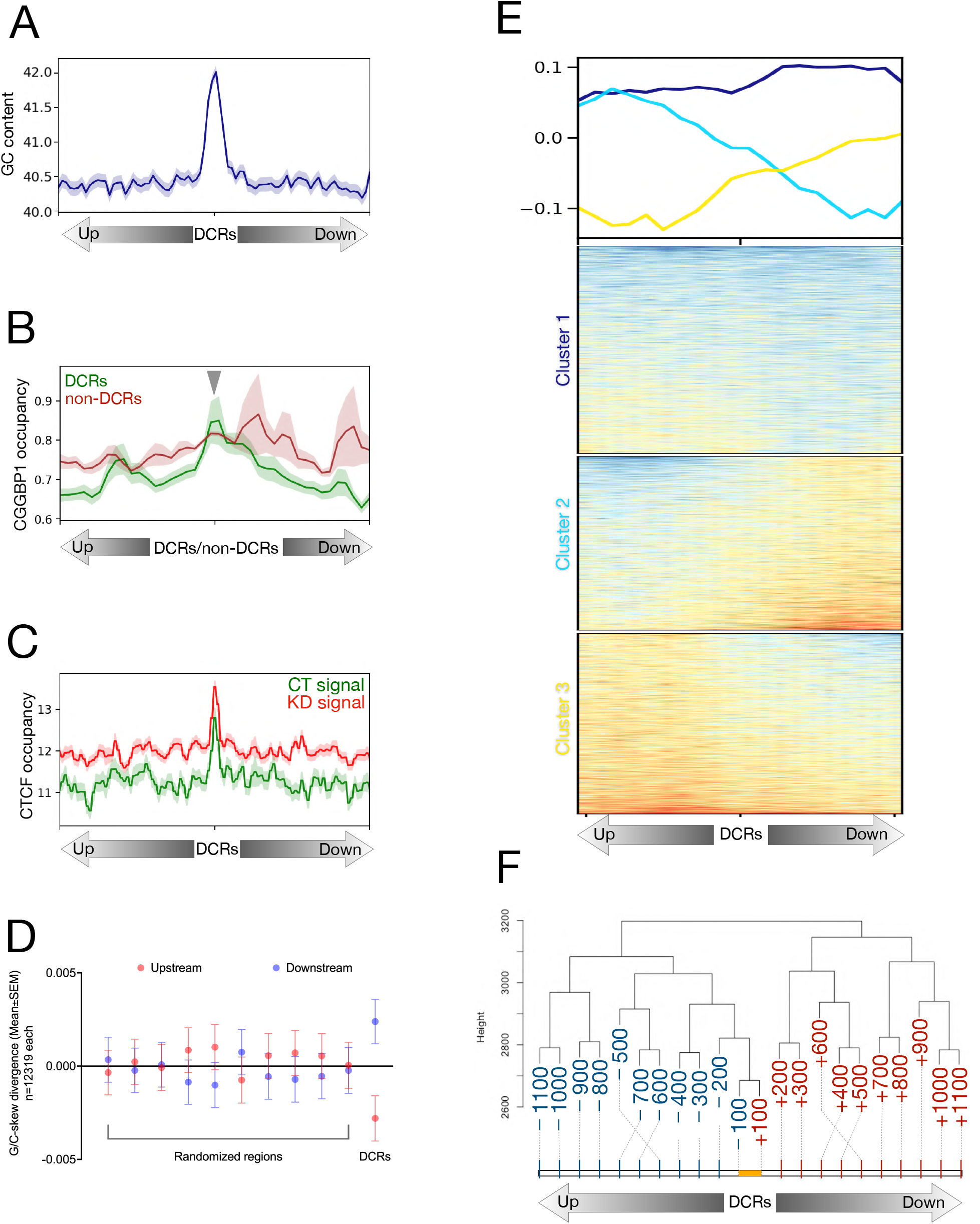
Features of the DCRs. A: The DCRs show that these CGGBP1-regulated genomic regions have stronger GC content than their immediate flanks of 2 kb. The GC content signal is plotted as mean±SE, and the Y-axis represents the signal values. B: The signal from genome-wide occupancy of CGGBP1 under growth stimulation (GSE53571) has been plotted on the 1 kb flanks from the center of DCRs and non-DCRs. The DCRs (green) show enrichment of CGGBP1 occupancy in their centers as compared to the non-DCRs (dark red) with a background level of the signal devoid of any specific enrichment. The lines were drawn as mean±SE, and the Y-axis shows the signal values. C: The non-canonical CGGBP1-dependent CTCF binding sites show that upon knockdown of CGGBP1, the occupancy of CTCF on DCRs has heightened occupancy (dark red) than seen at normal levels of endogenous CGGBP1 (green). The signal is plotted in 10 kb flanks from the center of the DCRs with the lines representing the mean±SE of the signal and the Y-axis representing the signal values. D: The DCRs show a quantitative stronger G/C-skew as measured in their 1 kb flanks compared to the same number of randomly expected genomic regions sampled 10 times. The randomly expected genomic regions show a similar difference in G/C skew on their 1 kb flanks as compared to the DCRs. E: The heatmap shows a qualitative and asymmetric distribution of G/C-skew on the 1 kb flanks of the DCRs. The DCRs are clustered to show stronger changes in G/C-skew in the flanks (cluster 2 and cluster 3) as well as minimum to no change in the G/C-skew (cluster 1). The lines represent the mean signal of G/C-skew calculated for the DCRs. F: The G/C-skew has been binned into 0.1 kb bins in 1 kb flanks of DCRs and forms distinct clusters. The upstream and downstream bins form separate clusters wherein the respective bins cluster differently with the proximal bins. The yellow line represents the length of DCRs and the numbers represent the distance (0.1 kb) from the center of the DCRs either upstream (blue) or downstream (red). The Y-axis shows an arbitrary distance from the 0.1 kb bins. The dashed lines indicate the end of the 0.1 kb bins. The 0.1 kb bins are not drawn to scale. The cluster has been generated in R using the hierarchical method “manhattan”.

G/C-skew is a property of DNA regions regulated by CGGBP1^45^. This includes non-canonical CTCF-binding sites described in HEK293T cells at which CTCF occupancy depends on CGGBP1^5^. The DCRs were also rich in CTCF-binding sites (Fig 5C). However, the CTCF-binding was observed at comparable levels in non-DCRs as well (not shown). It seems that the CTCF occupancy coincides with G4 prevalence and the regulation of CTCF occupancy or G4 formation by CGGBP1 may be acting through independent mechanisms. A G/C-skew calculation showed that the DCRs, including 1 kb flanking regions, had an unexpectedly high G/C-skew (Fig 5D). Interestingly, the G/C-skew was asymmetrically distributed across the DCRs (Fig 5E). The upstream and downstream 0.1 kb regions spanning in the 1 kb flanks were segregated into separate clusters (Fig 5F) due to unexpectedly high transitions of G/C-skew at DCRs.

Collectively, our findings showed that using *AbC* G4-ChIP, we could reliably capture and identify G4s genome-wide from the cells. *AbC* G4-ChIP minimizes the capture of false positive G4s (the population most common between G4-ChIP and G4 DNA-IP^*nat*^). It also captures a previously unreported set of G4s (the population most common between G4 DNA-IP^*denat*^) and thus eliminates false negatives. We could apply this method to HEK293T cells with and without the endogenous levels of CGGBP1 (CT and KD, respectively) and discover regions where mitigation of G4 formation by CGGBP1 coincides with unexpected G/C-skew and CGGBP1-regulated CTCF occupancy.

## DISCUSSION

G4s are some of the most well-studied DNA secondary structures. Decades of *in vitro* studies have defined the chemical environment (presence of KCl) in which the quadruplexes can form on some canonical sequences. Very important and lasting conclusions about the nature of bonding leading to the formation of G-tetrads and the resultant formation of quadruplexes have been generated using *in vitro* assays. A combination of G-triplets separated by spacers has long been considered model sequences for G4 formation. Many computational tools rely on such sequences to report potential quadruplex formation. However, two major advances have challenged the restriction of quadruplexes to such canonical sequences. First, the antibodies such as 1H6 detect G4s formed by a variety of T and G combinations not limited to the model sequences. Second, characterization of G4s genome-wide using the anti-G4 antibody BG4 does not show any restriction of quadruplexes to the canonical sequences. In fact, these studies underscore the complexity of DNA secondary structure landscape in cellular nuclei.

Simplistically, it is expected that the G4-ChIP sequencing datasets are likely to have signature sequences at peak centers, using which a G4 consensus sequence can be derived. However, such a signature sequence for G4s in G4-ChIP datasets has remained elusive. This is reaffirmed by the findings reported elsewhere^36^ as well as by running the G4-ChIP sequencing data through programmes such as *allquads*. The description of G4 landscapes from chromatin available in the literature so far suffers from a lack of calibration of the G4 potential of the genomic DNA. After all, G4 formation is a property of the DNA not necessarily contingent upon any chromatin component other than the DNA itself. Calibration of the G4 potential of the genomic DNA would give a reference frame of the possible G4s in the genome against which the chromatin-derived G4 landscape could be compared. In the absence of such a calibration of the G4 potential of the genome, it becomes difficult to gauge the completeness and genuineness of the chromatin-derived G4 landscape. This is especially important because unlike the ChIP-based characterization of DNA-protein interactions, the G4 landscape characterization from cells is not based on a presumed DNA-protein interaction and yet employs the same biochemical principles in the G4-ChIP protocol. The G4s are expected to be captured in G4-ChIP if they survive the primary treatments of the chromatin with formaldehyde, detergents and sonication (proneness to false negatives) or form *de novo* (proneness to false positives) during these steps. A comparison against the total G4 potential of the human genomic DNA allows a more objective evaluation of these two types of errors in G4 landscapes captured from the chromatin.

The two *in vitro* DNA-IPs described in this study provide a G4 calibration potential for human genomic DNA. Although we performed these assays on DNA purified from HEK293T cells, the G4 potential of human DNA shown by these assays is expected to be near-universal for any human DNA except for any effects of DNA chemical modifications, such as cytosine methylation, on G4 formation. Interestingly, we found two different types of G4s: one which formed by spontaneously overcoming the double strandedness at 4°C and another which formed readily when the strand separation is facilitated, as we achieved through heat denaturation. Expectedly, the second type of facilitated G4s was widespread, G/C-rich and consisted of the spontaneously forming G4s as a subset, whereas the spontaneously forming G4s were limited and relatively G/C-poor. By comparing against this superset of possible G4s that can form in the human genomic DNA, the following possibilities can be tested for any chromatin-derived genome-wide G4 landscape: What fraction of the possible G4s form *in vivo* on the chromatin? What fraction of the chromatin-derived G4s are spontaneous or facilitated types of G4s? Is G4 formation in the chromatin facilitated such that the G4s form in the chromatin beyond the *in vitro* determined G4 potential of the DNA, or is there a mitigation of G4 formation in the chromatin as proposed elsewhere^46^? In order to answer these questions precisely, we must be able to compare an absolutely *in vitro* generated G4 potential of genomic DNA against an absolutely chromatin-derived G4 landscape with no possibility of any false positives or false negatives. Our description of the *AbC* G4-ChIP alongside the *in* vitro-derived G4 potential provides a powerful dataset to understand the scope and nature of G4 formation in chromatin that is largely free of false positive and false negative G4s. We observe that the spontaneously forming G/C-poor G4s are prevalent in G4-ChIP but nearly absent from *AbC* G4-ChIP, showing a reduction in the false positive G4 capture. This reduction of spontaneously forming G4s in *AbC* G4-ChIP is due to the elimination of any post-fixation and sonication incubation with the antibodies. Also, the limited but larger presence of G/C-rich facilitated G4s in the *AbC* G4-ChIP as compared to that in G4-ChIP shows the elimination of false negative G4s. This larger scale of G4 capture in *AbC* G4-ChIP is due to the usage of antibodies for the stabilization of G4s formed in the chromatin. In the G4-ChIP protocol, such G4s would either escape crosslinking if not bound by proteins or escape capture if protein-bound due to steric hindrances to G4-antibody binding.

The disconnect between our knowledge from *in vitro* studies on G4s and the complex G4 landscapes discovered in cells also leads to questions central to G4 biology: Are there sequence properties other than [G_3_-N_(1-7)_]xn that form the quadruplexes as well? Are there different mechanisms to regulate G4 formation at different types of sequences? Are there protein regulators of G4 formation? What are these proteins? Do they work in a sequence-dependent manner? Some of these questions can be answered by analyzing G4 landscapes and their sequence properties with or without perturbation of putative G4 regulatory genes. Despite our knowledge of multiple G4-binding proteins, such targeted studies to dissect subtypes of quadruplex-forming sequences based on their dependence on putative regulator genes have not been reported.

Our findings using the *AbC* G4-ChIP and Control DNA show that G4 formation *in vitro* is different from the G4 landscape in the cellular nuclei. While the *in vitro* G4 formation is unconstrained, the nuclear environment presents conditions which mitigate G4 formation. The *AbC* G4-ChIP captures the cellular G4 landscape with a discernible reduction in *in vitro* artifacts as compared to the conventional G4-ChIP protocol.

The application of the *AbC* G4-ChIP protocol to a CGGBP1 knockdown system also allows us to query the quadruplex formation and changes therein show overlaps with the sequences previously reported to be regulated by CGGBP1. This validatory detection of differential G4 formation at sequences with properties previously reported for regulation by CGGBP1 shows that the *AbC* G4-ChIP protocol is reliable and reports the quadruplexes such that the data are interpretable with reference to previously reported independent experiments. The *AbC* G4-ChIP avoids the capture of the G4s formed rapidly *in vitro* during the incubation steps, such as the incubation with the antibody. The regulation of G4 by any gene can be best studied only if there is a broad capture of G4s with no *in vitro* background. Functional studies aiming to find G4-regulatory genes, such an *in vitro* background as returned by G4-ChIP would confound conclusions as it is independent of any biological features of the samples and depends solely on DNA strandedness and sequence properties. The *AbC* G4-ChIP protocol offers a precise and identification of G4s from the nuclei with a greater diversity of G4s allowing its usage in G4 regulatory functional studies more reliably. We also discover a generic and widespread sequence property of interstrand G/C-skew that makes the DNA prone to G4 formation. With all the advantages *AbC* G4-ChIP offers over the conventional G4-ChIP, it is a protocol that takes longer to perform, the initial steps of permeabilization and antibody-capture are very sensitive and require nearly 5-10 times more antibody. For subsequent sequencing of the purified *AbC* G4-ChIP DNA, it is likely that pooling of samples from multiple runs will be needed.

Our analysis also reveals that the *allquads* G4 signature is strongly associated with *in vitro* formation of these structures. The G4 formation in the cellular nuclei is relatively less dependent on the [G_3_-N_(1-7)_]xn signature. We did not find any sequence property of G4s enriched in the chromatin-derived G4s using *AbC* G4-ChIP as compared to the *in vitro-* derived G4s. These findings suggest that G4 formation is a biochemical feature of the DNA with no evolutionary selection for stabilization of long-term facilitation of G4. This conclusion is supported by the fact that there are no known G4-binding domains or proteins which would bind to and stabilize G4s. The complex chromatin organization of nuclear DNA would benefit by preventing G4 formation and resorting to resolving them by recruiting helicases only when the prevention of G4 formation fails. It follows that protein binding to G4-forming sequences and capture of G4s at the same regions are hence mutually exclusive, a possibility supported by our observations of poor CGGBP1 occupancy at G4s.

We observe that the constraints of the nuclear environment discretize as well as restrict G4 formation. This is reflected in the number of peaks, which is larger in CT^*nuc*^ than in G4 DNA-IP^*nat*^ but less than in G4-DNA-IP^*denat*^. In the absence of any heat denaturation, the G4 DNA-IP^*nat*^ retains its stable long-term G4 landscape with no significant *de novo* formation of G4s at random regions during the antibody incubation. Most importantly, the G4s captured in G4 DNA-IP^*denat*^ are a range of G4s that are formed upon a range of heat energy input to melt the DNA strands. Apparently, the regions with the highest energy thresholds form the most stable G4s and get identified as peaks. The possibility of overcoming such extreme energetic requirements is an impossibility in the nuclear environment. This suggests that the energetic constraints enforced by double stranded nature of the DNA and its sequence are a biochemical impediment to G4 formation in the nuclear environment. The inverse correlation between G4 DNA-IP^*denat*^ peaks and G4-ChIP or *AbC* G4-ChIP peaks demonstrates this. However, beyond the peaks, the G4 DNA-IP^*denat*^ data contains G4 enrichment which is present in *AbC* G4-ChIP peaks more than in G4-ChIP peaks. The nature of the constraints of the nuclear environment on G4 formation is expected to be widespread, given the hindrances to DNA processes imposed by the random formation of secondary structures. Clearly, there is a larger repertoire of helicases that resolve these secondary structures, including the G4s, than the possibility of some proteins with dedicated quadruplex stabilizing functions. The growing number of G4-interacting proteins does not indicate any such quadruplex stabilizing function, although a paucity of targeted studies tests this possibility.

Our study reports such a targeted test of function for CGGBP1 on G4 formation. CGGBP1 is not a G4-binding protein. Similarly, it is not established for the many G4-binding proteins if their binding is to the actual quadruplex structure or to the immediate vicinity of the structure. This argument is supported by the absence of any quadruplex-binding domain common to these proteins^27^. An additional set of proteins important in this context are those which can bind to the DNA sequences capable of forming G4s but not to the quadruplex structures per se. The known DNA-binding properties of CGGBP1 indicate that it is capable of binding to G/C-rich sequences with G-repeats^27,43,51,52^. These sequences are strong candidates for G4 formation. The binding to quadruplex-forming sequences but not the quadruplexes per se is logically a feature of proteins, which through binding to the DNA, would preclude quadruplex formation. These proteins, such as CGGBP1, would be able to exert constraints of the nuclear environment on quadruplex formation. Our results show that CGGBP1 binds to sequences capable of G4 formation and exerts constraints of the nuclear environment on quadruplex formation. The weak CGGBP1 occupancy at sequences enriched as G4s reaffirms the notion that CGGBP1 binding could be an impediment to G4 formation. The DCRs, with their preference for CTCF occupancy, underscore the potential functions of G4-forming regions as chromatin boundary elements. The preponderance of G4s in the heterochromatin^54^ and mitigation of G4 formation at CTCF-binding sites suggest that heterochromatinization, G4 formation and regulation by CGGBP1 are interlinked phenomena. Interestingly, our results highlight a property of quadruplex-forming DNA sequences, which is much less appreciated in G4 biology; G/C-skew. The *AbC* G4-ChIP presents a powerful technique to decipher the cellular G4 landscape and its regulation and it has the potential to be adapted for discovering any DNA secondary structures genome-wide against which reliable antibodies are available.

## METHODS

### Synthesis and cloning of the Control and the carrier DNA

We have designed and procured a set of oligonucleotides (Table S11) to generate the Control and the carrier DNA by PCR amplification. The synthesized fragments were gel-purified and cloned into the pGEM-T Easy vector (A1360; Promega) as per the manufacturer’s protocol. The carrier DNA has the same order of base composition, however, the GGG in [*G_3_-N_5_-G_3_-N_5_-G_3_-N_5_-G_3_*] were replaced with ATG. The sequences of the control and carrier DNA are provided below: The sequence of the Control DNA: 5’-TAATACGACTCACTATAGGGCACCCCTACATCGCAGCGGTCTTTCGGGCTAACGGGATCATGGGACTCAGGGGACAGCCTCAAGCAACATGCCCAGTCCTGACCTTCAATAAGGAAGCAAACTGGGAAGGAGGGTGTCAGGGATAAAGGGGAGTCCTAGTCAAGGTGTCGGATGTCCTAAGACTTATGATCATTTTCTTAGGGTCTAAGGGCTCGAGGGTCTGCGGGTCGGTTTCCTTCTAGAATTAGTATCTTCTATAGTGTCACCTAAAT-3’

The sequence of the carrier DNA: 5’-TAATACGACTCACTATAGGGCACCCCTACATCGCAGCGGTCTTTCATGCTAACATGATCATATGACTCAATGGACAGCCTCAAGCAACATGCCCAGTCCTGACCTTCAATAAGGAAGCAAACTATGAAGGAATGTGTCAATGATAAAATGGAGTCCTAGTCAAGGTGTCGGATGTCCTAAGACTTATGATCATTTTCTTAATGTCTAAATGCTCGAATGTCTGCATGTCGGTTTCCTTCTAGAATTAGTATCTTCTATAGTGTCACCTAAAT-3’

Biophysical characterization of control DNA sequence:

### Sample preparation

Commercially synthesized single-stranded 5’-GGGTCTAAGGGCTCGAGGGTCTGCGGG-3’ (oligo1) and 5’-CCCGCAGACCCTCGAGCCCTTAGACCC-3’ (oligo2) oligodeoxyribonucleotide sequences were purchased from Bioserv Biotechnologies, India. These oligodeoxyribonucleotides were diluted to a final concentration of 40 μM in 50 mM Tris-HCl (pH 7.4) buffer with different KCl concentrations (0.05 M, 0.1 M, 0.15 M, 0.5 M and 1 M) for oligo1 and 0.05 M KCl concentration for oligo2. Subsequently, the DNA samples (oligo1 alone or called DG) and oligos 1 and 2 together (called WC) were heated at 95°C for 5 minutes and cooled down to room temperature for 3 hours. The secondary structural conformational preferences of the DNA oligonucleotides were explored by acquiring the CD spectra immediately after the sample preparation.

### Circular Dichroism (CD) spectroscopy

The CD spectroscopic study was carried out in a JASCO-1500 spectrophotometer at 25°C, and the spectra were processed using spectral manager software (www.jascoinc.com). The spectral data were collected in triplicate in the wavelength region of 190-320 nm with 1 nm bandwidth in a 1 mm path length cuvette. The baseline correction was done using the Tris-HCl buffer in the presence of an appropriate KCl concentration. Note that only 0.05 M KCl was used for WC, unlike DG. The triplicate average of the CD spectra was considered for the analysis. Further, the thermal melting was carried out for the DG by heating the sample in the range of 15°C to 95°C temperature at 1°C interval and a 30-second incubation time before the acquisition of data at each temperature. The triplicate averaged thermal denaturation curve was used for the analysis.

### Electrophoretic mobility shift assay

Electrophoretic mobility shift assay (EMSA) was carried out using 14% native polyacrylamide gel pretreated with ethidium bromide (EtBr). The native polyacrylamide gel was prepared using 1X TAE buffer (pH 7). Subsequently, 25 μM concentration of the DG sample in 0.05 M Tris-HCl (pH 7.4) buffer with varying concentrations of KCl (0.05 M to 1 M) was mixed with 50% of glycerol and loaded into the well. The electrophoresis was carried out using 1X TAE running buffer at 60 V, 4°C for two hours. After the completion of electrophoresis, the gel was photographed under UV light using SmartView Pro 1100 Imager System, UVCI-1100 from Major science.

### Modeling of DNA conformation

Based on the information obtained from CD and EMSA, the secondary structural preference of DG was modelled using 3D-NuS web server^55^. Pymol 2.1.1 software package (https://pymol.org/2/) was used for the visualization of the generated models.

### Cell culture and transfection of control DNA

HEK293T cells were transduced with a control (CT) or CGGBP1-targeting (KD) shRNA as described elsewhere^5^. These cells were maintained in DMEM (SH30243.01, Cytiva HyClone) supplemented with 10% heat-inactivated fetal bovine serum (FBS) (10270106, Invitrogen) and selected with 0.3 μg/ml of the final concentration of puromycin (CMS8861; HiMedia). For the equality of transfection of the Control DNA, the cloned-in carrier DNA in the same vector was cotransfected in CT and KD using the calcium-phosphate transfection method. 1.25 μg of each of the Control DNA and the carrier DNA (cloned in pGEM-T Easy vector) and 20 μl of 2M CaCl_2_ were mixed. A final volume of 200 μl is attained with sterile autoclaved water. This mix was added dropwise in 200 μl of 2X HEPES buffered saline (51558; Sigma) with gentle agitation on a vortex mixer to form calcium phosphate-DNA precipitate. The precipitate was incubated at room temperature for 20 minutes and added drop-by-drop on approximately 2 × 10^6^ cells per 10 cm dish. On the next day, the cells were processed for *AbC* G4-ChIP.

### *in vitro* immunoprecipitation of the Control and the carrier DNA using 1H6

1.25 μg of each of the Control DNA and the carrier DNA were mixed in a 1.5 ml centrifuge tube at room temperature, and the volume was made up to 100 μl using deionized autoclaved water. 22.5 μl of 1 M KCl was added to the final concentration of 150 mM in a final volume of 135 μl. This mix was heated in a dry bath at 95°C for 5 minutes. It was cooled down at room temperature and continued for a further 30 minutes to allow the formation of G4s. Following this incubation, 15 μl of fetal bovine serum (SV30160.03; Cytiva HyClone) was added to the final concentration of 10% v/v. The formed G4s were allowed to be recognized by 1 μl (1 μg) of Anti-G4 1H6 antibody. The mixture was flicked a couple of times and incubated at 4°C for 1 hour. The antibody-G4 interaction was crosslinked using 4 μl of 37% formaldehyde to the final concentration of 1% at room temperature for 10 minutes with a gentle invert-mixing. 15 μl of 1.25 M glycine was added to the final concentration of 125 mM, gently mixed and incubated for 5 minutes to quench the reaction. These antibody-crosslinked G4s were included in the cell lysates for samples CT^*ut*^ and KD^*ut*^ and sonicated, as mentioned in *AbC* G4-ChIP.

For *in vitro* assays described in figure S17, G4s formed on Control DNA *in vitro* were immunoprecipitated using the same protocol as described above. Eluted products were subjected to qPCR assays as described later.

### G4 DNA-IP

Genomic DNA from HEK293T cells was isolated using the phenol-chloroform method stored in 10 mM Tris (pH 8) at −20°C. 1 μg of genomic DNA was denatured at 95°C for 5 minutes and then snap-chilled for the sample G4 DNA-IP^*denat*^. Following this step, the sample was incubated with 10% v/v FBS. At this point 1 μg of genomic DNA, for the sample G4 DNA-IP^*nat*^ also incubated with 10% v/v FBS. The rest of the steps were followed as described for the *in vitro* immunoprecipitation of the Control and carrier DNA.

### *AbC* G4-ChIP

G4s in CT^*nuc*^, KD^*nuc*^, CT^*tr*^, KD^*tr*^, CT^*ut*^ and KD^*ut*^ were captured using the *AbC* G4-ChIP protocol. The culture medium was removed from 10 cm dishes with or without transfection for each sample. The cells were gently washed with ice-cold 1X Ca^2+^-free PBS (SH30256.01, Cytiva HyClone) twice. After the last wash, the cells were treated with 1 ml of trypsin per dish and incubated at 37°C for 10 minutes. Following the trypsinization, 5 ml of serum-containing DMEM was added to each dish, and the cells were declumped by vigorous pipetting. Then the cells were collected in a clean 50 ml screw-capped centrifuge tube by passing the cell suspension through a 40 μm cell strainer (431750; Corning). The volume was adjusted to a total of 30 ml with serum-free DMEM. The cells were crosslinked with 4 ml of 37% formaldehyde stock solution and incubated for 5 minutes at room temperature to fix DNA-protein interactions. The suspension of cells was quickly quenched with 4 ml of 1.25 M Glycine along with 4 ml of 1 M KCl. The suspension was quickly spun down gently at 1,250 rpm for 5 minutes at room temperature and resuspended in 5 ml of Tris-KCl buffer (50 mM Tris, pH 7.5; 150 mM KCl). The cells were washed once more in the Tris-KCl buffer, spun down at 1,250 rpm for 5 minutes at room temperature, and the supernatant was carefully removed. The cell pellet was resuspended and permeabilized in 15 ml of Tris-KCl buffer supplemented with 0.5% of Triton X-100 for 10 minutes at room temperature with gentle invert mixing. Following the incubation, the suspension was spun down gently at 1,250 rpm for 5 minutes at room temperature and resuspended in 15 ml of Tris-KCl buffer supplemented with 0.01% v/v of Triton X-100 (called TKT buffer) to maintain the permeability of the cell membrane. To the suspension, 10% v/v FBS was added immediately and incubated at room temperature for 30 minutes (for the first 15 minutes with intermittent invert-mixing and then stationary until the end of the 30th minute). The suspension was spun down at 1,250 rpm for 5 minutes at room temperature, and all but 5 ml of the supernatant was gently removed without disturbing the pellet. The pellet was resuspended gently with the help of a pipette to avoid mechanical rupturing of the cells, and the suspension of cells was then transferred into a fresh 15 ml screw-capped tube. This suspension of cells was incubated with anti-G4 1H6 antibodies (Sigma; MABE1126) as 5 μg of antibody/100 μl of cell pellet for two hours at room temperature. Following the incubation, the suspension was spun down at 1,250 rpm for 5 minutes at room temperature, and the pellet was gently washed once with the TKT buffer and resuspended again in the same. The antibody-G4 interactions were then crosslinked with 1% final concentration of formaldehyde and incubated at room temperature for 10 minutes with gentle invert-mixing. 125 mM of the final concentration of glycine was added to quench formaldehyde, and the mix was incubated at room temperature for 5 minutes. The cells were pelleted down at 1,250 rpm for 5 minutes at room temperature and lysed in SDS lysis buffer (50 mM Tris-HCl, pH 8.0; 1% SDS and 25 mM EDTA), supplemented with 1X v/v Halt Protease and Phosphatase inhibitor (87786; Thermo Scientific), twice the volume of cell pellet. The cells were lysed on ice for 30 minutes with intermittent tapping of the tube. The lysate was equally distributed into 1.5 ml centrifuge tubes (no more than 500 μl/tube) and sonicated in Diagenode Bioruptor sonicator for 20 cycles (with 30 seconds ON - 30 seconds OFF) at constant preset power. The sonicated fraction was quickly pelleted down at 4°C and the supernatant was stored separately at 4°C. To increase the recovery of sonicated fraction, the pellet was resuspended in an equal volume of ChIP dilution buffer (0.01% SDS; 1% Triton X-100; 1.2 mM EDTA; 16.7 mM Tris-HCl, pH 8.0 and 167 mM NaCl) as the volume of sonicated fraction stored earlier. At this stage, the antibody-crosslinked Control DNA (for samples CT^*ut*^ and KD^*ut*^) was included and sonicated for 10 cycles (with 30 seconds ON - 30 seconds OFF) at constant power. The respective sonicated fractions were pooled and centrifuged at 14,000 rpm for 30 minutes at 4°C. Meanwhile, Protein G Sepharose^®^ 4 Fast Flow (GE; 17061801) beads were washed thrice gently with 1X PBS. The beads were gently centrifuged during every wash at 1,000 rpm for 3 minutes to remove the supernatant. The clear sonicated fraction was transferred into a 15 ml tube and diluted with the ChIP dilution buffer to ten times of the volume of SDS lysis buffer used. 10% of this diluted fraction was kept as input. In the rest, 10% v/v slurry of beads was added and tumbled at low rpm for two hours at 4°C. The beads were gently collected by centrifugation at 1,000 rpm for 3 minutes at room temperature and washed with PBST (0.1% Triton X-100 in 1X PBS) five times. The last wash supernatant was removed. Finally, the beads were vigorously vortexed in the presence of an elution buffer (1% SDS; 0.1 M sodium bicarbonate) for 30 minutes. The elution was pooled in 500 μl volume by collecting 250 μl of eluted fraction after 15 minutes and another 250 μl at the end of the 30th minute. The eluted fractions and the inputs were subjected to reverse-crosslinking by adding 20 μl of 5 M NaCl and incubating overnight at 65°C. The reverse-crosslinked DNA was digested with Proteinase K (2 μl of 10 mg/ml stock) in the presence of 10 μl of 0.5 M EDTA and 20 μl of 1 M Tris-HCl (pH 6.8) at 42°C for 1 hour. The DNA was purified using the ethanol precipitation method and dissolved in deionized autoclaved water.

For each sample, eight 10 cm dishes were pooled and subjected to the *AbC* G4-ChIP protocol.

### Library preparation, basecalling, sequence alignment and annotation

The CT^*nuc*^, KD^*nuc*^ and Input^*nuc*^ were sequenced on the IonTorrent S5 sequencer while CT^*tr*^, KD^*tr*^, CT^*ut*^, KD^*ut*^ and respective inputs were sequenced on MinION Mk1 B device from the Oxford Nanotechnologies (ONT) sequencing platform. The sequencing libraries were prepared with either Ion XpressTM Plus Fragment Library Kit (Cat. no. 4471269) or PCR sequencing kit (SQK-PSK004) for the ONT platform as per the manufacturer’s instructions. To generate sufficient sequencing depth, the sequencing library for each sample was pooled from four replicates every time. The samples G4-ChIP, *in vitro* G4 DNA-IPs and the corresponding inputs were sequenced on MinION Mk1B as well. The base-calling of the reads was carried out in real-time. For reads sequenced on MinION, the poor-quality reads were further subjected to basecalling by a locally installed GPU-based *guppy_basecaller*. The sequenced reads were subjected to removal of the sequencing adapter using Porechop using *default* options. The repeat-unmasked hg38 genome (excluding unannotated, mitochondrial chromosomes and chromosome Y) with or without the control DNA was indexed by using *minimap2 -x map-ont* option. The filtered reads were aligned using *minimap2* on the indexed genome, however, the samples, CT^*nuc*^, KD^*nuc*^ and Input^*nuc*^, were aligned on the indexed genome without the control DNA. Only uniquely aligned reads were fetched for further analyses. A detailed description of the analyses is mentioned in Appendix-I. The sequence data are deposited to NCBI GEO vide ID GSE202456.

### Immunoprecipitation of CGGBP1 of HEK293T cells

One fully confluent 10 cm dish of HEK293T was washed with ice-cold 1X PBS and pelleted at 1,250 rpm in a 1.5 ml microcentrifuge tube at room temperature. The pellet was lysed with RIPA lysis buffer (50 mM Tris, pH 8; 150 mM NaCl; 1% NP-40; 0.5% sodium deoxycholate and 0.1% SDS) added with 1% v/v of Halt Protease inhibitor cocktail at 1:1 (pellet:RIPA lysis buffer) ratio. The lysis was carried out on the ice for 30 minutes with gentle tappings every 10th minute. The lysate was spun down at 14,000 rpm for 30 minutes at 4°C. 0.05% of the clear supernatant was stored as input, while in the rest, 2 μl of anti-CGGBP1 antibody (PA5-57317; Invitrogen) was added and incubated overnight at 4°C. Following the incubation, a 10% v/v slurry of protein A agarose beads was added and incubated at room temperature for two hours. The beads were then washed thrice with RIPA lysis buffer, spun at 1,000 rpm for two minutes, and the supernatant was discarded. The beads were finally washed with icecold 1X PBS, spun at 100 rpm for two minutes, and the supernatant was discarded. The beads were then subjected to vigorous vortexing in the presence of elution buffer (0.2M glycine and 10 mM Tris, pH 8) and eluted by centrifugation at high speed. The eluted CGGBP1 was stored at −20°C until further use.

### Quantitative PCR-stop assay using Control DNA and CGGBP1

Quantitative PCR (qPCR) stop assay was performed as described elsewhere^53^. The polymerase employed in our qPCR stop assays was exonuclease and proof-reading activity deficient Klenow fragment (M0212L; New England Biolabs). The Control DNA (Fig 3A) was generated with a T7 sequence (5’-TAATACGACTCACTATAGGG-3’) at the 5’ end and an SP6 sequence (5’-TTCTATAGTGTCACCTAAAT-3’) at the 3’ end. The G-rich strand of the Control DNA was generated using T7 primer and used as a template in qPCR stop assays. 100 ng of double-stranded Control DNA was used as a template per reaction with 250 uM of each dNTP, 10 picomoles of T7 primer, 1X ThermoPol reaction buffer and 1.25 units of Taq DNA Polymerase (M0267L; New England Biolabs) in a 50 μl reaction volume. The following PCR condition was used: (95°C for 5 minutes) X 1 cycle, (95°C for 5 minutes, 55°C for 30 seconds and 72°C for 45 seconds) X 25 cycles and (72°C for 5 minutes) X 1 cycle. The single-stranded DNA generated using this method was purified using PCR purification kit (A9285; Promega) and quantified using Nanodrop. 200 ng of single-stranded G-rich DNA was used in qPCR stop assay. The reaction mix comprised of 250 uM of each dNTP, 10 picomoles of SP6 primer, 1X reaction buffer and 1.5 units of Klenow (M0212L; NEB). The SP6 primer (5’-ATTTAGGTGACACTATAGAA-3’) at the 3’ terminus of the G-rich strand was extended such that the amount and the length of the primer extension products obtained were inversely proportional to the polymerase stop exerted by G4s. As described in various experiments, the purified single-stranded G-rich DNA was incubated with 150 mM KCl (P9541; SIGMA) or 150 mM LiCl (L9650; SIGMA) at 95°C for 5 minutes, followed by at room temperature for 30 minutes before incubation with Klenow for polymerase activity and primer extension. In experiments where the effects of CGGBP1 were studied, the single-stranded G-rich DNA was preincubated with rCGGBP1, or immunoprecipitated CGGBP1 or isotypecontrol IgG immunoprecipitates at room temperature for 30 minutes followed by incubations with KCl or LiCl and the primer extension steps. The single-stranded primer extension assay products were quantified using a primer pair (5’-CAGCCTCAAGCAACATGCCCAGTC-3’ and 5’-AGTTTGCTTCCTTATTGAAGG-3’), located downstream from the SP6 site on the Control DNA closer to the T7 site. The specific amplification from the primer pair was established by resolving the PCR products on agarose gel and visualization using Ethidium bromide staining. qPCRs were performed on a HiMedia Insta Q96 real-time PCR machine using the iTaq Universal SYBR Green Supermix (1725125, Bio-Rad) as the fluorophore for PCR product real-time quantification. Each qPCR was performed in triplicates or more replicates, as indicated in the figures. Melt curves were obtained to ascertain PCR specificity in each run. The Ct values were obtained for amplification curves with comparable efficiencies as early as possible at the onset of the logarithmic phase of the PCR. The Ct values were exported in text files and analyzed using the double delta Ct method in LibreOffice spreadsheets. Data were plotted using GraphPad PRISM. Outliers samples were excluded from the analyses. The statistical significance of differences was calculated using indicated t-tests inbuilt in PRISM.

### Generation of correlation matrices

The human genome was binned into 0.2 kb bins. Coverage of the uniquely aligned reads on those genomic bins was calculated using *bedtools coverage* with the option *-counts*. Any genomic bins with read coverage of less than 3 and more than 100 were excluded. The bins satisfying this criterion were used to fetch the reads from all the samples and the inputs. The reads were then converted into *bam* format using *bedtools bedtobam* function. The bam files were then indexed using *samtools* and converted to *bigWig* signal using *deepTools bamCoverage* with a bin size of 200. These *bigWig* signals were used to compute the correlation using *deepTools multiBigwigSummary*. The heatmaps of correlation matrices were generated using *deepTools plotCorrelation* with the options *-c spearman -p heatmap -- skipZeros --removeOutliers*.

### G/C-skew calculation

The human genome hg38 was binned in 100bp regions using *bedtools makewindows* and fasta sequences were fetched using *bedtools getfasta* from the unmasked hg38. For every bin, the presence of “G”s and “C”s were calculated separately using *emboss fuzznuc*. Any corresponding bin having a *zero* count was eliminated from the G/C-skew calculation. Once all the bins with “G” and “C” counts were available, G/C-skew was calculated using the formula [(G-C)/(G+C)] for each 100 bp genomic bin. This information was used to generate bigWig signal using UCSC *bedGraphToBigWig* for generating heatmap.

### Discovery of Differentially Captured Regions (DCRs)

The coverage of uniquely aligned sequenced reads was computed for all *AbC* G4-ChIP samples in the 0.2 kb genomic bins. The coverage counts were normalized using sequencing depth in the corresponding CT or KD samples. Those 0.2 kb genomic bins were chosen wherein the raw coverage counts follow the criterion of ≥ 3 and ≤ 100 for all three sets of *AbC* G4-ChIP samples. Then the corrected coverage counts were used to calculate the “diff/sum” ratios [(KD-CT)/KD+CT)] for each pair of the *AbC* G4-ChIP samples. The same exercise was performed for the inputs, however, no limit to coverage counts was applied. Thus, the diff/sum ratios for the three *AbC* G4-ChIP samples and three inputs were calculated for each bin individually, and a separate one-tailed T-Test was applied for each bin with n = 3 for CT as well as KD. The 0.2 kb genomic bins with p-value < 0.01 (true discoveries established through q-value < 0.01) were selected and denoted as differentially captured regions (DCRs).

### GGG/CCC weighted content calculation

GGG/CCC weighted content was calculated on all peak datasets. Fasta sequences for the peaks were fetched from the repeat-unmasked hg38 followed by “GGG” and “CCC” patterns were searched using *emboss fuzznuc* with every 1-base succession for each peak. Any peaks with zero count of either “GGG” or “CCC” patterns were discarded for GGG/CCC weighted content calculation. The weighted GGG/CCC content of a peak was calculated by taking the product of the sum of “GGG” and “CCC” counts and the coverage of reads on the given peak. The GGG/CCC weighted content was plotted using GraphPad Prism9.

### Peak calling

Broad peaks were called on the datasets against the corresponding inputs as background noise using the *MACS2 callpeak* function. For all *AbC* G4-ChIP samples and *in vitro* G4 DNA-IP^*denat*^, a bandwidth of 3000 (*--bw 3000*) was used to call the broad peaks with the options: *--nolambda --broad --broad-cutoff 0.01*. However, for G4-ChIP and G4 DNA-IP^*denat*^ samples, broad peaks were called at the default bandwidth value of 300. The details of the discovered peaks are listed in table S2.

### Jaccard Index calculation

Jaccard indices on the peaks were calculated using *bedtools jaccard* option regardless of strandedness. The indices between samples are mentioned in figure S8.

## DECLARATIONS

The authors declare no conflict of interest.

## ACKNOWLEDGEMENTS

The work was supported by the grant CRG/2021/000375 from SERB and IITGN internal support to US. SD was supported by a fellowship from MHRD, GoI through IITGN. MP, DP, AM and CG contributed to this work during their studentships at IITGN. Then MP and DP were supported by CSIR fellowships and CG and AM were supported by MHRD and IITGN. The 1H6 antibody was generously provided by Dr. Peter M. Lansdorp (University Medical Center Groningen). Contributions by CS and TR were at IITH. SD, MP, DP, CG and AM performed experiments under supervision of US, CS performed experiments under supervision of TR, all authors participated in manuscript writing led by US. We acknowledge the anonymous reviewers whose comments and suggestions improved this manuscript significantly.

---

**Fig S1:**
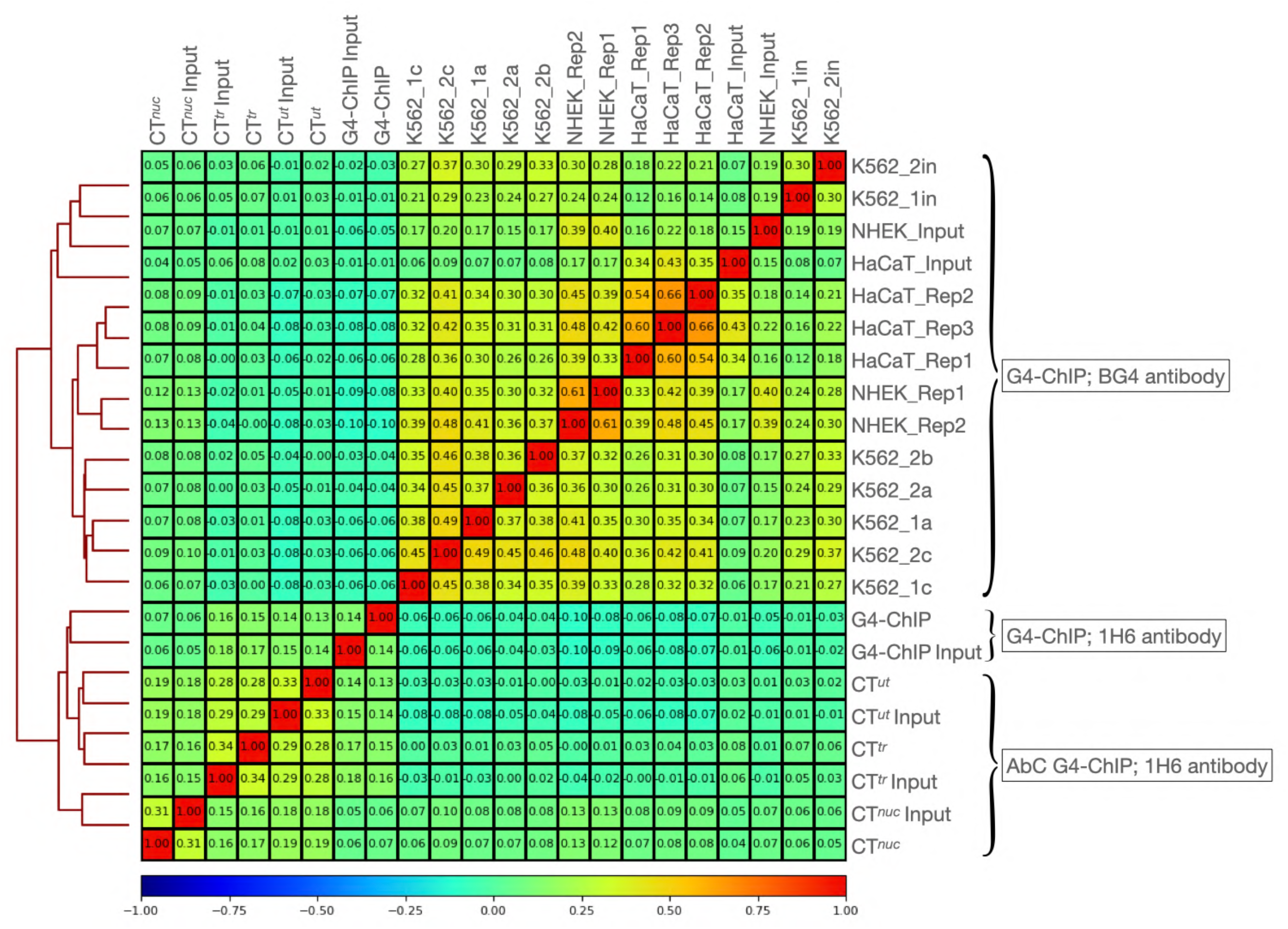
Spearman’s correlation matrix and clustering of G4 profile for *in vitro* G4 DNA-IPs: Genome-wide comparison of signals in IPnat and Ipdenat against a common input shows that the two IPs differ from the input as well as from each other. Thus, the G4 profiles in G4 DNA-IP^*nat*^ and G4 DNA-IP^*denat*^ are very different. However, the slight similarity between G4 DNA-IP^*denat*^ and DNA-IP input is in sharp contrast to the dissimilarity between G4 DNA-IP^*nat*^ and G4 DNA-IP^*denat*^ or DNA-IP input. This suggests the widespread G4 potential of the genome captured in G4 DNA-IP^*denat*^ and a much more restrictive G4 capture in G4 DNA-IP^*nat*^. The signals in the samples were calculated on 0.2 kb bins genome-wide. The numbers indicate Spearman’s correlation coefficients for pairs of samples.

**Fig S2:**
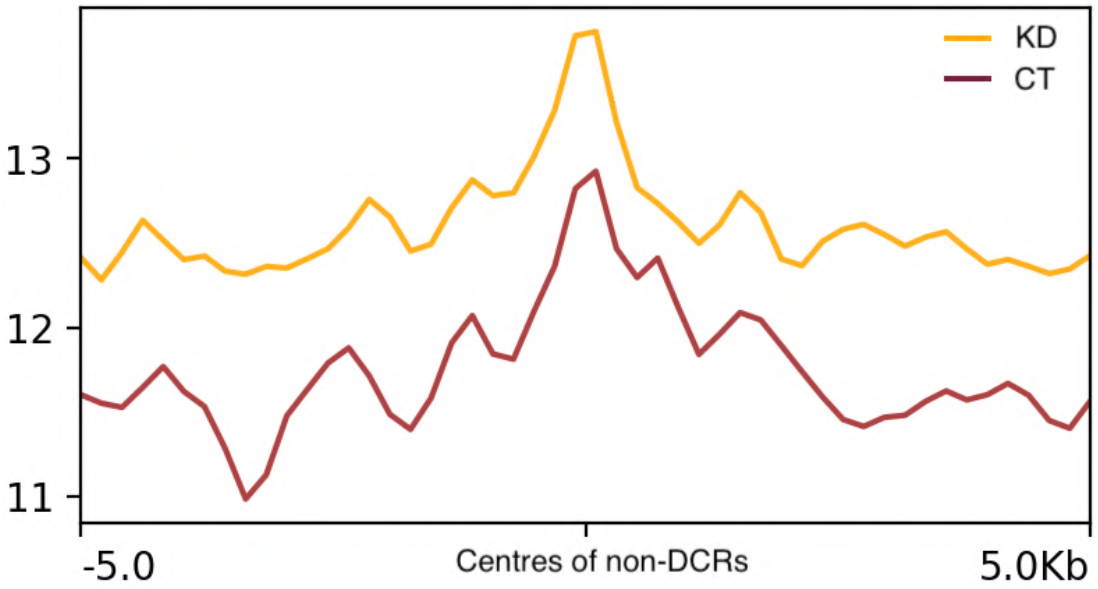
Comparison between G4 DNA-IP^*nat*^ and G4 DNA-IP^*denat*^ signals and peaks genomewide: The G4 signal from G4 DNA-IP^*nat*^ was poor at G4 DNA-IP^*denat*^ peaks (A), whereas the G4 DNA-IP^*denat*^ signals were highly enriched at G4 DNA-IP^*nat*^ peaks (B) suggesting that the G4 DNA-IP^*nat*^ pull-down is a subset of G4 DNA-IP^*denat*^ pull-down. A similar pattern was observed more emphatically at the peak level. The small number of G4 DNA-IP^*nat*^ peaks were almost entirely present as a subset of the G4 DNA-IP^*denat*^ peaks (C), whereas only a minor fraction of the G4 DNA-IP^*denat*^ peaks overlapped with G4 DNA-IP^*nat*^ peaks (D). The signals were plotted on 5 kb flanks from the centers of the peaks. Numbers indicate peaks identified in the respective sample, and the Y axes of the plots have signals plotted as mean±SEM.

**Fig S3:**
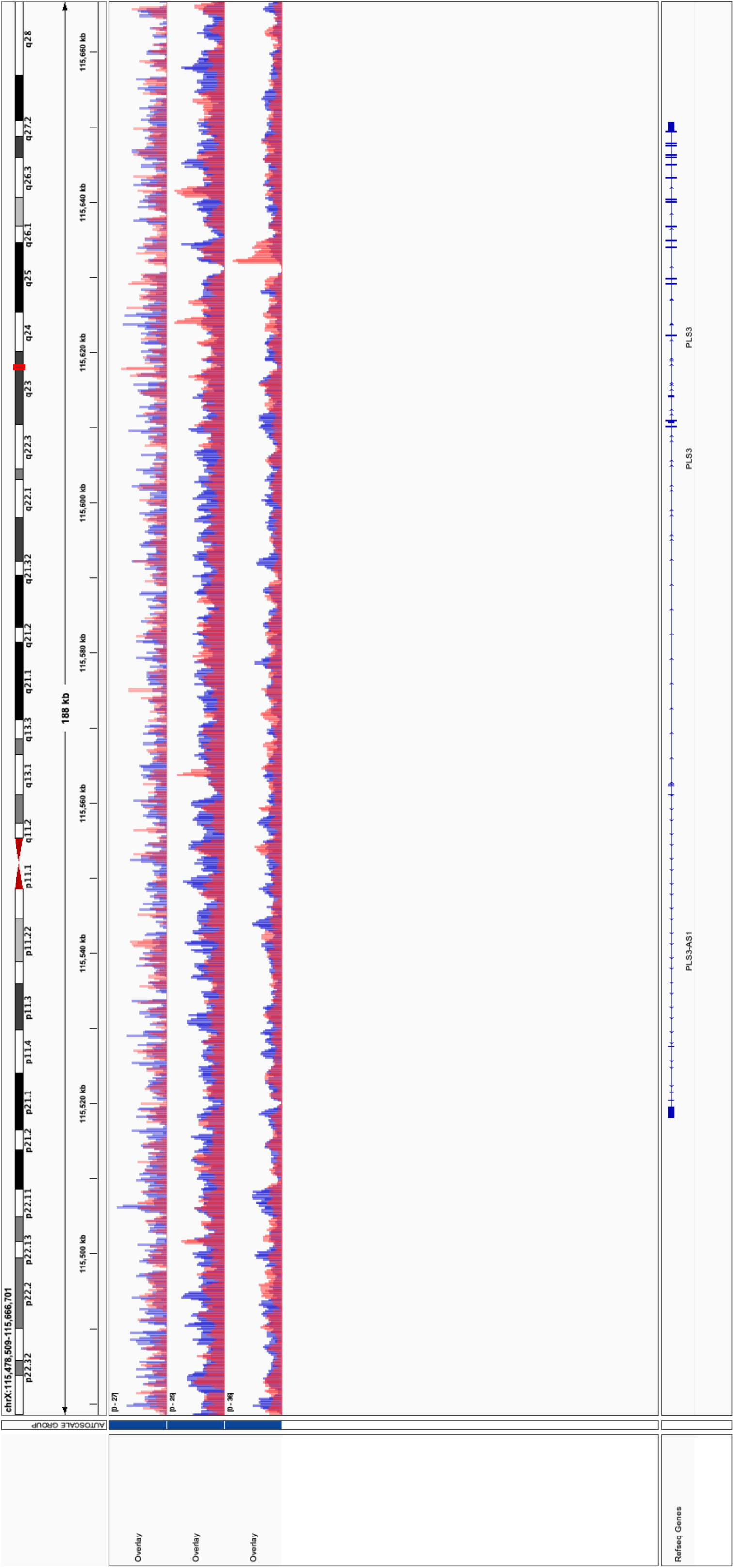
Violin plots of read representation-weighted GGG/CCC counts in peaks (product of GGG/CCC count and the number of reads mapping to each peak). The Y-axis shows the median and IQR on a log10 scale. Remarkably, CTnuc has the lowest GGG/CCC representation in peaks. Peak counts and properties are presented in table S2.

**Fig S4.**
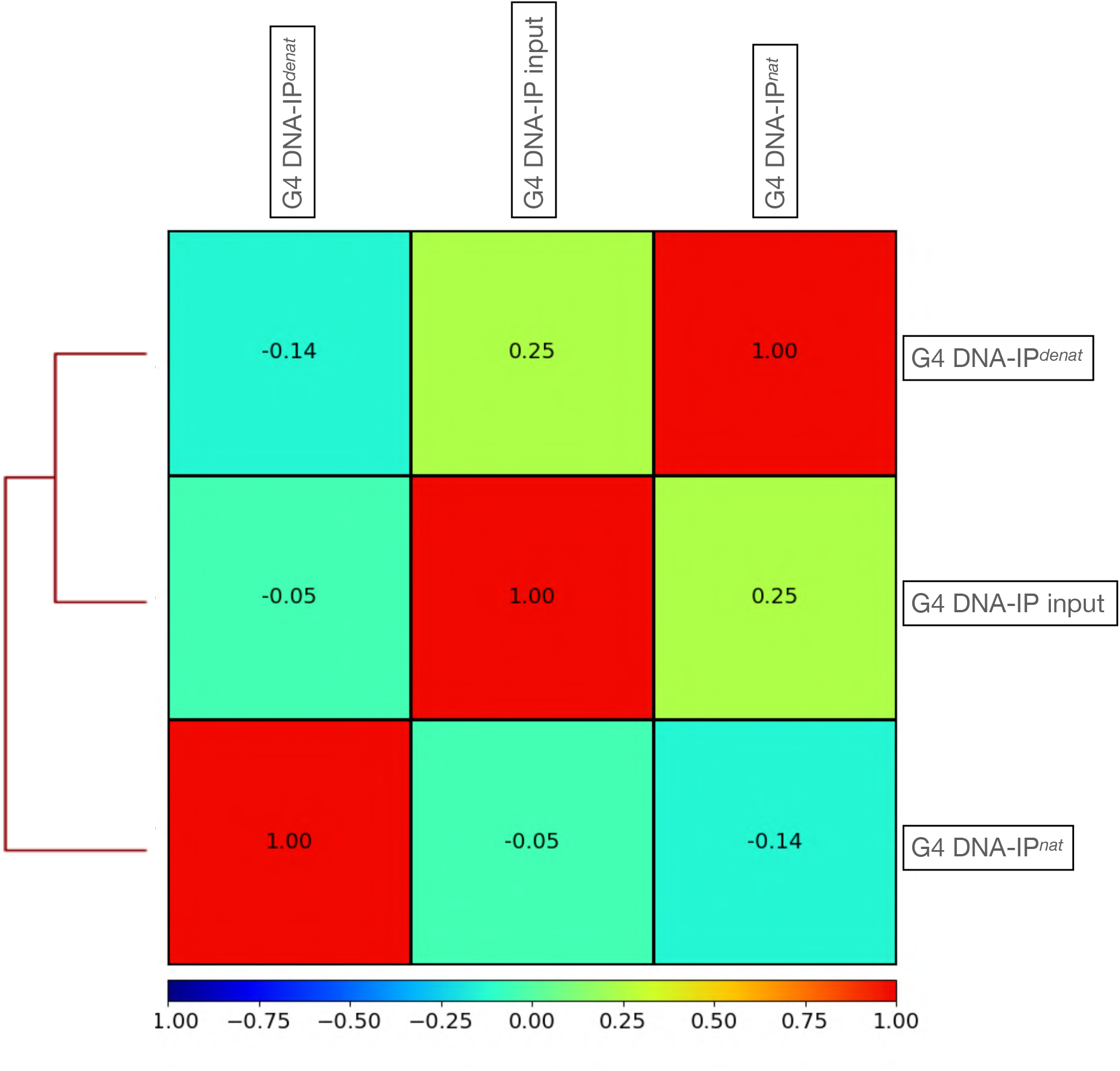
Correlation matrix and clustering for the G4 landscapes reported by various methods at signal and peak levels: A: The absolutely DNA-IPs (G4 DNA-IP^*nat*^ and G4 DNA-IP^*denat*^) show similarities with the conventional G4-ChIP, whereas the *AbC* G4-ChIP eliminates this similarity. Notably, all ChIP samples cluster with the respective inputs except for the G4 DNA-IP^*nat*^ which co-clusters with the G4-ChIP. B: The peaks in G4 DNA-IP^*nat*^ correlate with the peaks from G4-ChIP, suggesting that the spontaneously forming G4s in G4 DNA-IP^*nat*^ are also captured in G4-ChIP. This is unlike is CT^*nuc*^-which shows a poorer similarity with G4 DNA-IP^*nat*^. G4 DNA-IP^*denat*^ did not show similarity with G4-ChIP or CT^*nuc*^. The numbers in the insets are Spearman’s coefficients.

**Fig S5:**
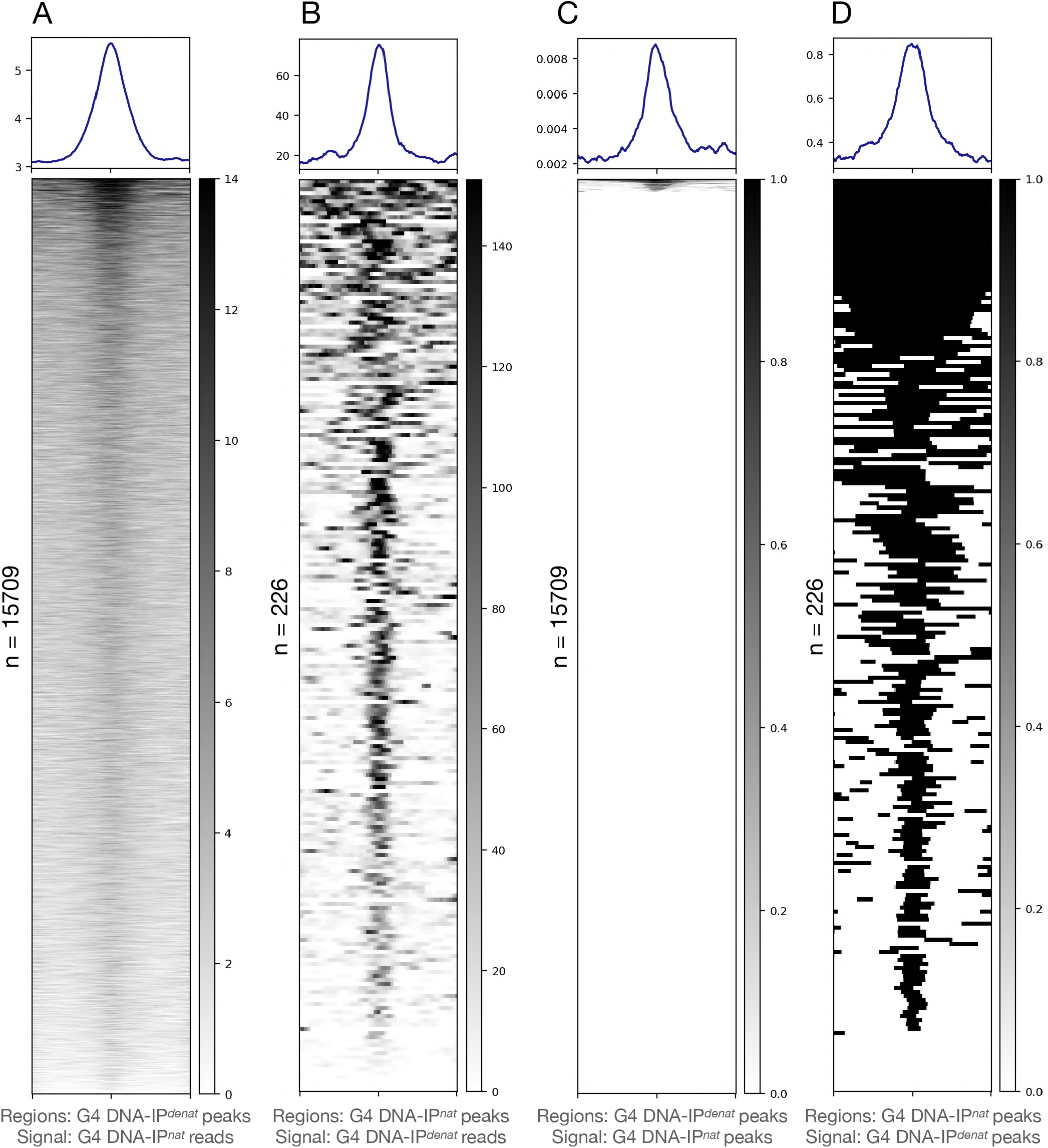
Comparison of G4 profiles of G4-ChIP with G4 DNA-IP^*nat*^ and G4 DNA-IP^*denat*^: A: G4 DNA-IP^*nat*^ peaks show a clear central enrichment of G4-ChIP signals. B: Only a very small fraction of G4 DNA-IP^*denat*^ peaks show enrichment in G4-ChIP. Most of the G4 DNA-IP^*denat*^ peaks are selectively devoid of G4-ChIP signals at the peak centres. C and D: In agreement with the other observations described so far, a larger fraction of G4-ChIP peaks contain G4 DNA-IP^*nat*^ signals, whereas (C) a much smaller fraction of G4-ChIP peaks contain G4 DNA-IP^*denat*^ signals (D). The signals were plotted on the 5 kb flanks from the centers of peaks. Numbers indicate the peaks identified in the respective sample, and the Y axes of the plots have signals plotted as mean±SEM.

**Fig S6.**
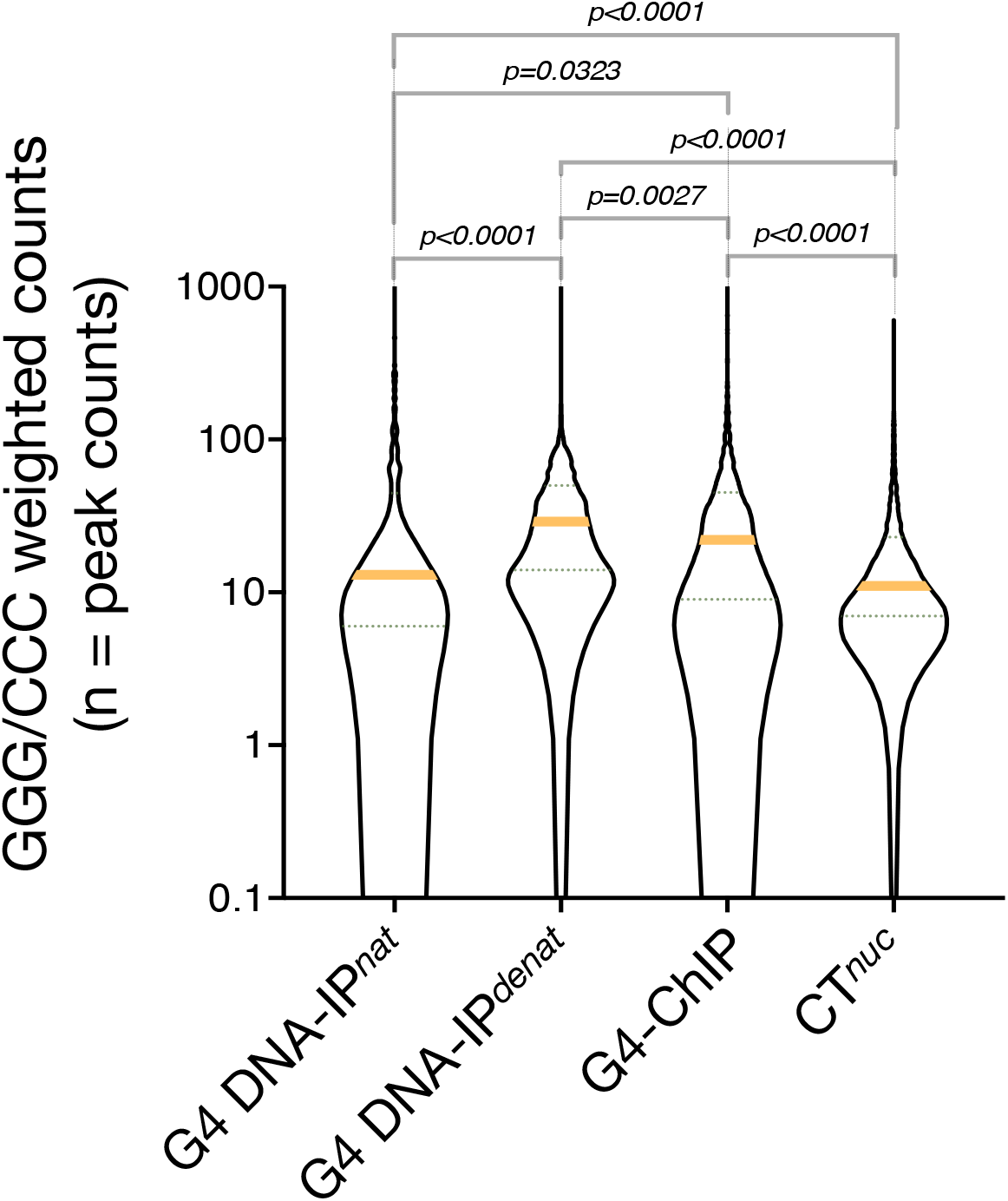
A comparison of G4-ChIP and CT^*nuc*^: At CT^*nuc*^ peaks, there is a weaker enrichment of G4-ChIP signals as compared to the G4-ChIP peaks, which show a much stronger signal with a high background of CT^*nuc*^ signals. As expected, a subset of CT^*nuc*^ peaks showed no signal from G4-ChIP, whereas all G4-ChIP peaks showed CT^*nuc*^ signals suggesting that the CT^*nuc*^ G4 profile is a superset of the G4-ChIP profile. The signals were plotted on the 5 kb flanks from the centers of peaks. Numbers indicate the peaks identified in the respective sample, and the Y axes of the plots have signals plotted as mean±SEM.

**Fig S7.**
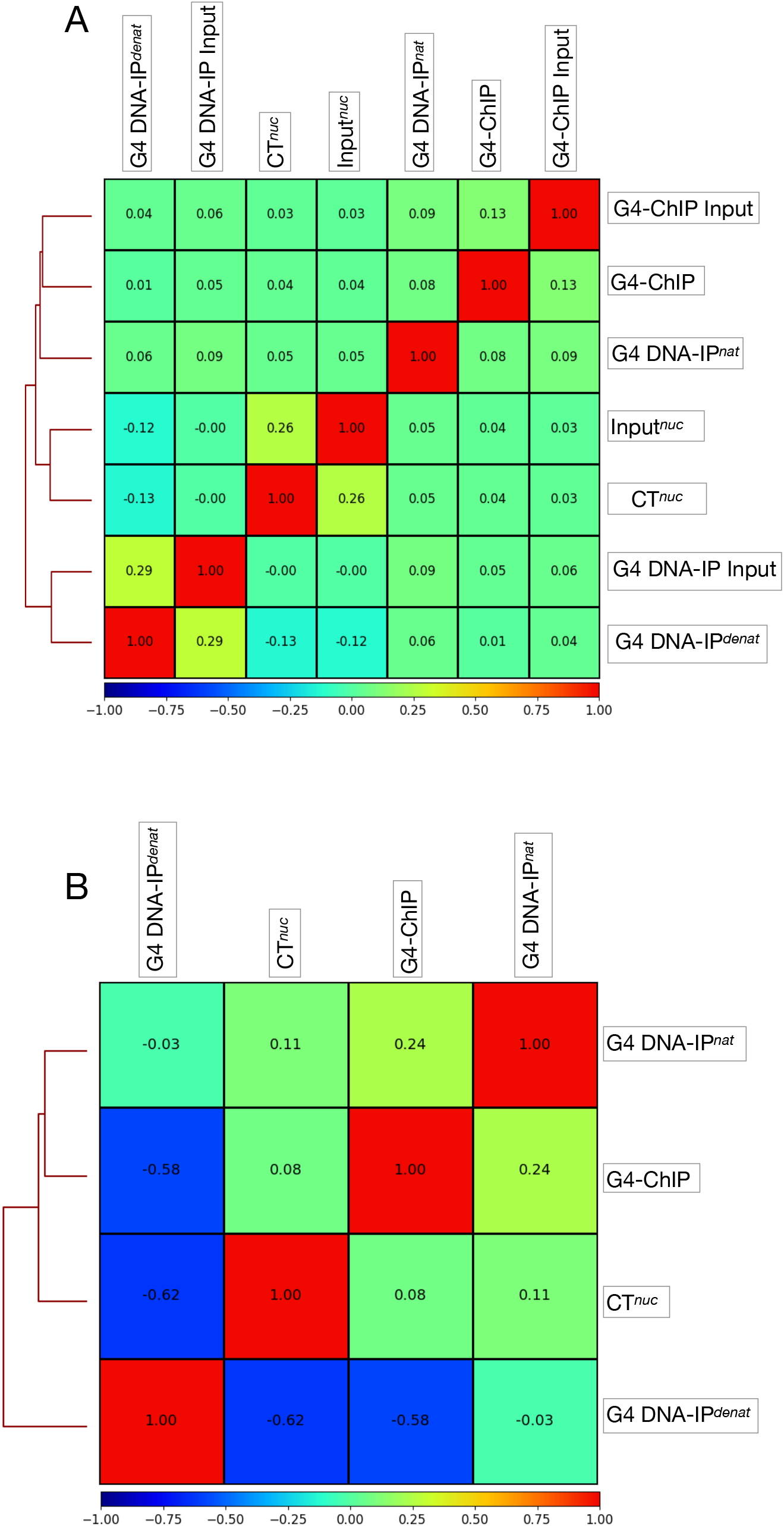
A comparison of signals from peaks of DNA-IPs and G4-ChIP on CT^*nuc*^ peaks: A and B: A very small population of CT^*nuc*^ peaks contain signals from G4 DNA-IP^*denat*^ peaks with no clear enrichment at the CT^*nuc*^ peak centers (A) while about half of the CT^*nuc*^ peaks contained signals with a lot of background from G4 DNA-IP^*denat*^ peaks (B) suggesting that peaks identified in CT^*nuc*^ (table S2) is a superset of G4 DNA-IP^*denat*^ peaks. C: The peak signals from G4-ChIP peaks showed a weaker presence on the CT^*nuc*^ peaks but with a similar background signal as shown by G4 DNA-IP^*denat*^ peaks. The signals were plotted on the 5 kb flanks from the centers of peaks. Numbers indicate the peaks identified in the respective sample and the Y axes of the plots have signals plotted as mean±SEM.

**Fig S8:**
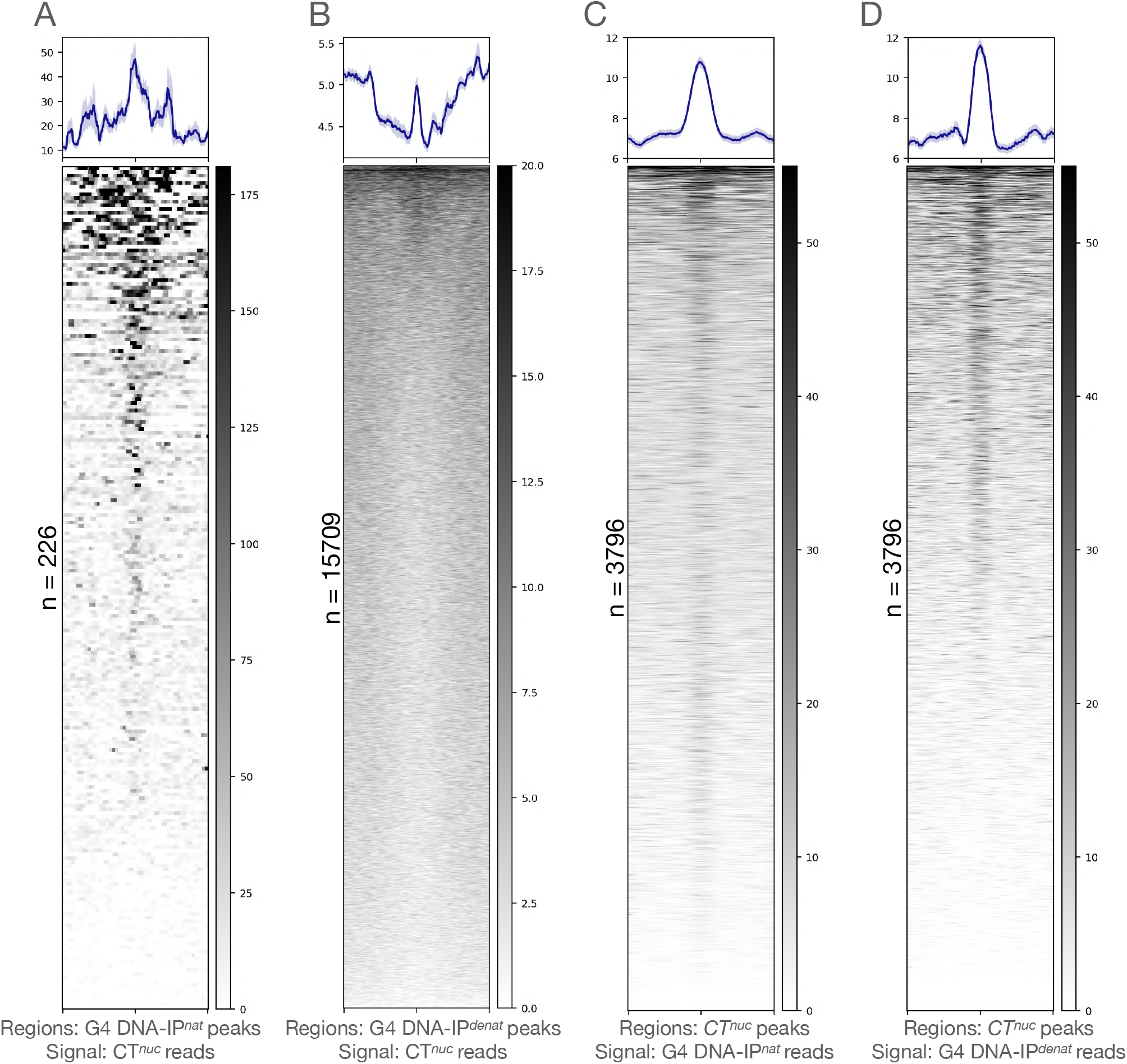
Jaccard indices of G4 peaks identified in CT^*nuc*^, G4-ChIP, G4 DNA-IP^*denat*^ and G4 DNA-IP^*nat*^. The numbers indicate the proximity of occurrences of G4 peaks genome-wide. The G4 DNA-IP^*denat*^ represents a very different profile of G4 peaks than in the rest of the samples. The profile of the G4 peaks in G4-ChIP showed a close resemblance to the G4 DNA-IP^*denat*^ and CT^*nuc*^. The lines with the bidirectional arrows indicate the pairs of samples within which the Jaccard indices were calculated. The numbers on the bottom of the lines are not on the scale.

**Fig S9:**
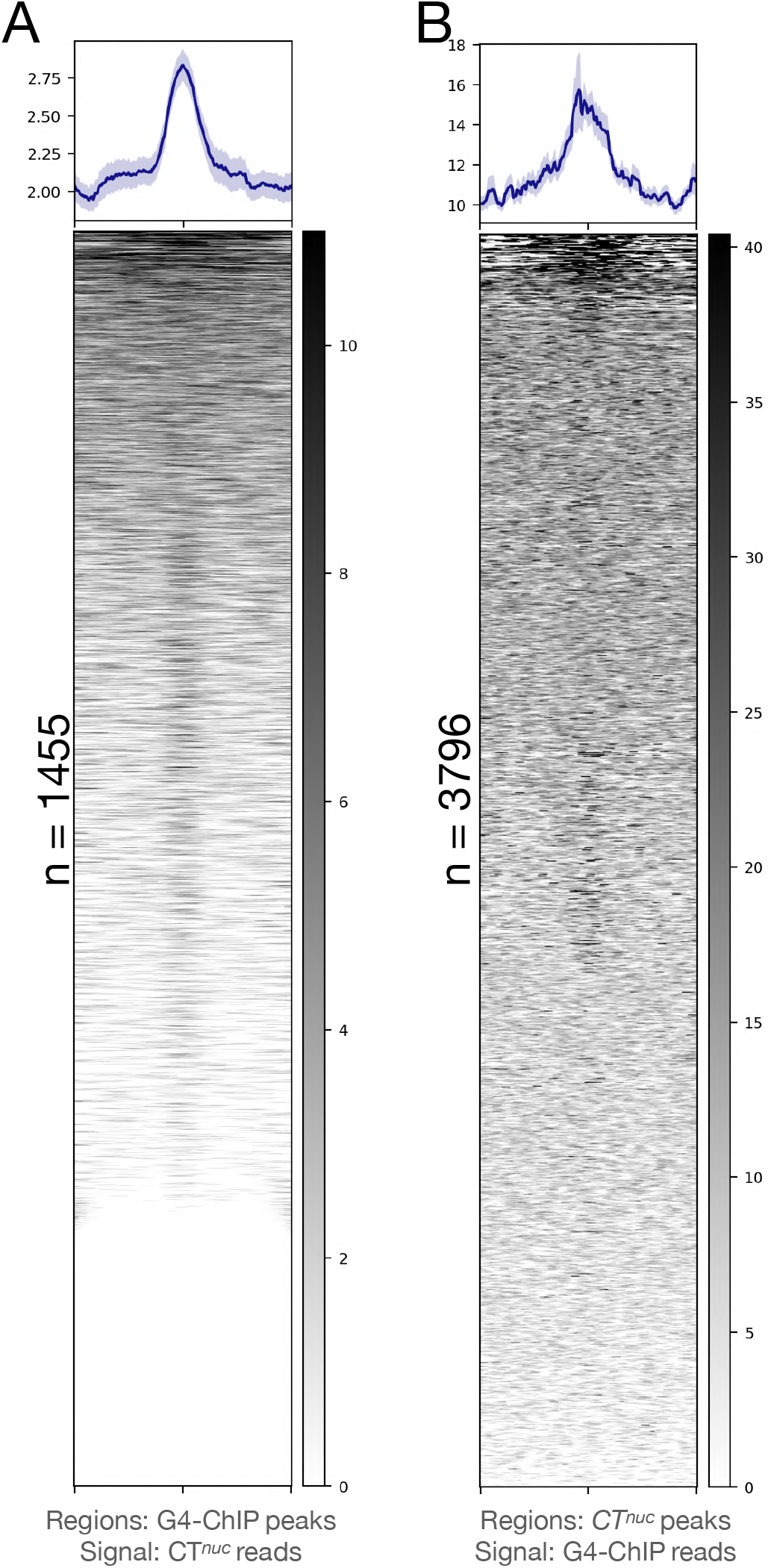
A comparison of signals from DNA-IPs, CT^*nuc*^ and G4-ChIP on randomly drawn genomic regions: A. Using 123190 (summation of 10 rounds of randomization of 0.2 kb genomic bins of 12319 regions, as in DCRs, independently from the hg38) randomly selected regions genome-wide, the G4 signals at random regions were calculated for the four samples as indicated. The CT^*nuc*^, in which *AbC* G4-ChIP was applied, showed a very restricted low level detection of G4 signals at these random regions, similar to the G4 DNA-IP^*denat*^. The G4-ChIP and G4 DNA-IP^*denat*^, on the other hand, showed a relatively high prevalence of G4 capture at random regions. These comparisons showed that the discrete and specific G4s are detected by the application of the *AbC* G4-ChIP protocol, and it minimizes *in vitro* G4s captured at random regions as artifacts. B-E. These randomly drawn 123190 genomic regions contain no or poor signal from G4 DNA-IP^*nat*^ (B), G4 DNA-IP^*denat*^ (C), G4-ChIP (D) and CT^*nuc*^ (E), respectively. The signals were plotted on the 5 kb flanks from the centers of the randomized regions. The Y axes of the plots have signals plotted as mean±SEM.

**Fig S10:**
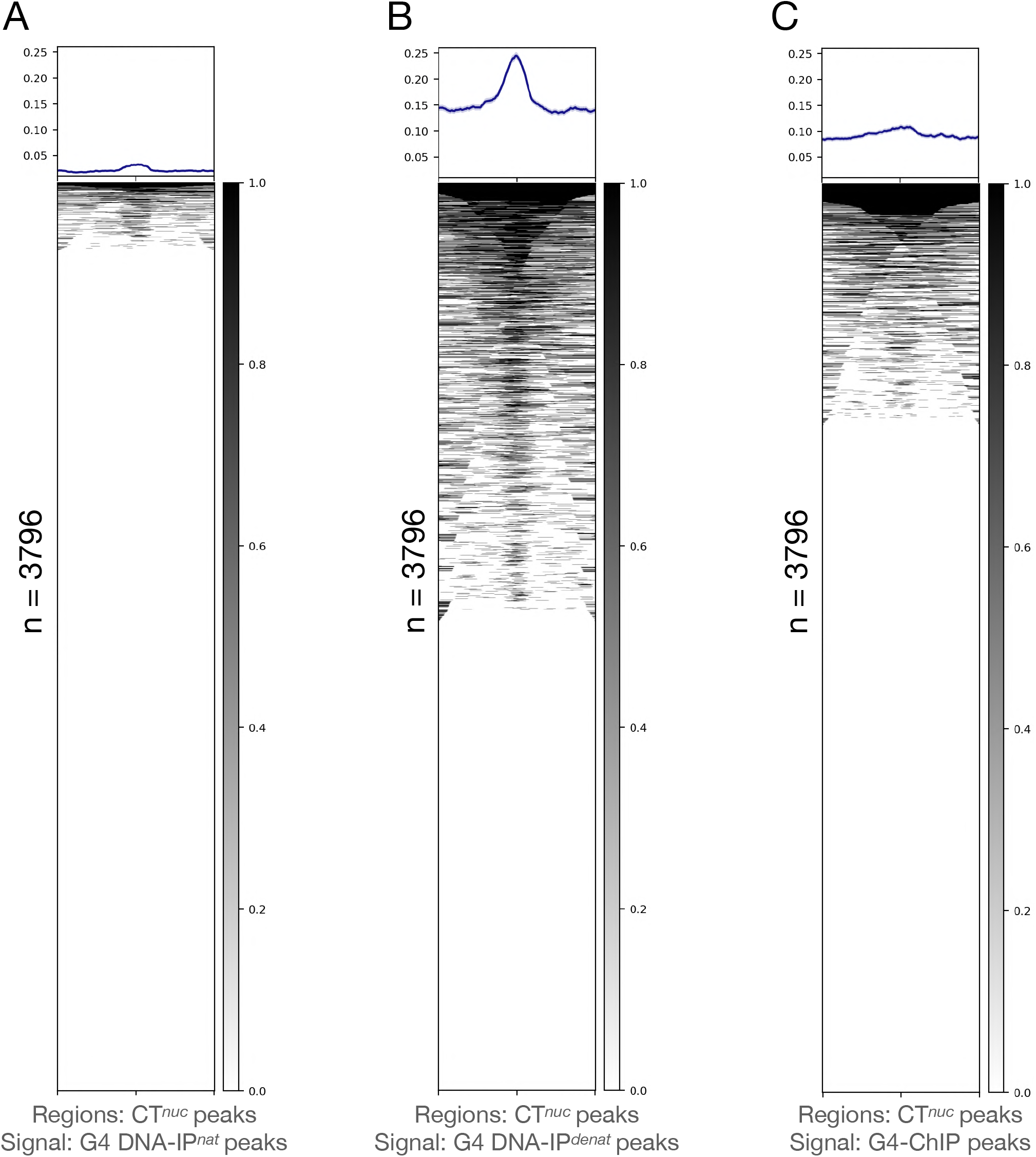
A comparison of publicly available G4-ChIP signals on genome-wide randomized regions: A-D. Publicly available previously reported G4-ChIP signals (indicated by the GSE datasets mentioned at the bottom of the heatmaps) were plotted on the 5 kb flanks from the centers of genome-wide randomized regions (as in figure S9). Overall, these regions carry a higher background from the reported G4-ChIP signals as compared to G4 DNA-IPs, G4-ChIP and CT^*nuc*^. This indicates that antibody 1H6 captures a very restrictive set of G4s. The Y axes of the plots have signals plotted as mean±SEM.

**Fig S11:**
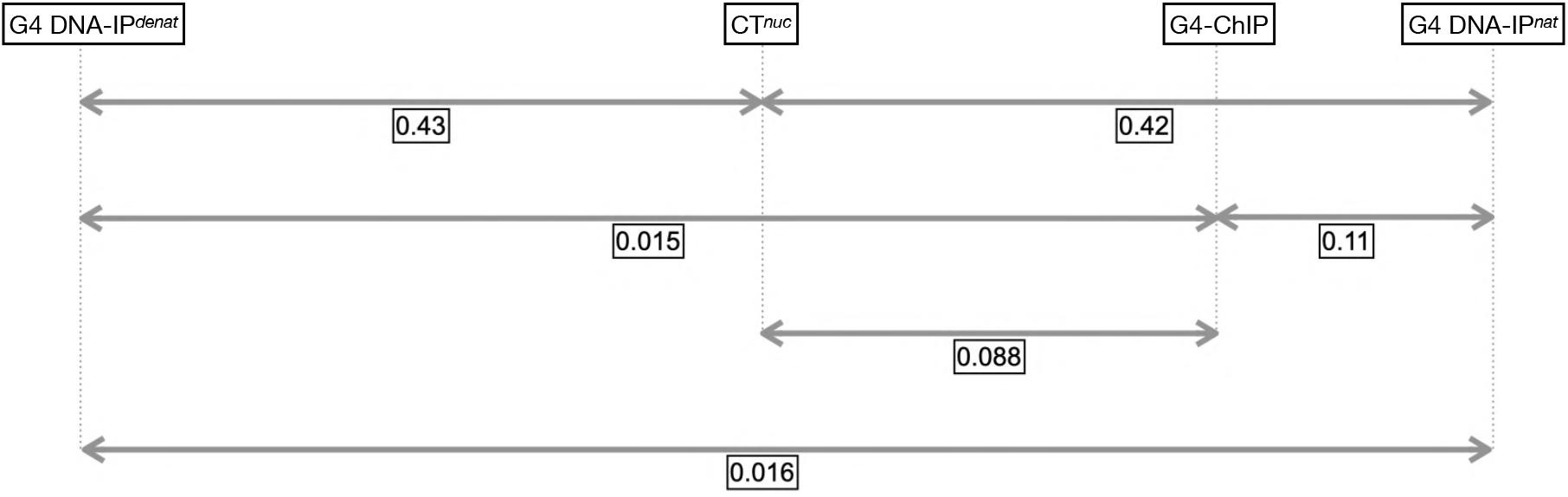
CD wavelength scan, CD thermal melt experiment and EMSA showing the preference for intramolecular hybrid G4 by the Control DNA unit sequence (5’-GGGTCTAAGGGCTCGAGGGTCTGCGGG-3’). A: CD spectra showing the preference for the hybrid G4 conformation by the DG in the presence of 50 mM Tris-HCl (pH 7.4) buffer irrespective of different KCl concentrations (0.05 M, 0.1 M, 0.15 M, 0.5 M and 1 M). Note the absence of such a conformational preference by the corresponding WC duplex (WC, colored in peach). B: The hypochromic thermal melting pattern seen for the DG at different KCl concentrations (i) 0.05 M, (ii) 0.1 M, (iii) 0.15 M, (iv) 0.5 M and (v) 1 M indicates the stable G4 formation. Note that the CD thermal melting was measured at 260 nm. C: EMSA corresponding to the DG indicates the intramolecular G4 formation at different KCl concentrations as it moves faster (Lanes 2-6) than the control d(T)_27_ (Lane 1). D: Possible intramolecular hybrid G4 conformation preferred by the DG, which is modeled using the 3D-NuS web server. The guanines of the G-quartet are colored green, and the loop residues are shown in pink.

**Fig S12:**
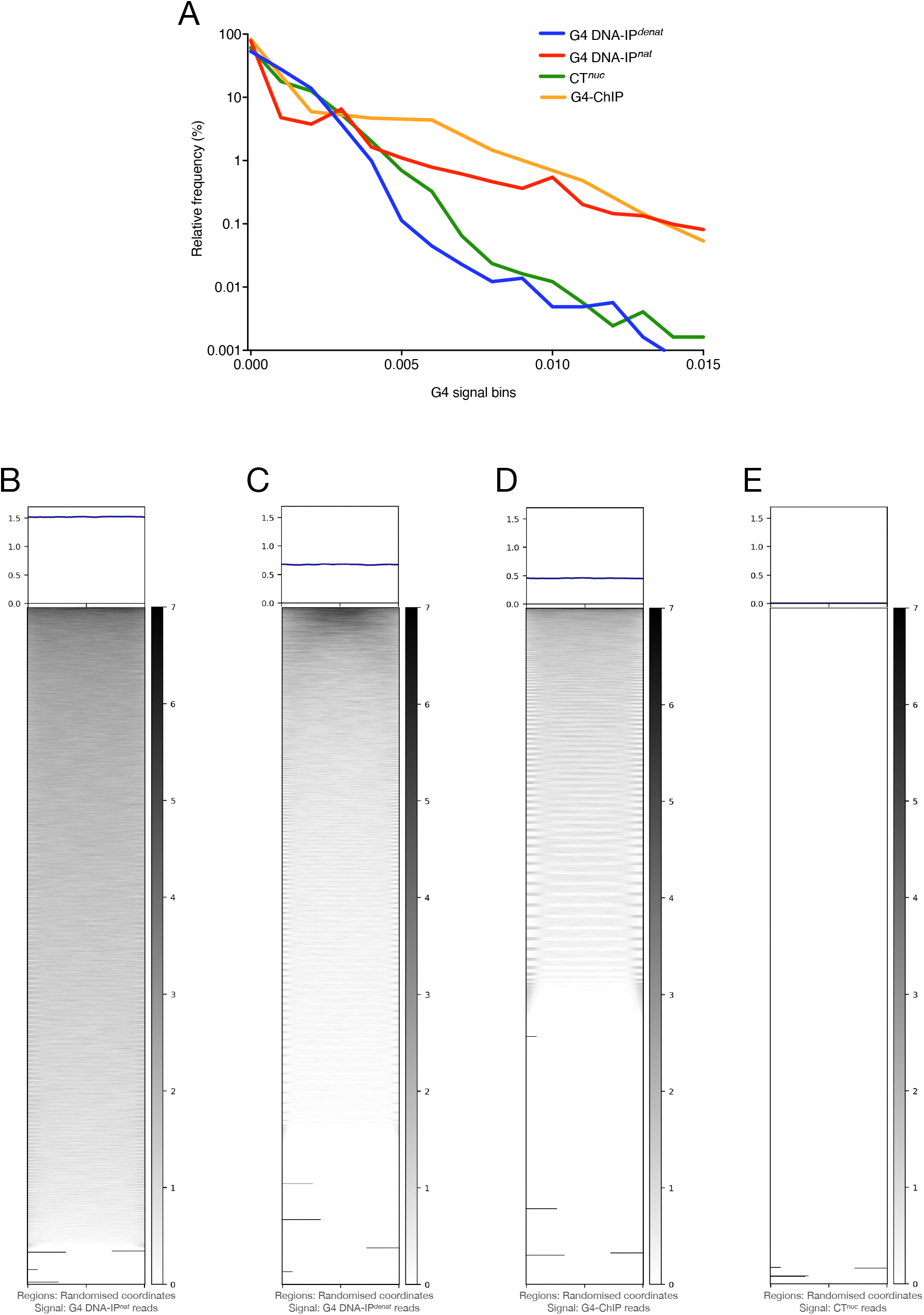
GGG/CCC content on CGGBP1 peak datasets (GSE53571). GGG/CCC content calculated in two different CGGBP1 peak datasets (growth-stimulated and starved) shows that the CGGBP1 occupancy is facilitated in genomic regions with higher GGG/CCC content upon growth stimulation. The difference in the CGGBP1 occupancy in these two datasets is denoted by a strong p-value. The orange lines indicate the median values presented by the Y-axis on a log10 scale.

**Fig S13:**
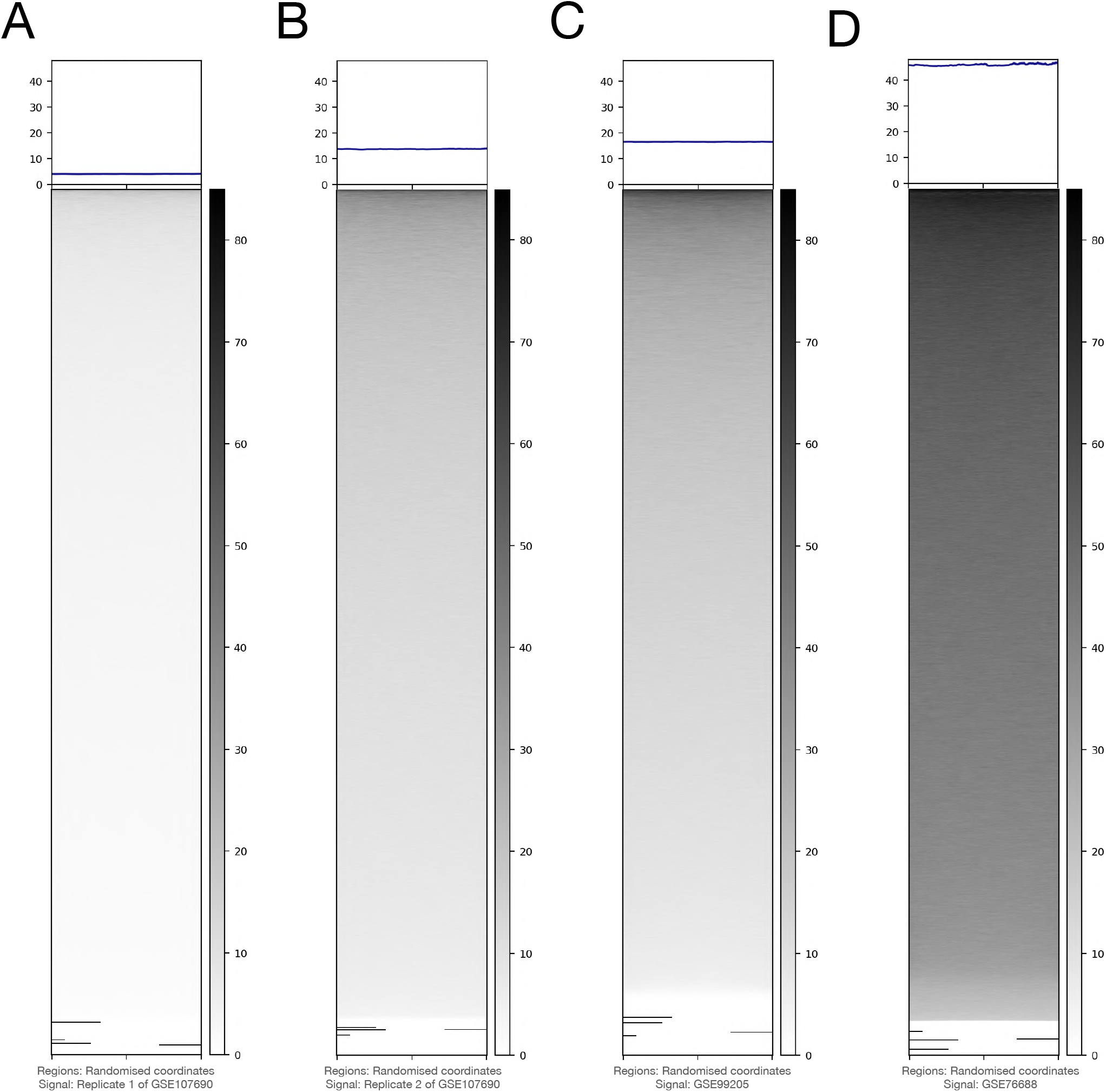
Western blot showing the knockdown of endogenous levels of CGGBP1. The levels of CGGBP1 and GAPDH are shown in the lower and upper panels, respectively. The level of knockdown of CGGBP1 is normalized to the levels of GAPDH. Approximately 99% knockdown of endogenous levels of CGGBP1 was observed in KD.

**Fig S14:**
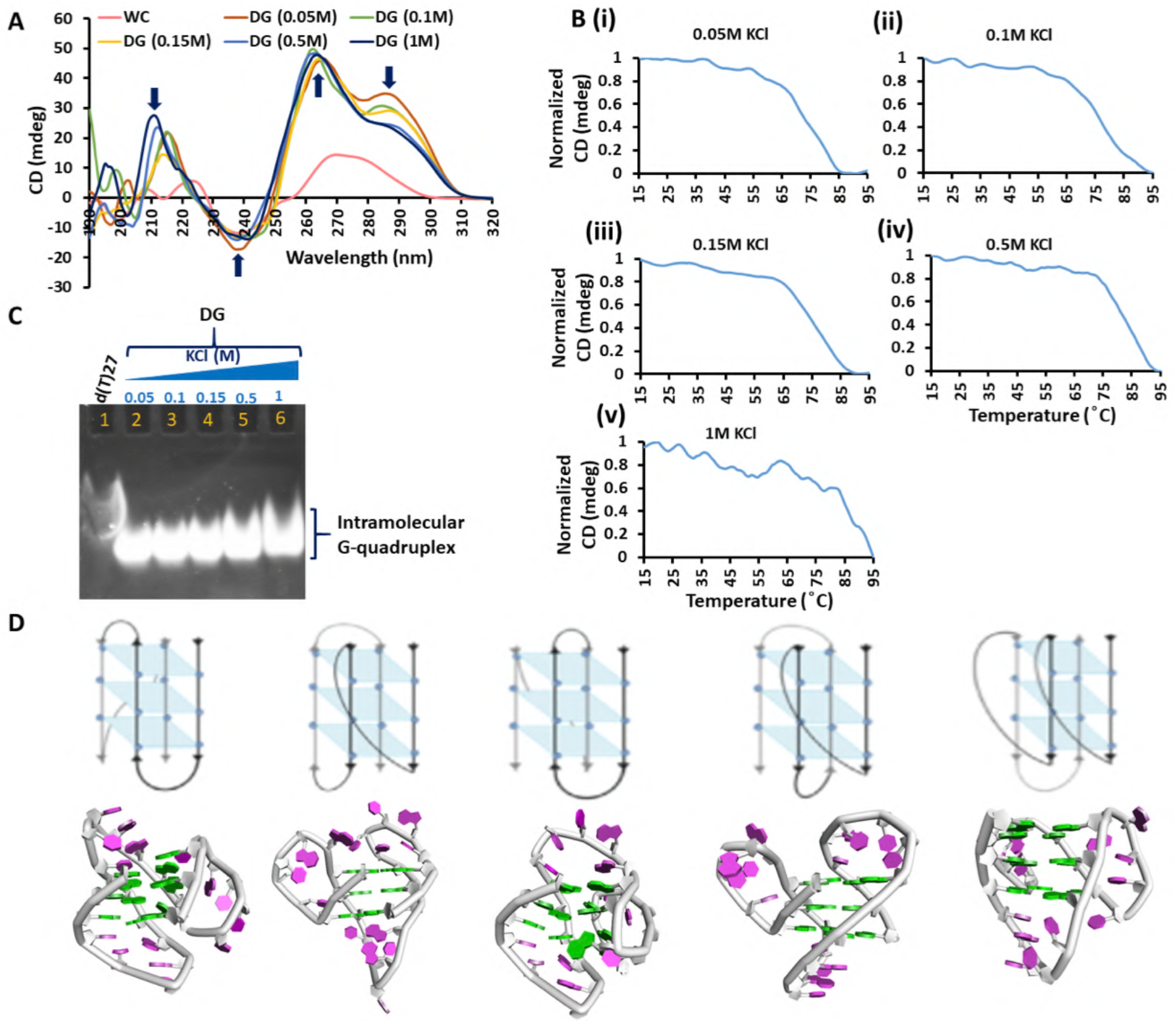
Recovery of control and carrier DNA in the inputs of CT^*ut*^, KD^*ut*^, CT^*tr*^ and KD^*tr*^. The qPCR quantification confirms the bias-free presence of the control and control DNA between CT and KD as quantified in their input samples. Notably, the *in vitro* addition of the control and carrier DNA in the untransfected samples yielded a higher amount of the control DNA than that can be quantified in the transfected samples. The Y-axis represents the concentration of the control and control DNA on a log10 scale.

**Fig S15:**
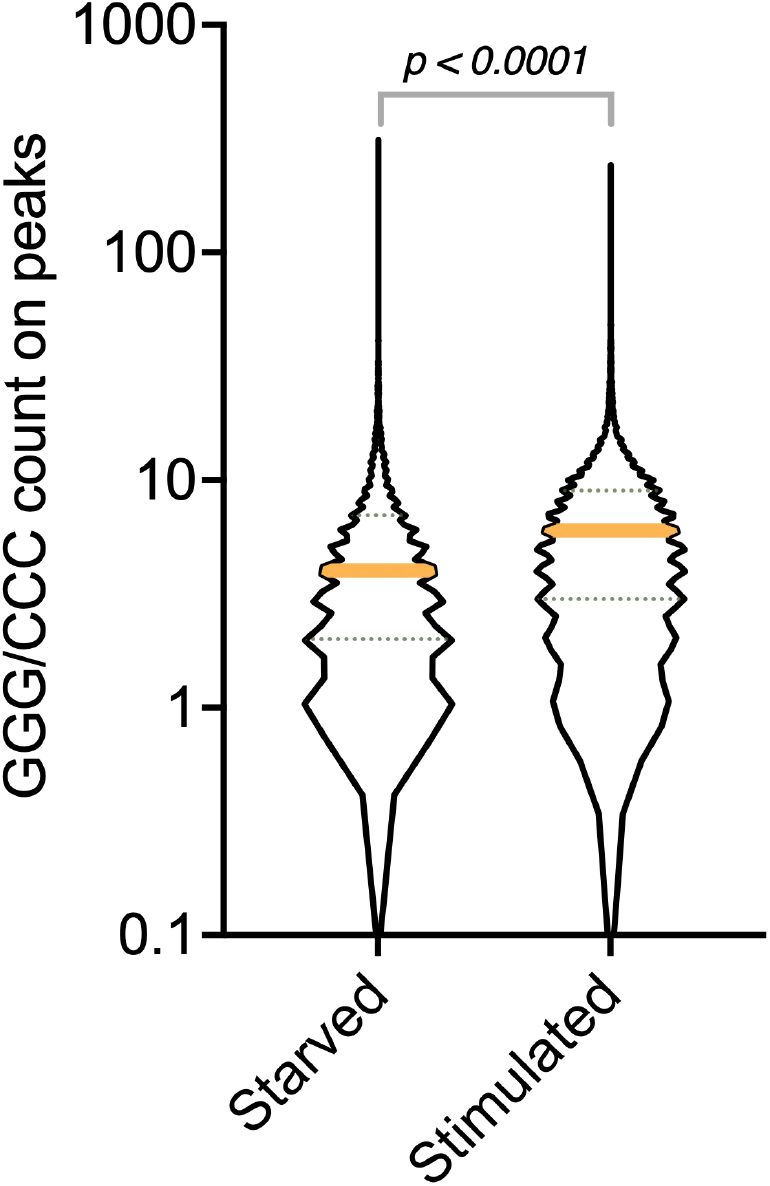
A comparison of *AbC* G4-ChIP signals reciprocally on the peaks identified in CT and KD samples: A-F: The *AbC* G4-ChIP signals were plotted reciprocally on peaks of CT samples and in the same order in the KD samples (G-L). The signals were plotted on the 5 kb flanks from centers of the peaks. The peaks identified in each sample of CT and KD show that other replicates also contain signals but weakly. Numbers indicate the peaks identified in the respective sample and the Y axes of the plots have signals plotted as mean±SEM.

**Fig S16:**
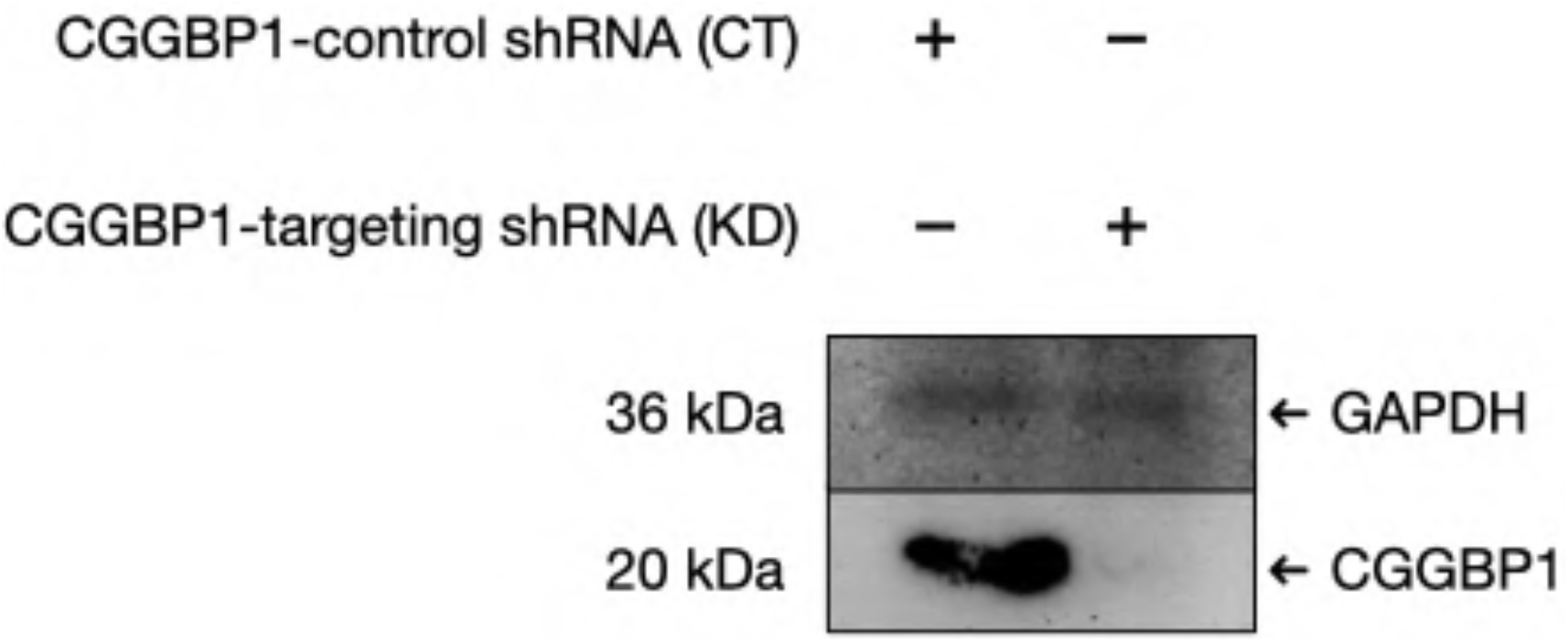
A volcano plot of 0.2 kb genomic bins compared pairwise in triplicates between CT and KD. The X-axis represents log2 fold change calculated as KD/CT, and the corresponding p-values on a -log10 scale are plotted on the Y-axis. The violin plots represent the density of the data points along the two axes. The red-green partition marks the non-DCRs and DCRs, respectively. The green data points correspond to the orange data points in fig 4A.

**Fig S17:**
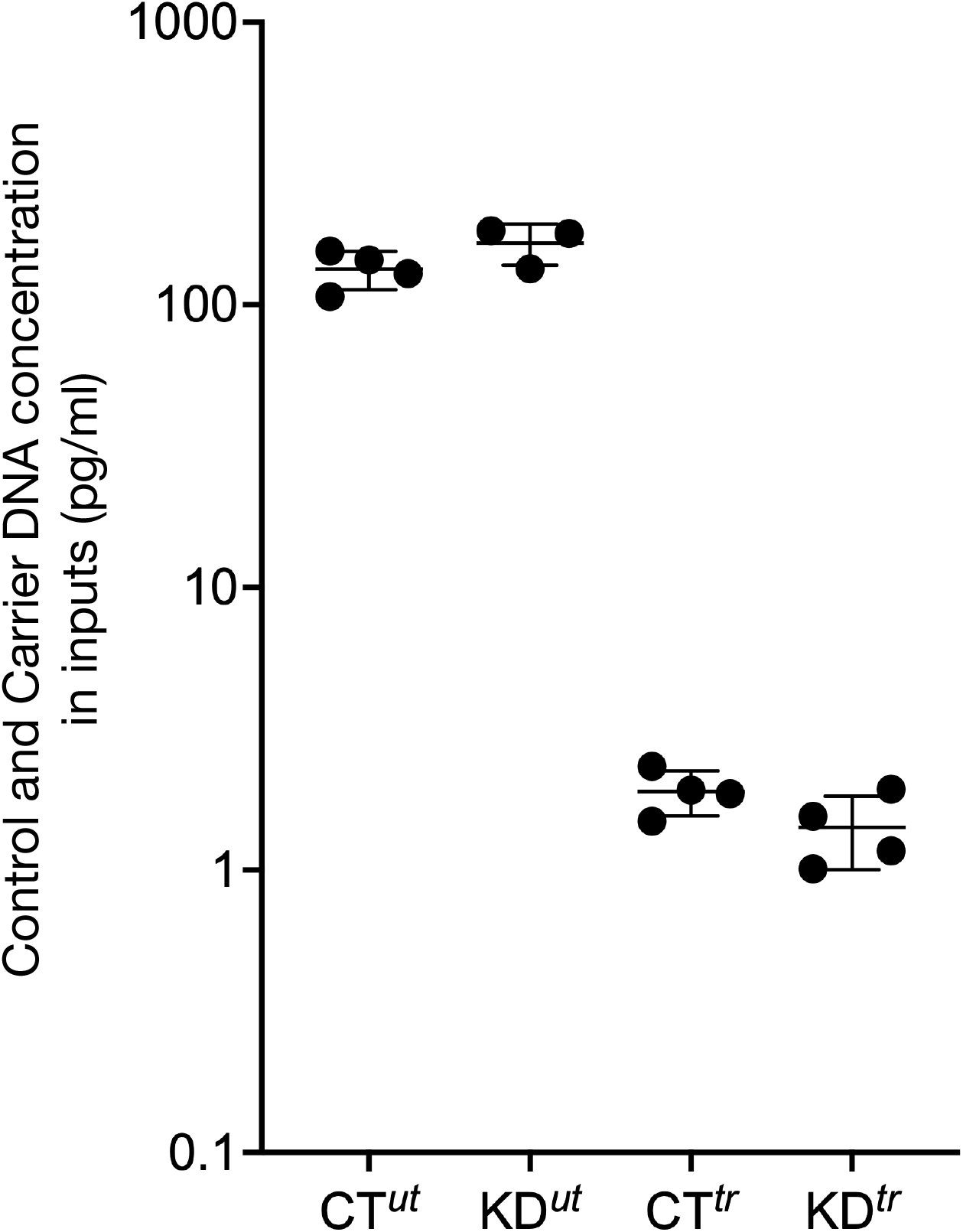
Functional analysis of DCRs: A. By using GREAT (web-based tool http://great.stanford.edu/) with an association condition of −5 kb to +1 kb from TSS for *cis* association and 10 kb in any direction for *trans* association, we found that nearly 90% of the DCRs are not associated with genes (red bar). B and C: By plotting the occurrence of known permissive enhancers (FANTOM5) against the bed coordinates of DCRs, we found that the DCRs have no association with enhancers, with about 75% of DCRs having no known enhancers in 5 kb flanks (B). A similar analysis of the occurrence of CpG islands against the bed coordinates of DCRs showed that the DCRs are preferentially located in CpG island-poor regions. D. The DCRs are also distant from the known origin of replication sites annotated by the mentioned markers (GSE28911, GSE37583 and GSE81165). Numbers indicate the regions identified in the respective sample, and the Y axes of the plots have signals plotted as mean±SEM.

**Fig S18:**
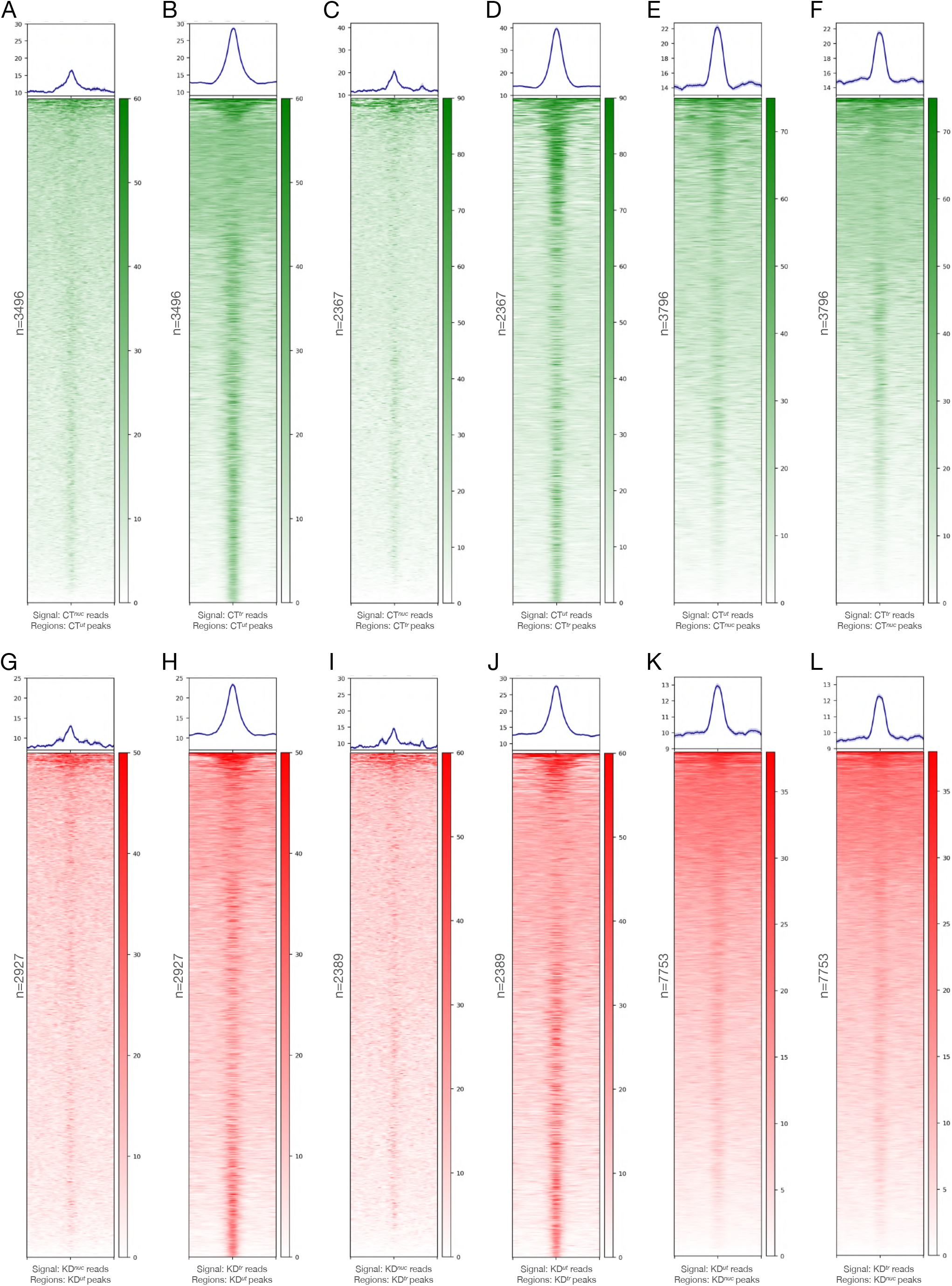
Discovery of signals from publicly available G4-ChIP datasets: A - D: The heatmaps showed the weak enrichment of G4 signals from independent experiments on the 5 kb flanks of the DCRs. This suggests that the DCRs identified from the current study are genuine G4-forming regions. Numbers indicate the regions identified in the respective sample, and the Y axes of the plots have signals plotted as mean±SEM. The signals have been derived from the GSE datasets mentioned below the heatmaps.

**Fig S19:**
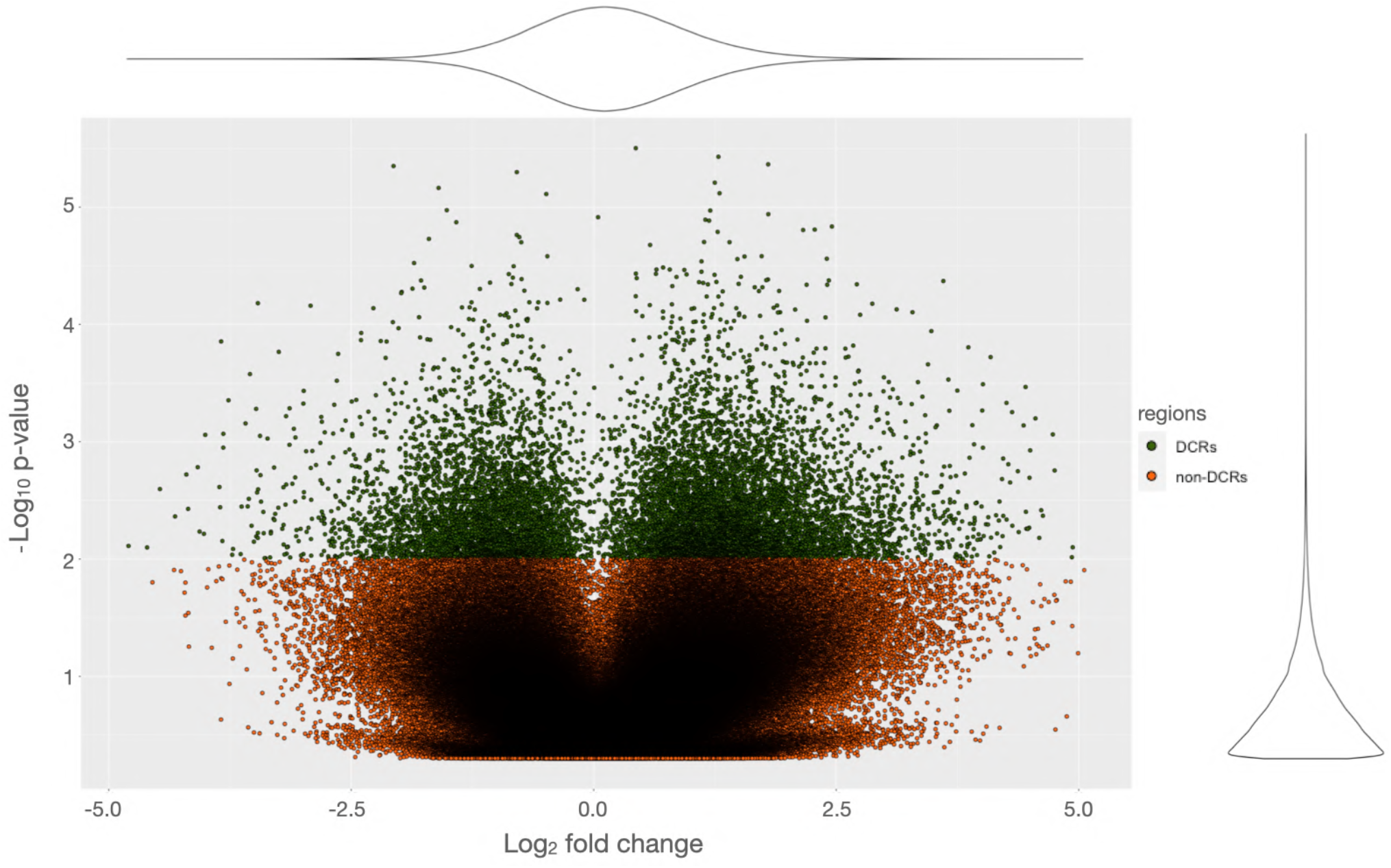
Quantitative analysis of G4 formation on the Control DNA: A. The Control DNA formed stable G4s in the presence of KCl and thus gave poor amplification in the polymerase stop assay compared to the sample of the Control DNA in the presence of LiCl. B and C. The Control DNA was pre-incubated with rCGGBP1 first and then either incubated in the presence of KCl (B) or LiCl (C). A stronger polymerase stop was observed in the presence of LiCl (C) than KCl (B), which suggests that rCGGBP1 occupies the Control DNA, perhaps in the absence of G4s. D. Western blot assay of immunoprecipitation of CGGBP1 from HEK293T cells under low salt condition. E. Pre-incubation of the Control DNA with immunoprecipitated CGGBP1 yielded a similar effect of polymerase stop. A non-specific isotype immunoprecipitate was used as a negative control. In the presence of LiCl, where G4-formation is less favoured, the polymerase stop showed a stronger effect when preincubated with immunoprecipitated CGGBP1 compared to KCl, wherein immunoprecipitated CGGBP1 did not yield a significant polymerase stop effect. The shaded circles indicate the replicates for respective samples, while the asterisk represents an order of significance level.

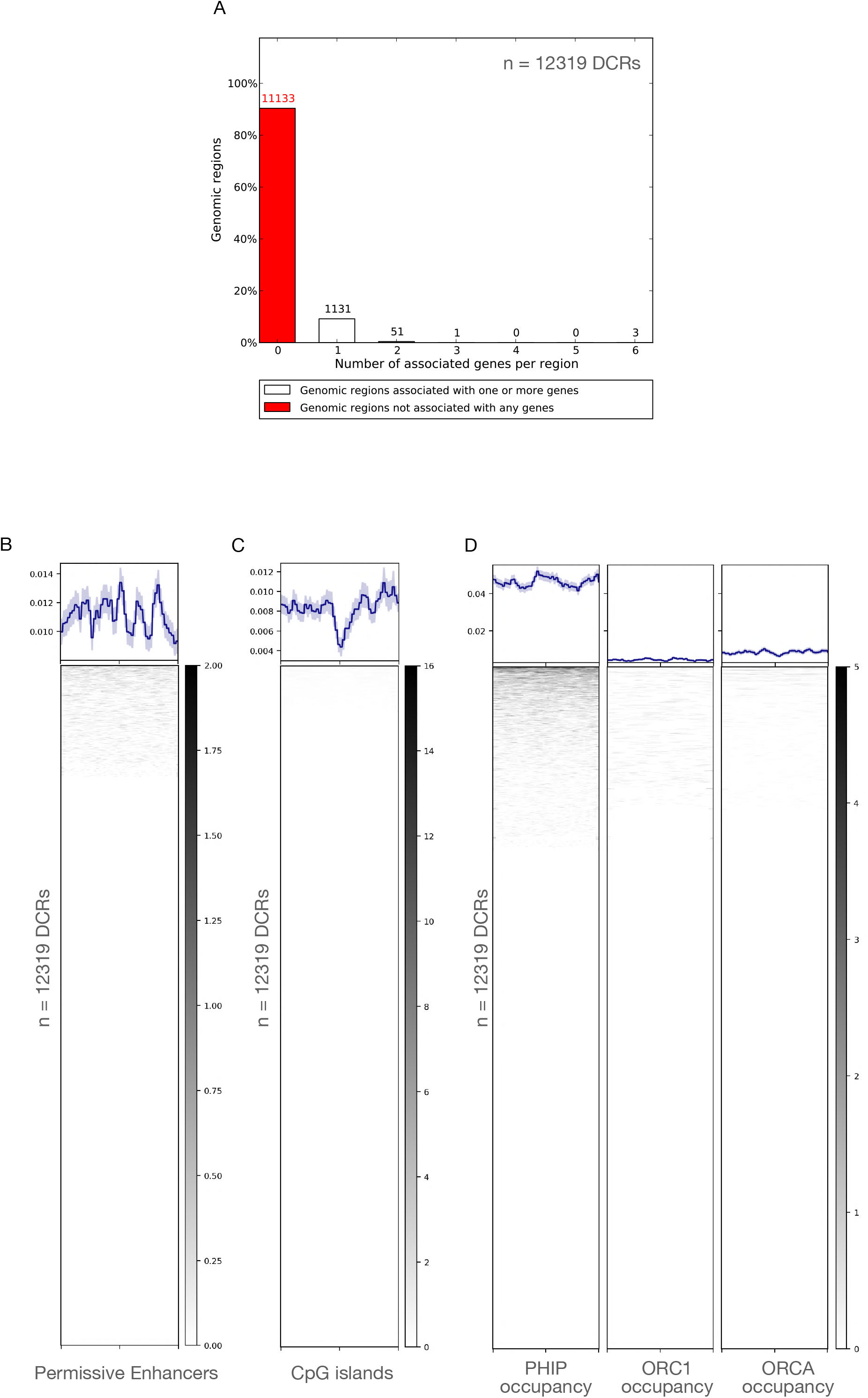

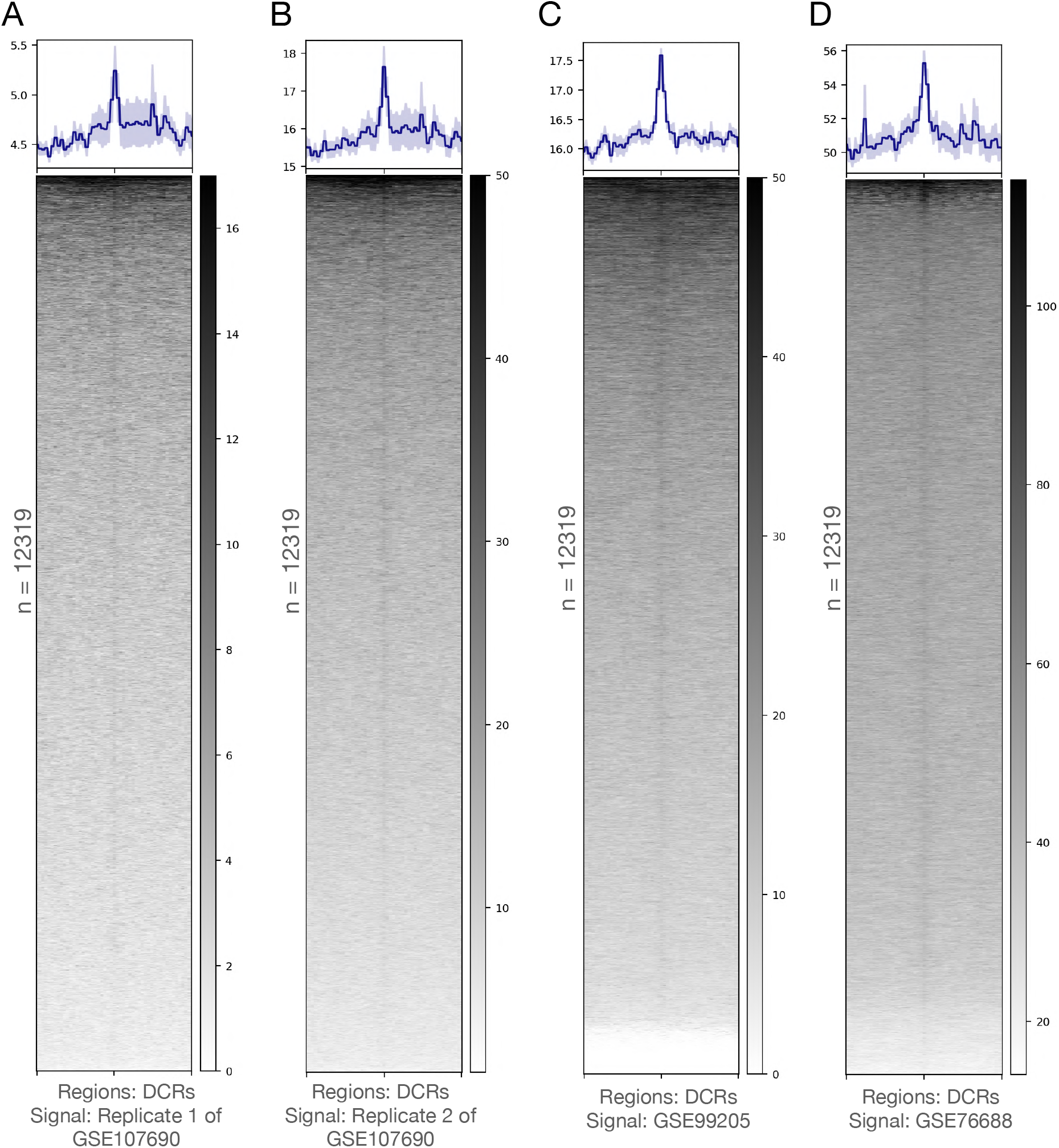

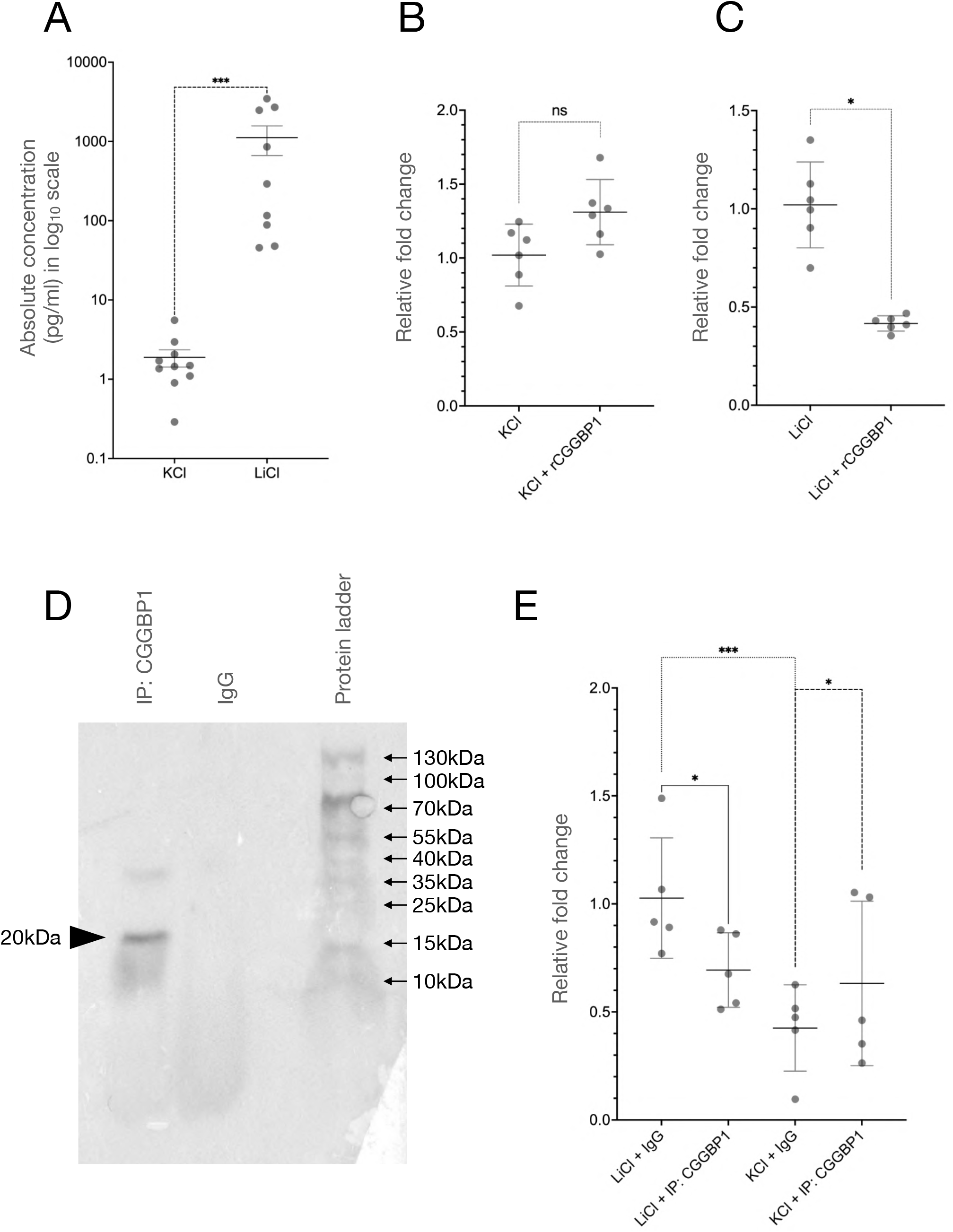

**Table S1:** Summary of sequencing data acquired for *in vitro* G4 DNA-IPs, G4-ChIP and the corresponding inputs.

| Samples | Total reads sequenced | Reads run through Porechop | Mean length of reads run through Porechop | % mapped reads run through Porechop |
| --- | --- | --- | --- | --- |
| **G4 DNA-IP<sup>nat</sup>** | 9972207 | 9966894 | 1095.6 | 62.26 |
| **G4 DNA-IP<sup>denat</sup>** | 13487569 | 13401504 | 667.29 | 54.40 |
| **G4-ChIP** | 7089256 | 7083791 | 1040.75 | 34.42 |
| **G4 DNA-IP Input** | 5845120 | 5809383 | 605.19 | 63.29 |
| **G4-ChIP Input** | 5570204 | 5558589 | 829.10 | 56.05 |

**Table S2:** Properties of peaks called on aligned sequences in the samples *in vitro* G4 DNA-IPs, G4-ChIP and CT^*nuc*^.

| Samples | Total peak count | Total peak length (bp) | Average peak length (bp) |
| --- | --- | --- | --- |
| **G4 DNA-IP<sup>nat</sup>** | 226 | 676935 | 2995.29 |
| **G4 DNA-IP<sup>denat</sup>** | 15709 | 33399525 | 2126.14 |
| **G4-ChIP** | 1455 | 3715550 | 2553.64 |
| **CT<sup>nuc</sup>** | 3796 | 5315403 | 1400.26 |

**Table S3:** Summary of *allquads* output on the sequences of *in vitro* G4-DNA-IPs, G4-ChIP, CT^*nuc*^ and the corresponding inputs.

| Samples | Reads subjected to *allquads* | Reads with G4 signature sequence identified by *allquads* | % Reads with G4 signature sequence identified by *allquads* |
| --- | --- | --- | --- |
| G4 DNA-IP<sup>nat</sup> | 9966894 | 1214180 | 12.18 |
| G4 DNA-IP<sup>denat</sup> | 13401504 | 820014 | 6.12 |
| G4-ChIP | 7083791 | 640838 | 9.05 |
| CT<sup>nuc</sup> | 45882202 | 1197364 | 2.61 |
| G4 DNA-IP Input | 5809383 | 341810 | 5.88 |
| G4-ChIP Input | 5558589 | 339856 | 6.11 |
| AbC G4-ChIP Input | 40114795 | 1020311 | 2.54 |

**Table S4:** Summary of *allquads* output on peaks called in *In vitro* G4 DNA-IP, G4-ChIP and CT^*nuc*^.

| Samples | Unique peaks with G4 | Peak length with G4 signature (bp) | % peaks having G4 signature | % of peak lengths containing G4 signature |
| --- | --- | --- | --- | --- |
| G4 DNA-IP<sup>nat</sup> | 41 | 10135 | 18.14 | 1.50 |
| G4 DNA-IP<sup>denat</sup> | 3598 | 139154 | 22.90 | 0.42 |
| G4-ChIP | 189 | 16203 | 12.99 | 0.44 |
| CT<sup>nuc</sup> | 380 | 35422 | 10.01 | 0.67 |

**Table 5:** Summary of *allquads* output on previously published data (GSE99205 and GSE76688).

| Samples | Reads trimmed for sequencing adapters | Reads with G4 signature sequence identified by *allquads* | % reads with G4 signature sequence identified by *allquads* |
| --- | --- | --- | --- |
| HaCaT_Replicate1 | 35539225 | 992034 | 2.79 |
| HaCaT_Replicate2 | 46171793 | 1408214 | 3.05 |
| HaCaT_Replicate3 | 134148412 | 3684342 | 2.75 |
| HaCaT_Input | 55700688 | 680770 | 1.22 |
| NHEK_Replicate1 | 274391548 | 4647856 | 1.69 |
| NHEK_Replicate2 | 276477916 | 5251417 | 1.90 |
| NHEK_Input | 179603728 | 2663305 | 1.48 |

**Table S6:** Summary of sequence data for CT^*nuc*^ and the corresponding input.

| Samples | Total reads sequenced | Total quality filtered reads (≥100 bp) | Basecount of quality filtered reads (≥100 bp) | Average length of quality filtered reads (bp) | % of quality filtered reads aligned using bowtie2 |
| --- | --- | --- | --- | --- | --- |
| CT<sup>nuc</sup> | 77630217 | 45882202 | 7530031562 | 164.12 | 89.35 |
| Input<sup>nuc</sup> | 71364574 | 40114795 | 6565787917 | 163.67 | 86.71 |

**Table S7:** Summary of sequencing data acquired for KD^*nuc*^.

| Samples | Total reads sequenced | Total quality filtered reads ($\geq 100$ bp) | Basecount of quality filtered reads ($\geq 100$ bp) | Average length of quality filtered reads (bp) | % of quality filtered reads aligned using bowtie2 |
| --- | --- | --- | --- | --- | --- |
| $KD^{nuc}$ | 77577363 | 43113118 | 6807407492 | 157.9 | 80.54 |

**Table S8:**
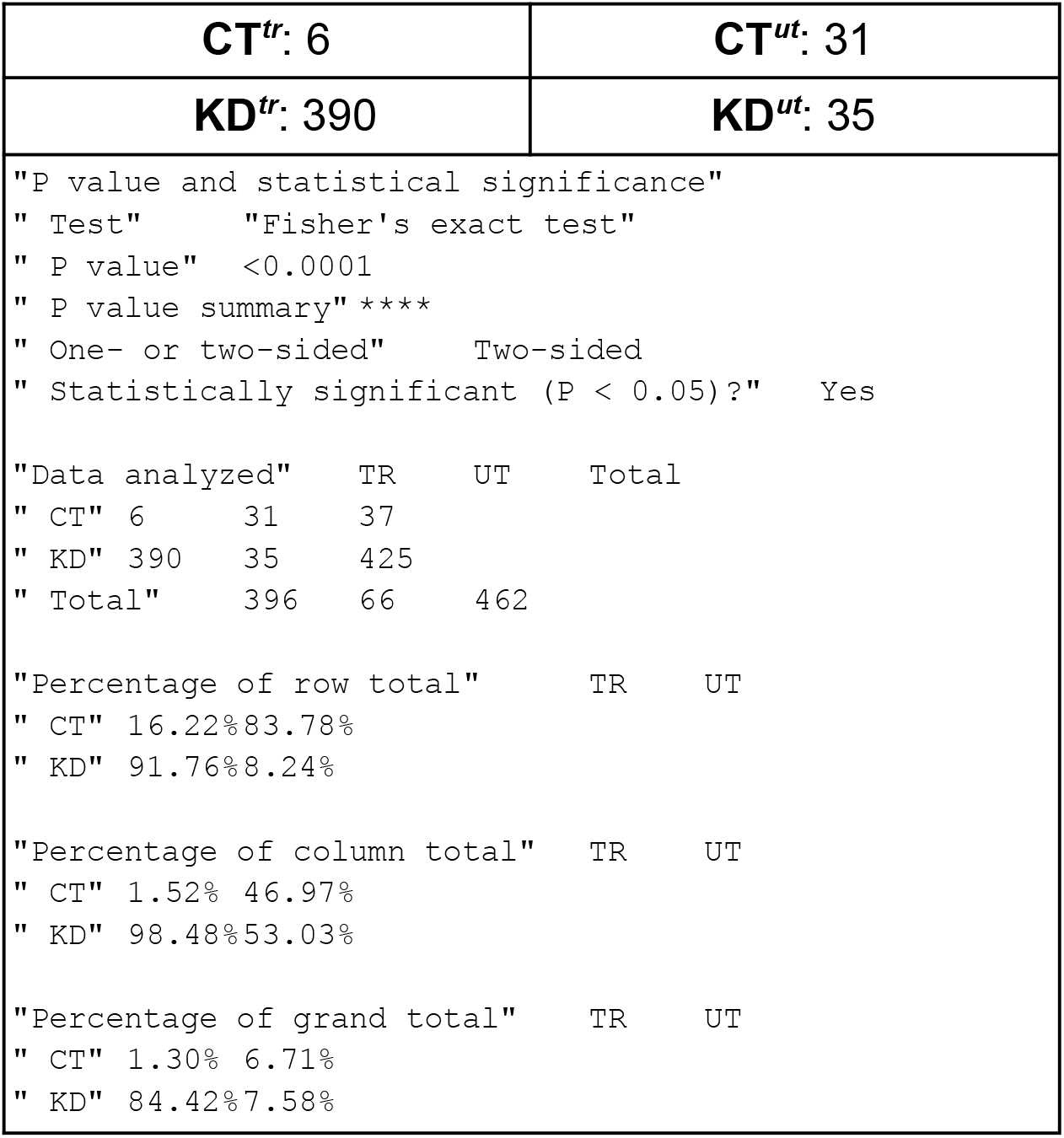
Fisher’s exact test on the number of control DNA sequences recovered from CT^*tr*^, KD^*tr*^, CT^*ut*^ and KD^*ut*^.

Table S9: Separately submitted as a spreadsheet.

**Table S10:** SINE and LINE contents of 12319 DCRs (lowest p values), non-DCRs (highest p values) and randomly drawn 0.2kb genomic regions derived using RepeatMasker.

| Repeat types | DCRs | non-DCRs | Random genomic regions |
| --- | --- | --- | --- |
| Total SINEs | 5.86 | 10.66 | 12.01 |
| ALUs | 3.22 | 7.92 | 9.78 |
| MIRs | 2.61 | 2.71 | 2.2 |
| Total LINEs | 13.65 | 16.68 | 20.01 |
| L1 | 11.05 | 13.03 | 16.98 |
| L2 | 2.1 | 3.08 | 2.59 |
| L3 | 0.29 | 0.39 | 0.3 |

**Table S11:**
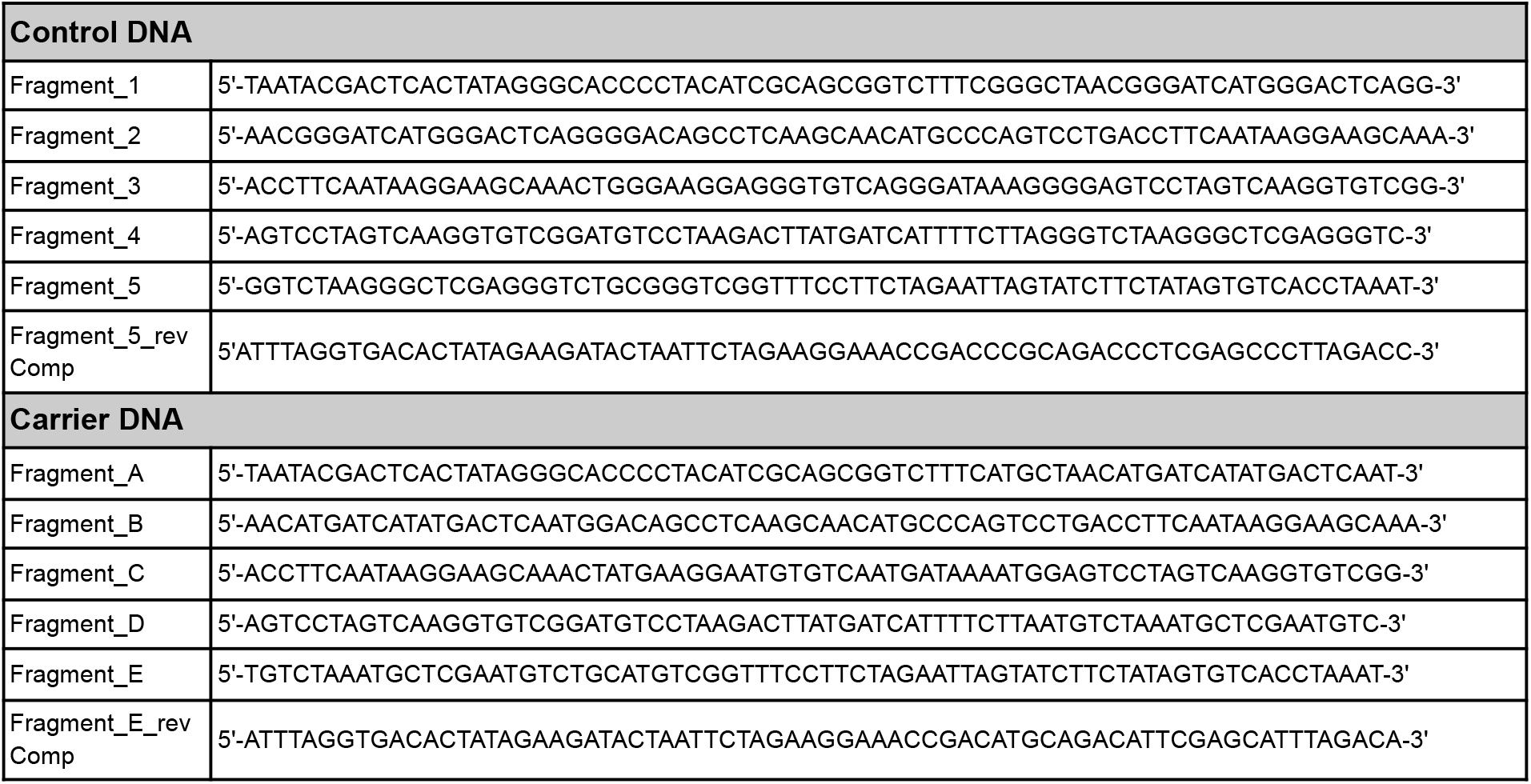
DNA oligonucleotides used for the synthesis of the Control DNA and the carrier DNA.

## REFERENCES

1. Wolffe AP, Guschin D. Review: chromatin structural features and targets that regulate transcription. J Struct Biol 2000; 129:102–22.

2. Klemm SL, Shipony Z, Greenleaf WJ. Chromatin accessibility and the regulatory epigenome [Internet]. Nature Reviews Genetics 2019; 20:207–20. Available from: http://dx.doi.org/10.1038/s41576-018-0089-8

3. Xiang J-F, Corces VG. Regulation of 3D chromatin organization by CTCF. Curr Opin Genet Dev 2021; 67:33–40.

4. Patel D, Patel M, Datta S, Singh U. CGGBP1-dependent CTCF-binding sites restrict ectopic transcription. Cell Cycle 2021; 20:2387–401.

5. Patel D, Patel M, Datta S, Singh U. CGGBP1 regulates CTCF occupancy at repeats. Epigenetics Chromatin 2019; 12:57.

6. Patel M, Patel D, Datta S, Singh U. CGGBP1-regulated cytosine methylation at CTCF-binding motifs resists stochasticity. BMC Genet 2020; 21:84.

7. Bochman ML, Paeschke K, Zakian VA. DNA secondary structures: stability and function of G-quadruplex structures. Nat Rev Genet 2012; 13:770–80.

8. Spiegel J, Adhikari S, Balasubramanian S. The Structure and Function of DNA G-Quadruplexes. Trends Chem 2020; 2:123–36.

9. Groth A, Rocha W, Verreault A, Almouzni G. Chromatin challenges during DNA replication and repair. Cell 2007; 128:721–33.

10. Li B, Carey M, Workman JL. The role of chromatin during transcription. Cell 2007; 128:707–19.

11. Robinson J, Raguseo F, Nuccio SP, Liano D, Di Antonio M. DNA G-quadruplex structures: more than simple roadblocks to transcription? Nucleic Acids Res 2021; 49:8419–31.

12. Datta S, Patel M, Patel D, Singh U. Distinct DNA Sequence Preference for Histone Occupancy in Primary and Transformed Cells. Cancer Inform 2019; 18:1176935119843835.

13. West AG, Fraser P. Remote control of gene transcription. Hum Mol Genet 2005; 14 Spec No 1:R101–11.

14. Hapgood JP, Riedemann J, Scherer SD. Regulation of gene expression by GC-rich DNA cis-elements. Cell Biol Int 2001; 25:17–31.

15. Hänsel-Hertsch R, Di Antonio M, Balasubramanian S. DNA G-quadruplexes in the human genome: detection, functions and therapeutic potential. Nat Rev Mol Cell Biol 2017; 18:279–84.

16. Nikolova EN, Kim E, Wise AA, O’Brien PJ, Andricioaei I, Al-Hashimi HM. Transient Hoogsteen base pairs in canonical duplex DNA. Nature 2011; 470:498–502.

17. Alvey HS, Gottardo FL, Nikolova EN, Al-Hashimi HM. Widespread transient Hoogsteen base pairs in canonical duplex DNA with variable energetics. Nat Commun 2014; 5:4786.

18. Rhodes D, Lipps HJ. G-quadruplexes and their regulatory roles in biology. Nucleic Acids Res 2015; 43:8627–37.

19. Kudlicki AS. G-Quadruplexes Involving Both Strands of Genomic DNA Are Highly Abundant and Colocalize with Functional Sites in the Human Genome. PLoS One 2016; 11:e0146174.

20. Murat P, Guilbaud G, Sale JE. DNA polymerase stalling at structured DNA constrains the expansion of short tandem repeats. Genome Biol 2020; 21:209.

21. Mendoza O, Bourdoncle A, Boulé J-B, Brosh RM Jr, Mergny J-L. G-quadruplexes and helicases. Nucleic Acids Res 2016; 44:1989–2006.

22. Frasson I, Pirota V, Richter SN, Doria F. Multimeric G-quadruplexes: A review on their biological roles and targeting. Int J Biol Macromol 2022; 204:89–102.

23. Varshney D, Spiegel J, Zyner K, Tannahill D, Balasubramanian S. The regulation and functions of DNA and RNA G-quadruplexes. Nat Rev Mol Cell Biol 2020; 21:459–74.

24. Gong J-Y, Wen C-J, Tang M-L, Duan R-F, Chen J-N, Zhang J-Y, Zheng K-W, He Y, Hao Y-H, Yu Q, et al. G-quadruplex structural variations in human genome associated with single-nucleotide variations and their impact on gene activity. Proc Natl Acad Sci U S A [Internet] 2021; 118. Available from: http://dx.doi.org/10.1073/pnas.2013230118

25. David AP, Margarit E, Domizi P, Banchio C, Armas P, Calcaterra NB. G-quadruplexes as novel cis-elements controlling transcription during embryonic development. Nucleic Acids Res 2016; 44:4163–73.

26. Lago S, Nadai M, Cernilogar FM, Kazerani M, Domíniguez Moreno H, Schotta G, Richter SN. Promoter G-quadruplexes and transcription factors cooperate to shape the cell type-specific transcriptome. Nat Commun 2021; 12:3885.

27. Zhang X, Spiegel J, Martínez Cuesta S, Adhikari S, Balasubramanian S. Chemical profiling of DNA G-quadruplex-interacting proteins in live cells. Nat Chem 2021; 13:626–33.

28. Brázda V, Hároníková L, Liao JCC, Fojta M. DNA and RNA quadruplex-binding proteins. Int J Mol Sci 2014; 15:17493–517.

29. Huang Z-L, Dai J, Luo W-H, Wang X-G, Tan J-H, Chen S-B, Huang Z-S. Identification of G-Quadruplex-Binding Protein from the Exploration of RGG Motif/G-Quadruplex Interactions. J Am Chem Soc 2018; 140:17945–55.

30. Williams P, Li L, Dong X, Wang Y. Identification of SLIRP as a G Quadruplex-Binding Protein. J Am Chem Soc 2017; 139:12426–9.

31. Ribeyre C, Lopes J, Boulé J-B, Piazza A, Guédin A, Zakian VA, Mergny J-L, Nicolas A. The yeast Pif1 helicase prevents genomic instability caused by G-quadruplex-forming CEB1 sequences in vivo. PLoS Genet 2009; 5:e1000475.

32. Yan KK-P, Obi I, Sabouri N. The RGG domain in the C-terminus of the DEAD box helicases Dbp2 and Ded1 is necessary for G-quadruplex destabilization. Nucleic Acids Res 2021; 49:8339–54.

33. Wallgren M, Mohammad JB, Yan K-P, Pourbozorgi-Langroudi P, Ebrahimi M, Sabouri N. G-rich telomeric and ribosomal DNA sequences from the fission yeast genome form stable G-quadruplex DNA structures in vitro and are unwound by the Pfh1 DNA helicase. Nucleic Acids Res 2016; 44:6213–31.

34. Ghosh M, Singh M. RGG-box in hnRNPA1 specifically recognizes the telomere G-quadruplex DNA and enhances the G-quadruplex unfolding ability of UP1 domain. Nucleic Acids Res 2018; 46:10246–61.

35. Hänsel-Hertsch R, Beraldi D, Lensing SV, Marsico G, Zyner K, Parry A, Di Antonio M, Pike J, Kimura H, Narita M, et al. G-quadruplex structures mark human regulatory chromatin. Nat Genet 2016; 48:1267–72.

36. Hänsel-Hertsch R, Spiegel J, Marsico G, Tannahill D, Balasubramanian S. Genomewide mapping of endogenous G-quadruplex DNA structures by chromatin immunoprecipitation and high-throughput sequencing. Nat Protoc 2018; 13:551–64.

37. Traganos F, Darzyndiewicz Z, Sharpless T, Melamed MR. Denaturation of deoxyribonucleic acid in situ effect of formaldehyde. J Histochem Cytochem 1975; 23:431–8.

38. Hoffman EA, Frey BL, Smith LM, Auble DT. Formaldehyde crosslinking: a tool for the study of chromatin complexes. J Biol Chem 2015; 290:26404–11.

39. Kuryavyi V, Patel DJ. Solution structure of a unique G-quadruplex scaffold adopted by a guanosine-rich human intronic sequence. Structure 2010; 18:73–82.

40. Lightfoot HL, Hagen T, Tatum NJ, Hall J. The diverse structural landscape of quadruplexes. FEBS Lett 2019; 593:2083–102.

41. Liu H, Wang R, Yu X, Shen F, Lan W, Haruehanroengra P, Yao Q, Zhang J, Chen Y, Li S, et al. High-resolution DNA quadruplex structure containing all the A-, G-, C-, T-tetrads. Nucleic Acids Res 2018; 46:11627–38.

42. Kazemier HG, Paeschke K, Lansdorp PM. Guanine quadruplex monoclonal antibody 1H6 cross-reacts with restrained thymidine-rich single stranded DNA. Nucleic Acids Res 2017; 45:5913–9.

43. Agarwal P, Enroth S, Teichmann M, Jernberg Wiklund H, Smit A, Westermark B, Singh U. Growth signals employ CGGBP1 to suppress transcription of Alu-SINEs. Cell Cycle 2016; 15:1558–71.

44. Agarwal P, Collier P, Fritz MH-Y, Benes V, Wiklund HJ, Westermark B, Singh U. CGGBP1 mitigates cytosine methylation at repetitive DNA sequences. BMC Genomics 2015; 16:390.

45. Patel D, Patel M, Westermark B, Singh U. Dynamic bimodal changes in CpG and non-CpG methylation genome-wide upon CGGBP1 loss-of-function. BMC Res Notes 2018; 11:419.

46. Lane AN, Chaires JB, Gray RD, Trent JO. Stability and kinetics of G-quadruplex structures. Nucleic Acids Res 2008; 36:5482–515.

47. Sathyaseelan C, Vijayakumar V, Rathinavelan T. CD-NuSS: A Web Server for the Automated Secondary Structural Characterization of the Nucleic Acids from Circular Dichroism Spectra Using Extreme Gradient Boosting Decision-Tree, Neural Network and Kohonen Algorithms. J Mol Biol 2021; 433:166629.

48. del Villar-Guerra R, del Villar-Guerra R, Trent JO, Chaires JB. G-Quadruplex Secondary Structure Obtained from Circular Dichroism Spectroscopy [Internet]. Angewandte Chemie 2018; 130:7289–93. Available from: http://dx.doi.org/10.1002/ange.201709184

49. Ajjugal Y, Kolimi N, Rathinavelan T. Secondary structural choice of DNA and RNA associated with CGG/CCG trinucleotide repeat expansion rationalizes the RNA misprocessing in FXTAS. Sci Rep 2021; 11:8163.

50. Zhou B, Geng Y, Liu C, Miao H, Ren Y, Xu N, Shi X, You Y, Lee T, Zhu G. Characterizations of distinct parallel and antiparallel G-quadruplexes formed by two-repeat ALS and FTD related GGGGCC sequence. Sci Rep 2018; 8:2366.

51. Singh U, Westermark B. CGGBP1--an indispensable protein with ubiquitous cytoprotective functions. Ups J Med Sci 2015; 120:219–32.

52. Müller-Hartmann H, Deissler H, Naumann F, Schmitz B, Schröer J, Doerfler W. The human 20-kDa 5’-(CGG)(n)-3’-binding protein is targeted to the nucleus and affects the activity of the FMR1 promoter. J Biol Chem 2000; 275:6447–52.

53. Identification of putative G-quadruplex DNA structures in S. pombe genome by quantitative PCR stop assay. DNA Repair 2019; 82:102678.

54. Hoffmann RF, Moshkin YM, Mouton S, Grzeschik NA, Kalicharan RD, Kuipers J, Wolters AHG, Nishida K, Romashchenko AV, Postberg J, et al. Guanine quadruplex structures localize to heterochromatin. Nucleic Acids Res 2017; 45:6253.

55. Patro LPP, Kumar A, Kolimi N, Rathinavelan T. 3D-NuS: A Web Server for Automated Modeling and Visualization of Non-Canonical 3-Dimensional Nucleic Acid Structures. J Mol Biol 2017; 429:2438–48.

